# Cell type-focused compound screen in human organoids reveals CK1 and MAPK11 inhibition protects cone photoreceptors from death

**DOI:** 10.1101/2023.10.09.561525

**Authors:** Stefan E. Spirig, Valeria J. Arteaga-Moreta, Zoltan Raics, Susana Posada-Céspedes, Stephanie Chreng, Olaf Galuba, Inga Galuba, Isabelle Claerr, Steffen Renner, Larissa Utz, P. Timo Kleindienst, Adrienn Volak, Jannick Imbach, Svitlana Malysheva, Rebecca A. Siwicki, Vincent Hahaut, Yanyan Hou, Simone Picelli, Marco Cattaneo, Josephine Jüttner, Cameron S. Cowan, Myriam Duckely, Daniel K. Baeschlin, Magdalena Renner, Vincent Unterreiner, Botond Roska

## Abstract

Human organoids that mirror their corresponding organs in cell-type diversity present an opportunity to perform large-scale screens for compounds that protect disease-affected or damage healthy cell types. However, such screens have not yet been performed. Here, we generated 20,000 human retinal organoids with GFP-labeled cone photoreceptors. Since degeneration of cones is a leading cause of blindness, we induced cone death and screened 2,707 compounds with known targets, for those that saved cones or those that further damaged cones. We identified inhibitors of CK1 or MAPK11 that protected cones, HSP90 inhibitors that saved cones in the short term but damaged them in the longer term, and broad HDAC inhibition by many compounds that significantly damaged cones. This work provides a database for cone-damaging compounds and describes compounds that can be starting points to develop neuroprotection for cones in diseases such as macular degeneration.

## INTRODUCTION

Human organoids are three-dimensional (3D) cellular assemblies grown from stem cells. They mimic the cellular architecture, cell-type diversity, and in some cases, the functionality of their corresponding organs^1,2^. A wide range of organoids have been generated that model organs such as the brain, retina, thyroid, heart, vasculature, lung, liver, gastrointestinal tract, pancreas, kidney, female reproductive tract, placenta, prostate, and testis^3–6^. These organoids, derived from both healthy and disease-affected individuals, serve multiple purposes such as unraveling the fundamental biology of organ development and deciphering the mechanisms of genetic diseases^2^.

Two other potential uses of human organoids are to screen for compounds that can improve or reverse a disease phenotype^7,8^ and, in toxicology, to screen for compounds with potential side effects on specific organs^9^. These two objectives rely on efficient and large-scale production of human organoids, as well as the comprehensive recording and analysis of phenotypic changes within 3D tissues. Noteworthy progress has been made in screening for compounds using mouse intestinal organoids^10^ and dissociated cells from mouse retinal organoids^11^, contributing to our understanding of organ biology and allowing for potential therapy development. However, therapies developed in mice do not always translate to humans due to variation in cell types and molecular pathways between the two species^12–14^. Screening of compounds in human organoids has only been done on a small scale, using fewer than sixty compounds and has been limited to cancer organoids^7,8^. Large-scale screening in human organoids for compounds that can alleviate disease phenotypes or cause harmful side effects, has not been described.

The brain, including the retina, is composed of numerous cell types^15–17^ and a fundamental characteristic of brain diseases is ‘selective vulnerability’, where the pathology predominantly affects specific cell types^18^. For instance, in Parkinson’s disease, the motor symptoms are primarily caused by targeted loss of dopaminergic neurons in the ventral substantia nigra pars compacta^19^. Similarly, in Huntington’s disease, medium spiny GABAergic neurons in the striatum are particularly affected^20^. Given the selective vulnerability of specific cell types in various brain diseases, compound screening to find drugs that mitigate disease phenotypes in brain or retinal organoids will be most effective when they focus specifically on the cell types affected by the disease.

The human retina is part of the brain^21^ and contains diverse cell types arranged in five layers^22–24^. Photoreceptors sense light and transmit information to about ten different types of bipolar cells. In turn, the bipolar cells further transmit visual information to an even greater variety of ganglion cells. Ganglion cells are the output neurons of the retina, with their axons forming the optic nerve through which information is broadcast to the rest of the brain. The transmission of information from photoreceptors to bipolar cells is influenced by horizontal cells, while transmission from bipolar cells to ganglion cells is modulated by a multitude of amacrine cell types^21^.

Photoreceptors in the retina are of two types: rods and cones. Rods are utilized in low-light conditions, such as at dusk, and lack of rod function results only in mild or no vision impairment^25^. Cones are primarily responsible for image formation in daylight and enable the high-resolution vision that allows reading and face recognition. The loss of cones or their functionality, mainly due to age-related macular degeneration or end-stage retinitis pigmentosa, results in blindness and affects over 200 million individuals worldwide^26,27^. In age-related macular degeneration, the dysfunction or death of cones is either a direct consequence of the disease or secondary to the dysfunction of the retinal pigment epithelium^28^. Retinitis pigmentosa, a group of monogenic retinal diseases, primarily affects rods^29,30^ and cones degenerate as a secondary consequence of rod death^31^. The reasons behind cone degeneration in age-related macular degeneration and retinitis pigmentosa are extensively studied but are not fully understood, and several approaches are being developed to halt the degeneration process^32–49^. A common theme for retinitis pigmentosa has emerged that cones are likely starving from lack of glucose^32,38,50–52^. Slowing down cone degeneration in patients has so far not been achieved. Given the critical importance of cones for human vision, preserving their viability remains a significant objective in medicine.

Retinal organoids were among the earliest organoids to be established^53,54^. Since then, the technology for developing human retinal organoids has advanced rapidly and allows for the generation of complex, five-layered organoids that are light sensitive and consist of multiple cell types resembling those found in the adult human retina^55,22,56^. Cones in human retinal organoids exhibit a close similarity to their counterparts in the adult human retina in terms of gene expression, morphology, and function^22,56^ and, thus, present a unique opportunity to study cone degeneration.

Here, we aimed to find compounds that either slow down or induce the death of cones. Compounds that can preserve cones in organoids may pave the way for therapies that mitigate cone loss in conditions such as macular degeneration and retinitis pigmentosa. Compounds that induce cone death and their targets can be valuable for assessing the safety of new compounds planned for clinical trials. We produced ∼20,000 human retinal organoids each 30 weeks old, which is a time when organoid cone transcriptomes are similar to those of adult human retina^22^. We targeted GFP expression specifically to cones using adeno associated viral vectors (AAVs) carrying a cone-specific promoter^14^ and then induced cone death by glucose starvation. We conducted 3D imaging of the organoids before and after the starvation process, at a time when approximately 40% of cones were lost. We then evaluated a library of 2,707 compounds with known targets^57^. This screening identified compounds that exacerbated cone death as well as compounds that counteracted the degeneration induced by glucose starvation. Analysis of the targets of the compounds revealed that broad inhibition of class I or II histone deacetylases (HDACs I/II) by various compounds resulted in significant damage to cone photoreceptors. Heat shock protein 90 (HSP90) inhibition countered cone degeneration for a few days but proved detrimental after a week. We also discovered two kinase inhibitors that consistently preserved cones over a prolonged period; both compounds also preserved rods. Through kinase profiling of both compounds and their inactive chemical analogues, we identified two of their targets: CSNK1G1 and MAPK11. Inhibiting casein kinase 1 (CK1, CSNK1G is a member of CK1 family) with each of three different compounds resulted in increased survival of both cones and rods. Additionally, two compounds that inhibit MAPK11 enhanced cone survival, one of which also improved rod survival. Taken together, we describe here a technology to perform cell type-focused screening in human organoids, and a publicly available resource of 2,707 compounds and their targets, together with their positive or negative effects on human cone survival (https://ConeTargetedCompoundScreen.iob.ch). Furthermore, we identify kinase inhibitors that could be starting points to develop medicine that counters cone and/or rod degeneration.

## RESULTS

### Specific and rapid live-labeling of cones in human retinal organoids

We modified the Agarose Multiwell Array Seeding and Scraping (AMASS) method^22^ to generate ∼20,000 five-layered human retinal organoids that were grown for 30 weeks (Figures 1A and 1B). These organoids contain the major cell classes and various cell types found in the human retina^22^ (Figure S1). They also exhibit functional cone photoreceptors with light-sensitive outer segments^22,56^, inner segments, cell bodies, and axons. Moreover, the transcriptomic profile of the cones in the organoids resembles that of adult human cones^22^.

**Figure 1:**
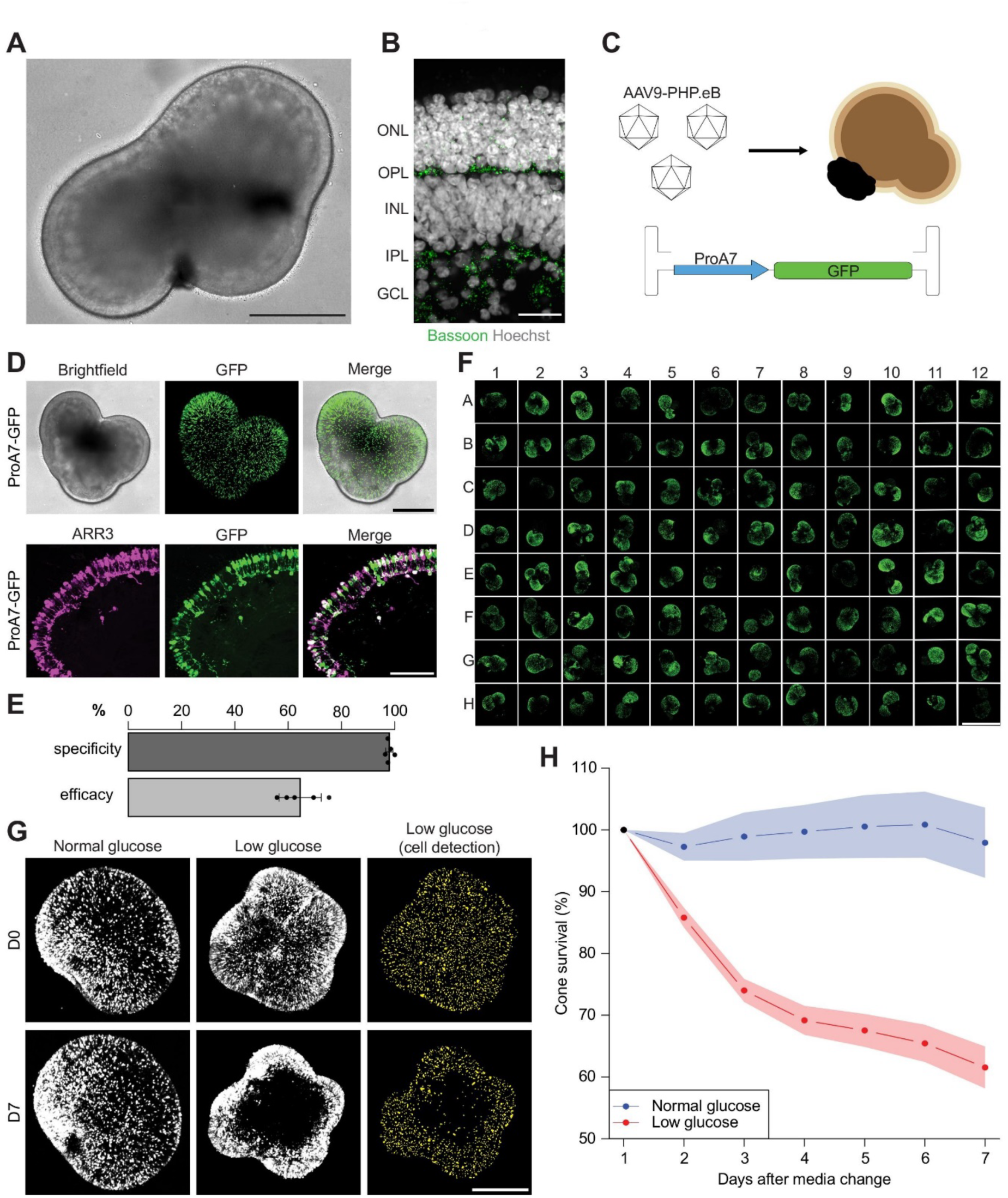
Glucose starvation induces rapid death of cones in human retinal organoids. **A:** Brightfield live image of a human retinal organoid (scale bar, 500 µm). **B:** Confocal image of a sectioned and stained human retinal organoid (scale bar, 25 µm). Bassoon, green; Hoechst, white. ONL: outer nuclear layer, OPL: outer plexiform layer, INL: inner nuclear layer, IPL: inner plexiform layer, GCL: ganglion cell layer. **C:** Schematic illustrating the transduction strategy for organoids. **D:** Top: live images of a ProA7-GFP AAV-transduced human retinal organoid (scale bar, 500 µm). Bottom: confocal image of a sectioned and stained ProA7-GFP AAV-transduced human retinal organoid (scale bar, 100 µm). ARR3, magenta; GFP, green. **E:** Quantification of the specificity and efficacy of cone labeling by ProA7-GFP AAV. Results are shown as mean ± sd. **F:** Representative image of a 96-well plate containing ProA7-GFP AAV-transduced human retinal organoids (scale bar, 2 mm). **G:** Left: representative live images of human retinal organoids at day 0 (D0) and day 7 (D7) in either normal or low glucose medium. Right: detected cones in organoids in low glucose. GFP, white. Detected cells, yellow. **H**: Quantification of cone survival in human retinal organoid in both normal and low glucose conditions over seven days. The quantification was performed using the 3D-additive-count algorithm. The results are shown as mean ± se.

To visualize living cones, we transduced organoids with AAVs expressing GFP under control of the cone-specific promoter ProA7^14^ (Figures 1C and 1D). We tested five different capsid variants and found that AAVs with the AAV9-PHP.eB capsid^58^ provided the highest number of labeled cones within four weeks. The cone labeling specificity in organoids was 97±1% (n=5), while the efficacy was 64±8% (n=5) (Figure 1E). We performed AAV transduction in cell culture flasks that allowed simultaneous labelling of ∼130 organoids, which were subsequently positioned into 96-well plates for 3D imaging (Figure 1F). As a result, we achieved GFP labeling of cones in a large number of organoids with high specificity, efficacy and uniformity.

### Glucose starvation induces cone death

To induce cone death, we cultured organoids individually in 96-well plates using low glucose medium^59^. We recorded cell death by monitoring the disappearance of cytosolic GFP as a reliable and rapid indicator of cell death^60–62^. This was realized by 3D confocal imaging of GFP in each well of the 96-well plate. The organoids were imaged for two weeks without a change of the medium to avoid any displacement of the organoids. Additionally, we included organoids in normal glucose medium in the same 96-well plate as controls (Figure 1G).

We developed three different cell-counting algorithms to determine the number of cones in each organoid (Figure 1G). The first (referred to as ’3D-additive-count’) quantifies cones image-by-image from a 3D confocal image stack. The second (’3D-count’) counts cones from the entire 3D stack. The third (’MIP-count’) counts cones from the maximum intensity projection of the 3D stack. While the primary measure of cone numbers was based on the 3D-additive-count, we also verified the results using the 3D-count and MIP-count methods.

We measured glucose concentration in the low and normal glucose media during organoid imaging without exchanging the media. Initially, glucose levels in the low glucose medium were 9% of those in the normal glucose medium. With time, glucose levels decreased in both the low and the normal glucose media, becoming undetectable after two days in the low glucose medium and seven days in the normal glucose medium (Figure S1).

Next, we examined number of cones over the two-week period in both types of media. In the low glucose medium, the organoids began losing cones after two days of starvation, and this cone loss continued over the two weeks. Conversely, cone numbers in the control organoids in normal glucose medium remained constant until day seven, after which they declined (Figures 1G, 1H and S1). Therefore, we conducted further experiments for a duration of seven days. Within this time frame, the glucose-deprived organoids lost ∼40% of their cones, leading to a significant difference in cone numbers in the low and normal glucose conditions (low glucose, n=26; normal glucose, n=6, p<0.001, Mann–Whitney U test, Figure 1H). Cone cell death occurred predominantly in the central region of the organoid rather than at the edges (Figure 1G). This spatial variation is likely attributable to the additional stress experienced by cells in contact with the bottom of the well. The finding that only about half of the cones were lost by the end of seven days in low glucose medium enabled us to screen for compounds that either slow down cone death or accelerate it.

We then performed two compound screens. In the primary screen, we tested all compounds at the same concentration of 10 µM. In the secondary screen, we selected compounds based on the results of the primary screen and retested them at multiple concentrations.

### Primary compound screen

In the primary screen, we used ∼15,000 organoids distributed across 175 separate 96-well plates. Each 96-well plate included eight control wells, consistently positioned. Four of these control wells contained organoids in normal glucose medium with dimethyl sulfoxide (DMSO, 0.1%) but no compounds. The other four contained organoids in low glucose medium (DMSO, 0.1%) without compounds. Each remaining individual well (88 wells) within a 96-well plate contained a unique compound, and each compound was present in five different plates, resulting in five replicates of each compound in the screen (Figure 2A). All compounds were stored in DMSO as the solvent. Although the compounds were stored in 384-well plates, we conducted the screening in 96-well plates because the human retinal organoids were too large to be cultured in 384-well plates.

**Figure 2:**
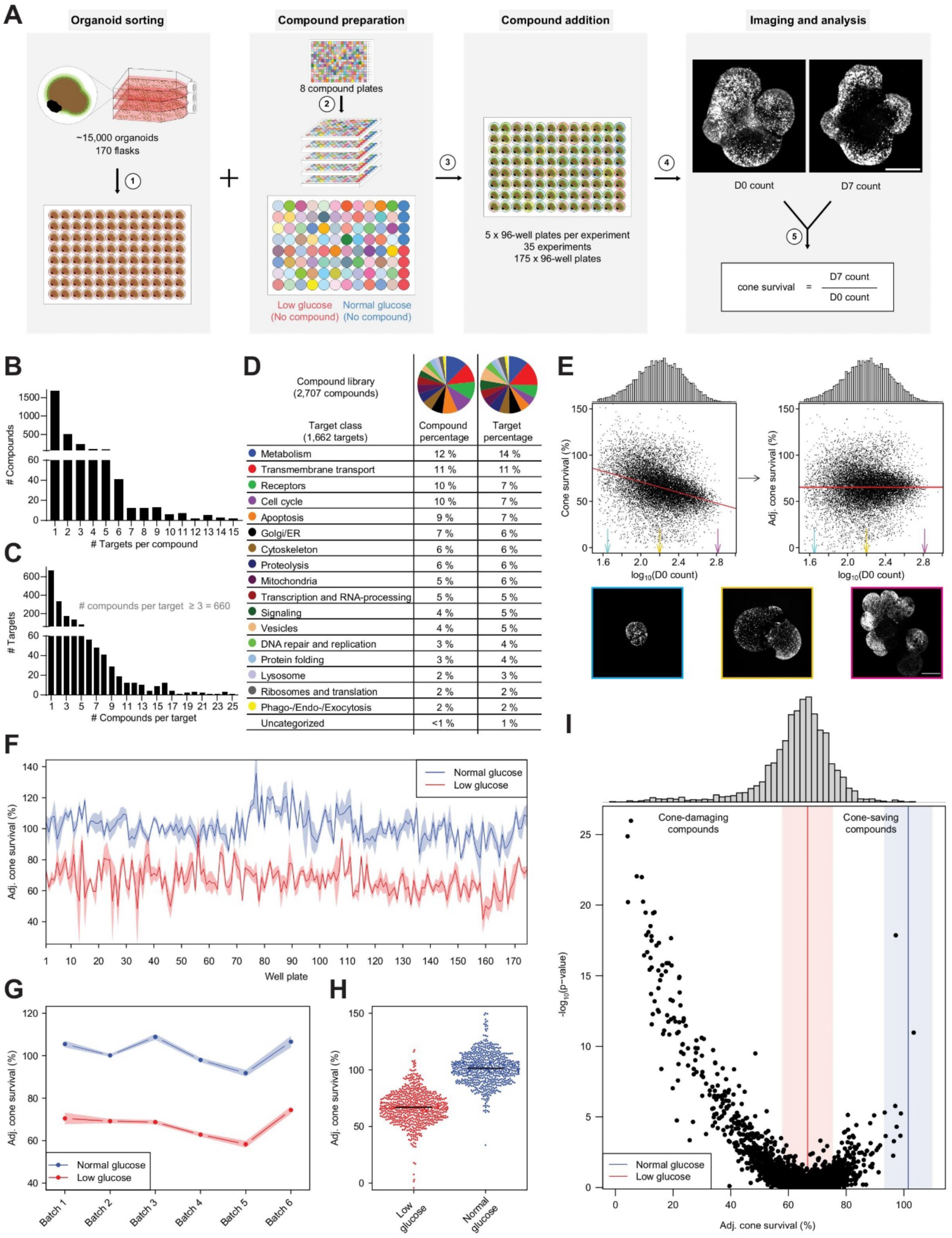
Primary screen of compounds that damage or save cones. **A:** Schematic illustration of the primary screen. 1. Transfer of human retinal organoids from cell culture flasks to 96-well plates. 2. Compound distribution from 384- to 96-well plates with five replicates of each compound. Low glucose controls, red. Normal glucose controls, blue. 3. Addition of compounds to human retinal organoids. 4. Human retinal organoid imaging. 5. Quantification of cones from the same human retinal organoid at D0 and D7. GFP, white. **B**: Bar chart showing the number of compounds with a given number of targets. **C:** Bar chart showing the number of targets with a given number of compounds per target. **D:** Categorization of targets and compounds of the compound library. **E:** Dependence of cone survival on the D0 cone count. Top left: cone survival as a function of the logarithm of the cone count at D0, with a fitted linear model; the regression line is shown in red. The distribution of D0 counts is shown above. Top right: adjusted cone survival as a function of the logarithm of the cone count at D0 with the transformed regression line in red. The distribution of D0 counts is shown above. Bottom: example images with different D0 cone counts (scale bar, 500 µm). Colored arrows and frames around images indicate corresponding D0 cone counts. GFP, white. **F-G**: Adjusted cone survival of normal and low glucose controls for individual well plates (F) and organoid batches (G) using the 3D-additive-count algorithm, mean ± se. **H:** Adjusted cone survival of all normal and low glucose control human retinal organoids. **I:** Effect of compounds on cone survival in the primary screen. Each dot corresponds to the effect of one compound, with the median of the adjusted cone survival of five human retinal organoids on the x-axis and the p-value comparing the cone survival between the compound and the low glucose controls on the y-axis. The median (line) and the interquartile range (shaded area) of normal (blue) and low (red) glucose controls are indicated. Top: the distribution of median adjusted cone survival for compounds. Results were obtained using the 3D-additive-count algorithm.

In preparation for the screen, we transferred compounds from a 384-well plate to their designated locations on 96-well plates and then dissolved them in low glucose medium. To initiate the screen, we first moved organoids from flasks containing normal glucose medium to 96-well plates and washed them there in low glucose medium. We then removed the low glucose medium from the 96-well plates containing the organoids and added the compounds dissolved in low glucose medium. We conducted a 3D confocal scan of organoids with GFP-labeled cones in each well of the 96-well plates at the beginning of the screen and again at the end, seven days later. We then used the three algorithms to quantify the number of cones at day zero (D0) and day seven (D7), defining the ratio of counts at D7 and D0 as the measure of cone survival (Figure 2A).

We screened a compound library of 2,707 compounds for their impact on cone survival upon glucose deprivation. The compound library, called ‘Mode of Action library’ (MOA library) contained compounds with known protein targets^57^. Some compounds had multiple targets, while others had only one (Figure 2B). Similarly, certain targets were affected by multiple compounds, while others were affected by only one compound (Figure 2C). The MOA library had a total of 1,662 targets that are involved in a wide range of biological processes (Figure 2D).

To exclude accidentally empty wells or organoids with little or no labeled cones, we applied a threshold on the D0 cone count and removed wells with lower D0 counts from the dataset (Figure S2). This was necessary since the ratio of D7 and D0 counts is sensitive to low D0 counts. Furthermore, we observed that the survival of cones was determined not only by the compounds but also by the initial cone count. There was a log-linear relationship between the D0 count and cone survival at D7 in low glucose (n=14,529, R^2^=0.12, p<0.001, 3D-additive-count) with all three cell-counting algorithms (Figures 2E and S2). To account for this relationship, we adjusted the values of cone survival using a linear transformation yielding the quantity ‘adjusted cone survival’. Adjusted cone survival therefore does not depend on the cone count at D0.

The locations of organoids on the 96-well plates did not influence cone survival: the mean adjusted survival at all positions (8.8% maximum difference between positions) remained within one standard deviation of that of the position exhibiting the least variation (12.9%), excluding the normal glucose well positions (Figure S2).

To investigate additional potential batch effects in the screen, we examined the distribution of the mean adjusted cone survival of the eight control organoids: four in normal and four in low glucose; first across the 175 separate 96-well plates (Figures 2F and S2), second across the 35 experiments, each consisting of five 96-well plates with the same compound (Figure S2), and third, across the six independent organoid productions used for the primary screen (Figures 2G and S2). We found no major differences between batches at any level. The mean adjusted cone survival of normal and low glucose control organoids was significantly different across all 35 experiments and across all six organoid productions, using all three cell-counting algorithms (p-values<0.001, Mann– Whitney U test, Figures 2G-H and S2). Cone survival values showed strong and significant correlations across the three cell counting algorithms (R=0.92-0.97, p-values<0.001, Pearson correlation, Figure S3), suggesting that the quantification of cone survival is robust across the different algorithms.

After accounting for potential confounding variables and batch effects, we proceeded to analyze the outcome of the primary screen. The mean adjusted cone survival of most compounds was not significantly different from the mean adjusted cone survival of the low glucose control. However, one set of compounds significantly impacted cone survival negatively (‘cone-damaging compounds’, p-values<0.05, ANCOVA with Benjamini Hochberg correction for multiple testing), while another set of compounds showed a beneficial effect on cone survival (‘cone-saving compounds’) (Figures 2I and S3).

### Secondary screen: cone-damaging compounds

In order to validate the cone-damaging and cone-saving compounds detected in the primary screen and to assess the concentration dependence of their actions, we proceeded to secondary screens.

First, we revisited the 33 most damaging compounds selected from a set of 146 compounds that caused significant damage beyond that induced by the low glucose medium in the primary screen (n=5 for each compound, p-values<0.05, ANCOVA with Benjamini Hochberg correction for multiple testing, Table S1). We investigated these compounds at four concentrations (0.01, 0.1, 1, 10 µM) in five replicates using normal glucose medium. We used normal glucose medium to confirm the damaging effects of these compounds on cones of healthy organoids. Furthermore, each 96-well plate included four wells with only DMSO in normal glucose medium, serving as a negative control (Figure 3A). As for the positive control, we utilized Staurosporine, a nonselective ATP-competitive kinase inhibitor known for inducing apoptosis^63,64^. This was tested also in five replicates at each of the four concentrations.

**Figure 3:**
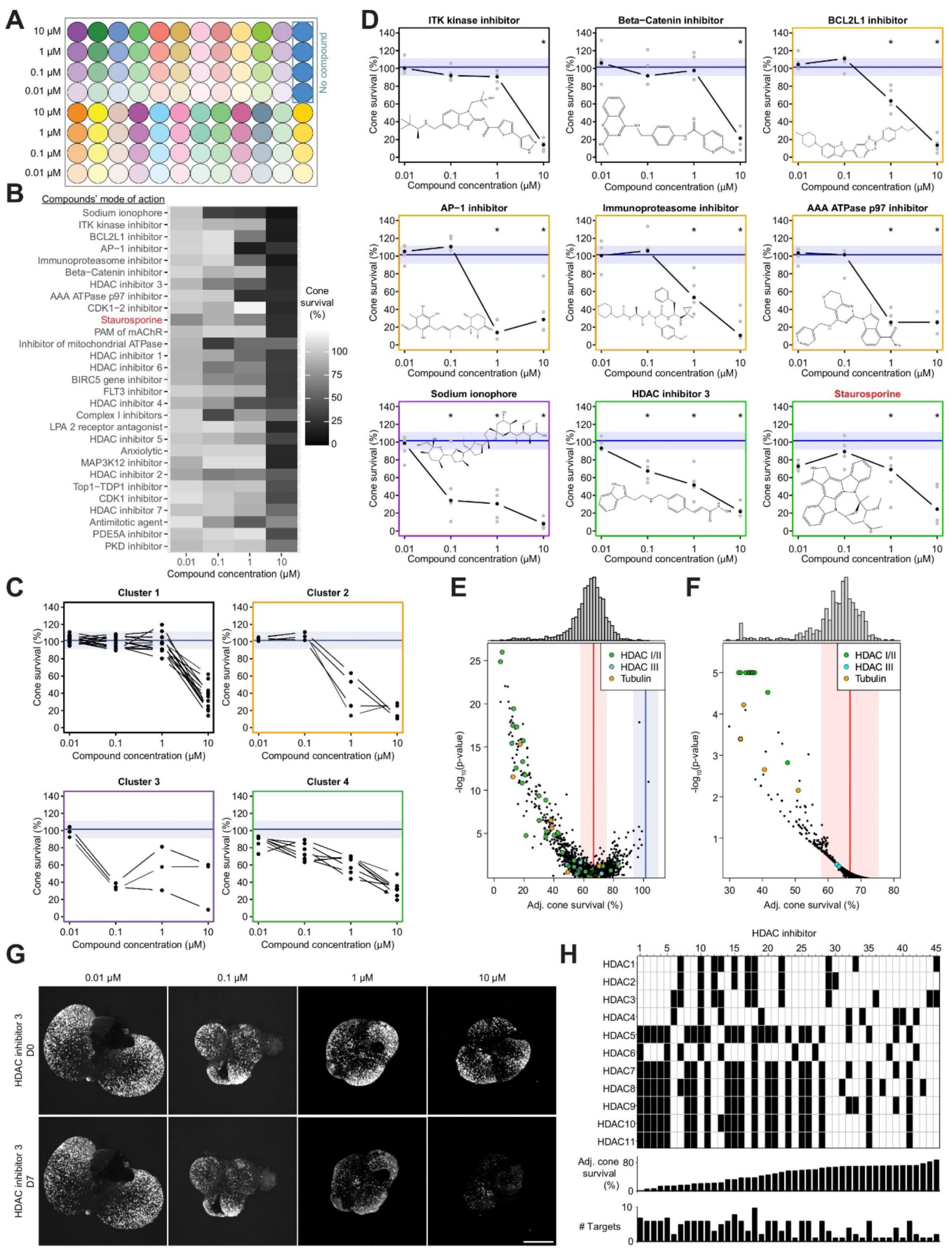
Secondary screen of cone-damaging compounds. **A:** Schematic illustration of the plate layout in the secondary screen for cone-damaging compounds. Different colors represent distinct compounds. No compound control, blue. **B:** Summary of compound effects for significant cone-damaging compounds (p<0.05 for at least one concentration after Benjamini Hochberg correction for multiple testing). The positive control Staurosporine is indicated in red. Compounds are ordered by minimum p-value per concentration, with the smallest p-value at the top. **C:** Dose-responses for significant compounds within the same cluster. Dots represent the median of cone survival at the indicated concentrations. The median (line) and the interquartile range (shaded area) of normal (blue) glucose controls are indicated. **D:** Dose-response curves of eight compounds with the lowest p-values (at any concentration) and Staurosporine (red) with their corresponding cluster indicated. The median and the interquartile range are labeled as in (C). The colored frame indicates the corresponding cluster from (C). The structure of each compound is shown in the plot. Significant compounds and concentrations are marked with *, denoting a p-value<0.05 (after Benjamini Hochberg correction for multiple testing). **E**: Adjusted cone survival and p-values in the primary screen. HDAC I/II (green), HDAC III (cyan), and tubulin (orange) inhibitors are labeled. The median (line) and the interquartile range (shaded area) of normal (blue) and low (red) glucose controls are indicated. **F:** Target analysis of primary screen. The means of the median adjusted cone survival for each target are shown, along with their p-values. Compound targets are labeled. HDAC I/II, green; HDAC III, cyan; tubulin, orange. The median (line) and the interquartile range (shaded area) of low (red) glucose controls are indicated. **G**: Example images of human retinal organoids before (D0) and after (D7) treatment with different concentrations of HDAC inhibitor 3 (scale bar, 500 µm). GFP, white. **H**: Top: different HDAC I/II inhibitors (numbered from 1 to 45) with their targets (black shaded areas). Compounds are sorted from left to right based on median adjusted cone survival. Middle: Adjusted cone survival for each of the 45 compounds. Bottom: Number of HDAC I/II targets of the 45 compounds.

Most compounds (29 out of 33) again induced a significant decrease in the number of cones compared to the negative control (n=5 for each compound and concentration, minimum p-value<0.05, ANOVA with Benjamini Hochberg correction for multiple testing, Figure 3B). Of these compounds, nine caused a more significant decrease in cone numbers than the positive control Staurosporine, which reduced cone numbers by 76%. The remaining compounds reduced cone numbers by at least 38% (Figures 3B, S4 and S5, Table S2).

We observed a variety of dose-response curves for cone-damaging compounds used in the secondary screen that could be clustered into four groups. The first cluster included compounds that led to cone death only at the highest concentration (10 µM), such as an ITK kinase inhibitor and a Beta catenin inhibitor. The second cluster of curves had an action threshold of 1 µM and included a BCL2L1 inhibitor, an AP-1 inhibitor, an immunoproteasome inhibitor, and an AAA ATPase p97 inhibitor. The third cluster of curves had an action threshold of 0.1 µM, for example a sodium ionophore had such a curve. For the fourth cluster of curves, cone death increased linearly with the logarithm of compound concentration. Most of these curves belonged to HDAC I/II inhibitors (Figures 3C, 3D and S5).

Of the 146 compounds that caused significant damage to cones in the primary screen (p-values<0.05, ANCOVA with Benjamini Hochberg correction for multiple testing), 19 were identified as HDAC I/II inhibitors (Figure 3E and Table S1). Furthermore, in the secondary screen, seven of the 28 compounds that significantly damaged cones were HDAC I/II inhibitors (Figures 3B and S4). To determine whether this high number of HDAC I/II inhibitors is a result of bias in the MOA library towards HDAC I/II inhibitors or whether HDAC I/II inhibitors tend to damage cones more frequently, we reanalyzed the results of the primary screen. We calculated the probability that the mean adjusted cone survival for a randomly selected set of 45 compounds is lower than the mean adjusted cone survival for the 45 different HDAC I/II inhibitors in the MOA library. This probability was smaller than 0.0001, implying that, on average, the 45 HDAC I/II inhibitors in the MOA library cause more damage to cones than a random selection of the same number of other compounds.

To systematically investigate whether modulating any specific target is significantly more harmful to cones than modulating other targets, we first selected targets that had a minimum of three compounds listed in the MOA library. We identified a total of 660 such targets (Figure 2C). Next, we calculated the probability of observing a lower mean adjusted cone survival when randomly selecting the same number of compounds, in comparison to the compounds associated with the target. We identified 10 targets with a p-value of less than 0.0001 and less than 0.05 after Benjamini Hochberg correction for multiple testing. These targets belonged exclusively to class I/II HDACs (Figure 3F). Furthermore, when evaluating the percentage of compounds that led to significant cone damage among all compounds targeting each specific target, HDACs had the highest values (Figure S4). Each HDAC I/II target was associated with 9-29 distinct compounds, out of which 44-67% induced significant damage to cones. Interestingly, compounds that acted on Sirtuins, which are class III HDACs, had no influence on cone survival (Figures 3E, 3F and S4). HDAC I/II inhibitors often affect multiple HDAC I/II targets and we found a significant negative correlation between the number of HDAC I/II targets of a specific inhibitor and adjusted cone survival (p<0.001, R=-0.6, Spearman correlation). Therefore, cones are specifically sensitive to HDAC I/II inhibition, and HDAC I/II inhibitors with a wide range of targets are more likely to result in cone damage than those that are more selective (Figures 3G and 3H).

Cones were also highly sensitive to the inhibition of tubulins (Figures 3E, 3F and S4). The probability that the mean adjusted cone survival for a random set of 10 compounds was lower than that of the 10 distinct tubulin inhibitors in the MOA library was 0.002. Of the 10 tubulin-targeting compounds, four resulted in a significant decrease in cone numbers. Notably, three of these compounds had a broad target range, affecting 19-20 different tubulins.

### Secondary screen: cone-saving compounds

Seven compounds in the primary screen significantly increased cone survival. We proceeded with these compounds as well as 24 other compounds that had the highest effect in protecting cones during glucose deprivation in the primary screen. These 31 compounds were tested at four different concentrations (0.01, 0.1, 1, and 10 µM) in five replicates in low glucose medium. Each 96-well plate included four wells containing only DMSO in normal glucose medium as a positive control, and four wells containing only DMSO in low glucose medium as a negative control (Figure 4A).

**Figure 4:**
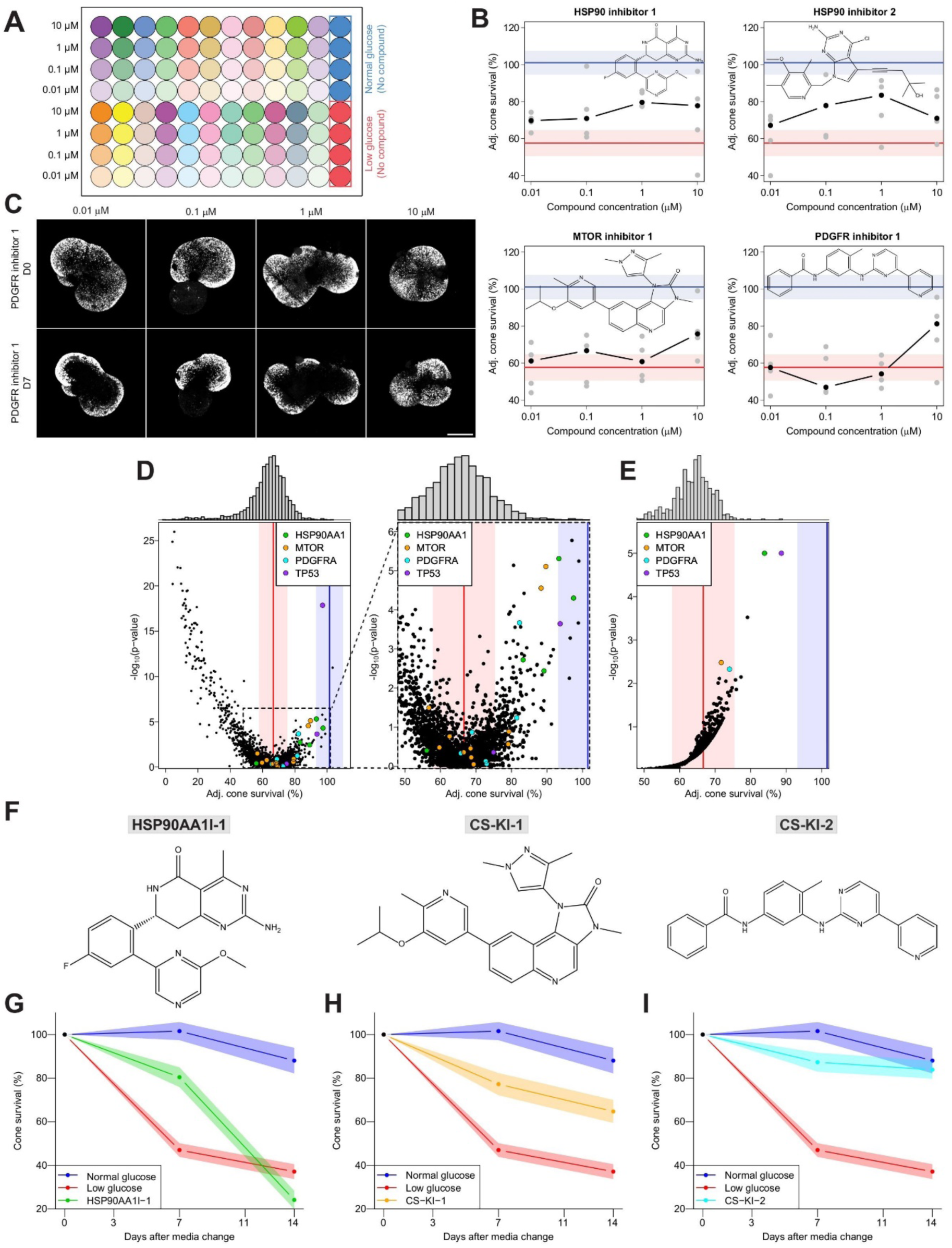
Secondary screen of cone-saving compounds. **A:** Schematic illustration of plate layout in the secondary screen for cone-saving compounds. Different colors indicate distinct compounds. Low glucose controls, red. Normal glucose controls, blue. **B:** Dose-response curves of the four significant compounds. Black dots, median adjusted cone survival; gray dots, individual adjusted cone survival values. The median (line) and the interquartile range (shaded area) of normal (blue) and low (red) glucose controls are indicated. The structure of each compound is shown in the plot and the annotated target is indicated on top. **C:** Example images of human retinal organoids at D0 and D7 with different concentrations of PDGFR inhibitor 1 (scale bar, 500 µm). GFP, white. **D:** Left: adjusted cone survival and p-values in the primary screen. Compound targets are labeled. HSP90AA1, green; MTOR, orange; PDGFRA, cyan; TP53, purple. The median and the interquartile range are labeled as in (B). Right: Zoom-in of plot on the left. **E**: Target analysis of the primary screen. The means of the median adjusted cone survivals for each target are shown, along with their p-values. HSP90AA1, green; MTOR, orange; PDGFRA, cyan; TP53, purple. The median and the interquartile range are labeled as in (B). **F:** Chemical structures and new names of the indicated compounds. **G-I**: Time course of cone survival in glucose-starved human retinal organoids with the indicated compounds for 14 days. Results are shown as mean ± se.

We confirmed that four out of the 31 compounds tested in the secondary screen have a significant positive impact on cone survival after correction for multiple testing (n=5 for each compound and concentration, minimum p-values<0.05, ANCOVA with Benjamini Hochberg correction for multiple testing, Figures 4B, 4C, S6, S7 and Table S3). These four compounds include an inhibitor of HSP90AA1 (‘HSP90 inhibitor 1’), an inhibitor of both HSP90AA1 and HSP90AB1 (‘HSP90 inhibitor 2’), an inhibitor of MTOR, PIK3CA, PIK3CB, and PIK3CD (‘MTOR inhibitor 1’), and an inhibitor of PDGFRA and PDGFRB (‘PDGFR inhibitor 1’). The two HSP90 inhibitors demonstrated stronger activity at lower concentrations (0.1 or 1 µM) and less activity at a concentration of 10 µM (Figure 4B). The dose-response curves of the MTOR inhibitor and the PDGFR inhibitor showed similar patterns, being effective only at the highest concentration of 10 µM (Figures 4B and 4C).

To understand the relationship between the four compounds and their targets in the context of cone protection, we re-analyzed the results of the primary screen, focusing on the targets of the four inhibitors that were confirmed to protect cones (HSP90AA1, HSP90AB1, MTOR, PIK3CA, PIK3CB, PIK3CD, PDGFRA, and PDGFRB). We found five compounds in the MOA library that target HSP90AA1, four of which resulted in adjusted cone survival higher than 80% (although some of these were not significant in the primary screen). For all other targets, the percentages of compounds with a positive effect on cone survival were only between 11 and 33% (Figures 4D and S8). These results suggest that HSP90AA1 is the target of HSP90 inhibitors 1 and 2 in cones. Additionally, they suggest that the other two compounds (MTOR inhibitor 1 and PDGFR inhibitor 1), which are both kinase inhibitors, have different causal targets in human cones than those originally listed in the MOA library.

To systematically examine whether modulating particular targets has a greater positive impact on cones compared to other targets, we conducted an analysis focusing on 660 targets that had at least three partner compounds listed in the MOA library. To rank targets in their ability to protect cones, we calculated the probability of observing a higher mean adjusted cone survival in the primary screen when randomly selecting the same number of compounds in comparison to the compounds associated with the target. HSP90AA1 emerged as one of the top-ranked targets (p<0.00001 and p<0.01 after Benjamini Hochberg correction for multiple testing). Targets such as HSP90AB1, MTOR, PIK3CA, PIK3CB, PIK3CD, PDGFRA, and PDGFRB were not significant because a large proportion of compounds acting on these targets had no impact on cone survival (Figures 4D, 4E and S8). TP53 was a further significant target (p<0.00001 and p<0.01 after Benjamini Hochberg correction for multiple testing), for which there were three compounds in the library, two inhibitors and one expression enhancer. However, both inhibitors and the expression enhancer increased cone survival in the primary screen, and the enhancer had no significant effect after retesting, suggesting that TP53 is not a target for cone protection.

To further explore the connection between the four validated compounds that protect cones and their targets, we compared available data on IC50 (the concentration at which a compound shows 50% of its maximum inhibitory effect) with the effect on cone survival of inhibitors of HSP90AA1, HSP90AB1, MTOR, PIK3CA, PIK3CB, PIK3CD, PDGFRA, and PDGFRB. We found a significant negative correlation between the reported IC50s and the adjusted cone survival for the inhibitors of HSP90AA1 (R=-0.7, Spearman correlation, p=0.03, Figure S8). This suggests that compounds with lower IC50 values (indicating greater potency) are linked to higher adjusted cone survival. However, the IC50s for the inhibitors of HSP90AB1, MTOR, PIK3CA, PIK3CB, PIK3CD, PDGFRA, and PDGFRB were not significantly correlated with cone survival. These findings further confirm that HSP90AA1 is a target that, when inhibited, effectively protects cones from the effects of glucose starvation at day seven. Therefore, we have renamed HSP90 inhibitor 1 as ‘HSP90AA1I-1’ and HSP90 inhibitor 2 as ‘HSP90AA1I-2’. On the other hand, HSP90AB1, MTOR, PIK3CA, PIK3CB, PIK3CD, PDGFRA, and PDGFRB are not causal targets that affect organoid cone survival. Since MTOR inhibitor 1 and PDGFR inhibitor 1 are kinase inhibitors, we renamed them as ‘Cone-Saving Kinase inhibitor 1’ (CS-KI-1), and as ‘Cone-Saving Kinase inhibitor 2’ (CS-KI-2), respectively (Figure 4F).

We then tested the effects of HSP90AA1I-1, CS-KI-1, and CS-KI-2 on organoids under normal glucose conditions. None of the compounds had a significant effect after seven days compared to the normal glucose control conditions (Figure S9).

We conducted additional tests on the three compounds HSP90AA1I-1, CS-KI-1, and CS-KI-2 over a 14-day period. We administered these compounds in the concentrations that had the strongest effect during the secondary screen: 1 µM for HSP90AA1I-1 and 10 µM for CS-KI-1 and CS-KI-2 (Figure 4B). We treated organoids with these compounds in low glucose medium and performed imaging after 7 and 14 days, using normal and low glucose media with DMSO (without the compounds) as negative and positive controls, respectively (Figures 4G-I).

HSP90AA1I-1 showed significant protection of cones from death after seven days but became detrimental to cones after 14 days compared to the low glucose control (n=10, 7 days: p<0.001, 14 days: p=0.01, Mann–Whitney U test, Figure 4G). In contrast, CS-KI-1 and CS-KI-2 had a significant rescuing effect on cones at both seven and 14 days (low glucose, n=10; CS-KI-1, n=9, 7 days: p<0.001, 14 days: p=0.002; CS-KI-2, n=10, 7 days: p<0.001, 14 days: p<0.001, Mann– Whitney U test, Figures 4H and 4I). These findings suggest that HSP90AA1 inhibition is detrimental to cone survival in the long term and that the protective effect of CS-KI-1 and CS-KI-2 on cones is of long duration.

### Effect of HSP90AA1I-1, CS-KI-1, and CS-KI-2 on rod photoreceptor death

HSP90AA1I-1, CS-KI-1, and CS-KI-2 each had a protective effect on cones after seven days of glucose starvation and we examined whether they would offer a similar protection to rods. To do this, we developed the promoter ProA330, which targeted rods in human retinal organoids. When we introduced AAV9-PHP.eB capsid-coated AAVs that expressed GFP under the control of the ProA330 into human retinal organoids, we observed GFP expression in rods with a specificity of 98 ± 3% (n=6) and an efficacy of 38 ± 3% (n=6) (Figures 5A and 5B). ProA330 also drove specific expression in rods of mouse retina in vivo with a specificity of 99.7% ± 0.5% and an efficacy of 41 ± 7% (n=3 retinas, 2 mice, Figure S10).

**Figure 5:**
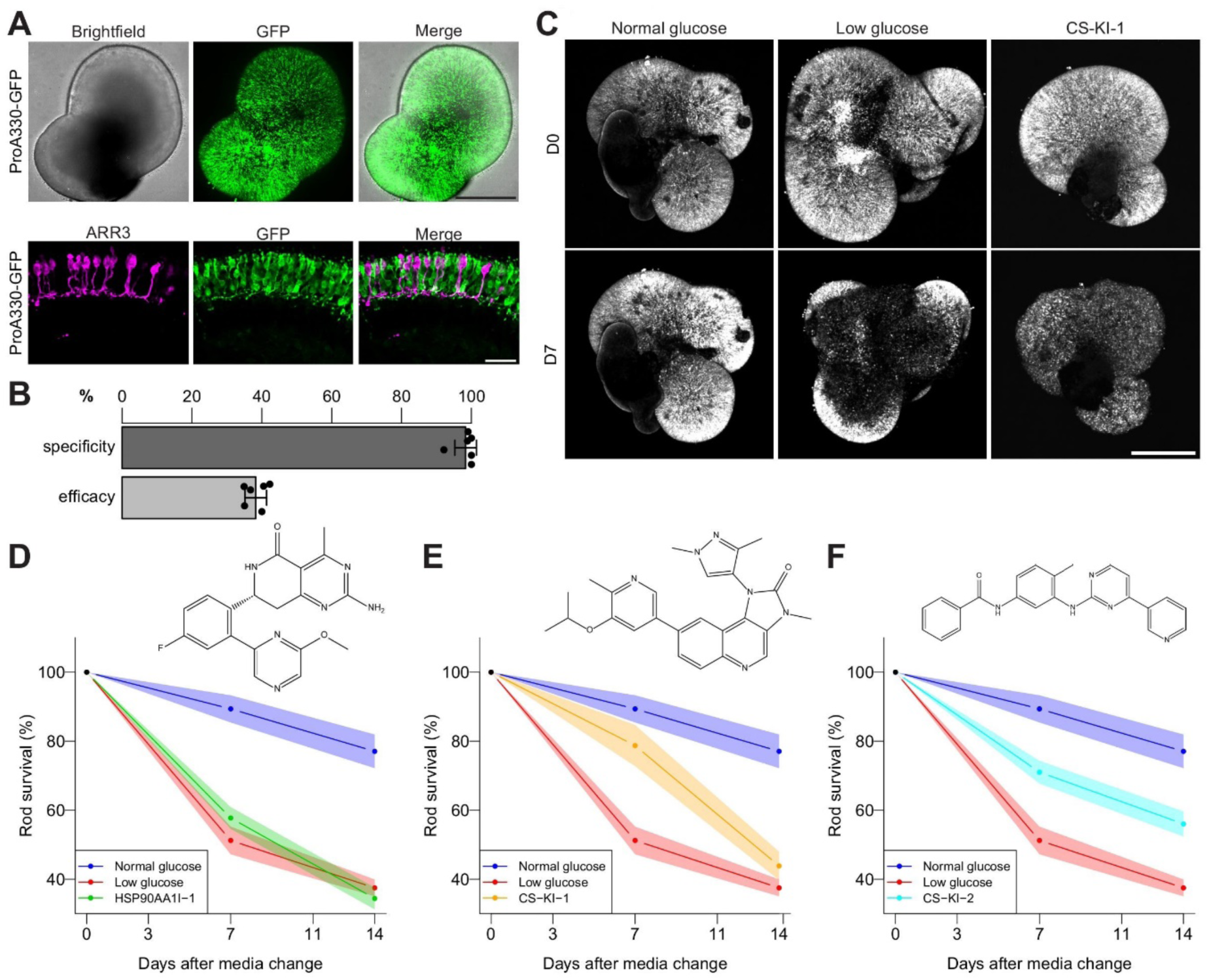
Effect of cone-saving compounds on rods. **A:** ProA330-GFP AAV transduction of human retinal organoids. Top: live image of a ProA330-GFP-transduced human retinal organoid (scale bar, 500 µm). Bottom: Confocal image of sectioned and stained transduced human retinal organoid (scale bar, 25 µm). Rods were identified as being present in the photoreceptor layer but negative for the cone-marker ARR3. ARR3, magenta; GFP, green. **B:** Quantification of the specificity and efficacy of rod labeling by ProA330-GFP AAV. Results are shown as mean ± sd. **C:** Example images of ProA330-GFP AAV-transduced human retinal organoids at D0 and D7 in either normal glucose, low glucose, or low glucose with CS-KI-1. GFP, white. **D-F**, Time course of rod survival in glucose-starved human retinal organoids treated with the indicated compounds for 14 days. Results are shown as mean ± se. Structures of the compounds are shown.

Similar to cones, rods also died during glucose starvation, with a survival by day seven of 51% in low glucose and 89% in normal glucose. The number of remaining rods was significantly different in low and normal glucose (low glucose, n=11, normal glucose, n=12, p<0.001, Mann–Whitney U test, Figures 5C and 5D). Rod survival dropped further after 14 days to 37 % in low glucose and 77% in normal glucose (low glucose, n=9, normal glucose, n=6, p<0.001, Mann–Whitney U test). Treatment with 10 µM CS-KI-1 or 10 µM CS-KI-2 led to an increase in rod survival after seven days of starvation (low glucose, n=11; CS-KI-1, n=18, p=0.002; CS-KI-2, n=18, p<0.001, Mann– Whitney U test, Figure 5D). However, only CS-KI-2 had a significant protective effect after 14 days (p<0.001, Mann–Whitney U test). In contrast to cones, we found no significant protective or detrimental effect of HSP90AA1I-1 on rod photoreceptors at any time.

### CS-KI-1 and CS-KI-2 nonfunctional analogs for identifying causal targets and transcriptional changes

A problem with identifying targets of CS-KI-1 and CS-KI-2 as well as transcriptional changes caused by CS-KI-1 and CS-KI-2 is that these compounds are known inhibitors of MTOR and PDGFR, respectively. At the same time, they also interact with unknown pathways that protect cones. As we argued before, the cone protection effect is most likely independent of both MTOR and PDGFR inhibition since other inhibitors of MTOR or PDGFR used in the primary screen did not save cones. Therefore, we used the following strategy to find MTOR- and PDGFR pathway-independent targets of CS-KI-1 and CS-KI-2 as well as to identify transcriptional changes induced by CS-KI-1 and CS-KI-2. We identified an MTOR inhibitor used in the primary screen that did not protect cones but had a very similar chemical structure to CS-KI-1; we named this CS-KI-1A (CS-KI-1 analogue). Similarly, we identified a PDGFRA inhibitor used in the primary screen that did not protect cones but had a very similar structure to CS-KI-2, which we named CS-KI-2A (Figure 6A).

**Figure 6:**
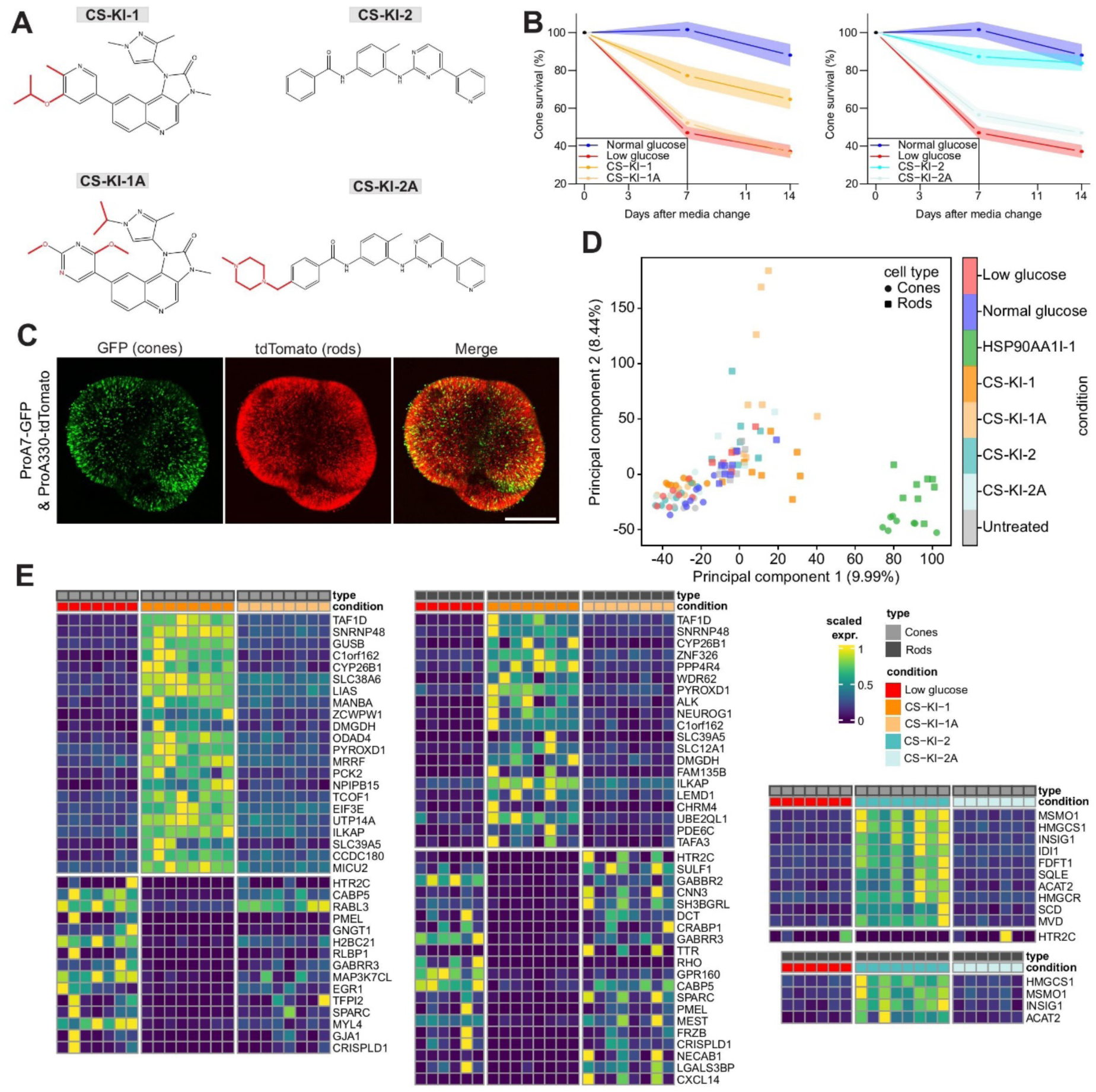
Transcriptomes of photoreceptors treated with cone-saving compounds. **A:** Chemical structures of CS-KI-1 and CS-KI-2 and their corresponding nonfunctional analogues. Structural differences between functional compounds and nonfunctional analogues are in red. **B:** Time course of cone survival in glucose-starved human retinal organoids with the indicated compounds for 14 days. Results are shown as mean ± se. Same data as in Figure 4H and 4I, but including the nonfunctional analogue compounds CS-KI-1A and CS-KI-2A **C:** Human retinal organoids transduced with ProA7-GFP (green) and ProA330-tdTomato (red, scale bar, 500 µm). **D:** Principal component analysis of transcriptomes under the indicated conditions. **E:** Differential gene expression comparing human retinal organoid cone and rod transcriptomes treated with the indicated conditions. Color scale indicates row-wise normalized gene expression levels. The top 40 differentially up- or downregulated genes are shown.

We repeated experiments with 10 µM CS-KI-1, CS-KI-1A, CS-KI-2, CS-KI-2A to assess their effectiveness in preventing cone death induced by glucose starvation. As in the primary screen, CS-KI-1A did not improve cone survival compared to low glucose and its effect was significantly different from that of CS-KI-1 (n=9, p=0.002, Mann–Whitney U test, Figure 6B). The same relation was observed between CS-KI-2A and low glucose and CS-KI-2 (n=10, p<0.001, Mann– Whitney U test, Figure 6B).

We then looked for targets of CS-KI-1 and CS-KI-2 that are not targets of CS-KI-1A and CS-KI-2A, respectively, using kinase profiling. Similarly, we looked for differential changes in transcription caused by CS-KI-1 and CS-KI-1A, as well as by CS-KI-2 and CS-KI-2A.

### Transcriptomic changes in cones and rods induced by cone-saving compounds

We generated dual color retinal organoids with green-fluorescent cones and red-fluorescent rods by transducing them with AAV9-PHP.eB capsid-coated ProA7-GFP and ProA330-tdTomato AAVs simultaneously (Figure 6C). This allowed us to isolate cones and rods from the same organoids using fluorescence-activated cell sorting (Figure S11). Subsequently, we analyzed their transcriptomes separately using bulk RNA-sequencing.

We aimed to identify genes and pathways involved in the survival of cones or rods, specifically those whose expression is altered by CS-KI-1 compared to CS-KI-1A and low glucose controls, or by CS-KI-2 compared to CS-KI-2A and low glucose controls. In addition, we also analyzed the effect of HSP90AA1I-1 inhibition on photoreceptor transcriptomes. We determined the transcriptomes of cones and rods in normal glucose control organoids and in organoids exposed to seven days of glucose starvation in the presence or absence of HSP90AA1I-1, CS-KI-1, CS-KI-1A, CS-KI-2 or CS-KI-2A.

Cells that were GFP positive expressed marker genes for cones, while tdTomato-positive cells expressed marker genes for rods. This was observed in both the normal and the low-glucose conditions, indicating, on the one hand, the effective isolation of the two cell types and, on the other hand, that the transcriptomic identity of cones and rods was not affected by glucose starvation (Figure S12).

The annotated target genes of HSP90AA1I-1, namely *HSP90AA1* and *HSP90AB1*, were highly expressed in both cones and rods. Expression of the annotated target genes of CS-KI-1, including *MTOR*, *PIK3CA*, *PIK3CB*, and *PIK3CD*, was low. Moreover, the annotated target genes of CS-KI-2, *PDGFRA* and *PDGFRB* were not or barely expressed in cones and rods (Figure S12). These findings further support the notion that HSP90AA1 and HSP90AB1 are targets of HSP90AA1I-1 in cones, while PDGFRA and PDGFRB are not targets of CS-KI-2 in cones and rods.

The remaining glucose is low in normal glucose medium after seven days and this may already subject organoids to a state of starvation. Therefore, we included a condition in which organoids received media exchanges every other day (‘untreated’ control). We found no differentially expressed genes when comparing cones and rods in normal glucose relative to untreated cones and rods (Figure S13). Principal component analysis showed the transcriptome of cones in low glucose to be close to the transcriptome of cones in normal glucose. This was also true for rods. Treatment with HSP90AA1I-1 induced a strong shift in the transcriptomes of both cones and rods (Figure 6D). CS-KI-1- or CS-KI-1A-treated rods differed markedly from untreated rods, which was not the case for cones. CS-KI-2 and CS-KI-2A both caused minor shifts in photoreceptor transcriptomes (Figure 6D).

Consistent with the principal component analysis, we identified only a limited number of genes differentially expressed between normal and low glucose controls: four genes were downregulated in cones, of which HSPA6 was the only gene also downregulated in rods when comparing normal to low glucose conditions (Figure S13). Following HSP90AA1I-1 treatment, cones exhibited differential expression relative to the low glucose condition in 776 genes, with 169 genes upregulated and 607 genes downregulated. In rods, 423 genes were differentially expressed, including 92 upregulated and 331 downregulated genes (Figure S13). Among these, 47 upregulated and 223 downregulated genes were differentially expressed in both cones and rods. Gene set enrichment analysis (GO terms) revealed that genes involved in RNA processing, which includes transcription, splicing, and translation, were upregulated in both cones and rods treated with HSP90AA1I-1 compared to the low glucose controls. This upregulation explains the significant number of genes differentially expressed. Additionally, genes associated with the unfolded protein response were upregulated in cones (Molecular Signature Database). This finding aligns with the known function of HSP90 as a molecular chaperone that assists in protein folding (Figure S14).

For CS-KI-1, we examined genes that showed differential expression when comparing the transcriptomes of samples treated with CS-KI-1 against those treated with CS-KI-1A and the low glucose control samples. We sought to identify genes that are differentially expressed in CS-KI-1 compared to low glucose controls, but not CS-KI-1A and the low glucose controls. We identified 22 upregulated and 15 downregulated genes in cones, and in rods we found 39 upregulated and 130 downregulated genes (Figures 6E and S15). Notably, 10 upregulated or downregulated genes were common to both cones and rods. Gene set enrichment analysis (Molecular Signature Database) revealed that apoptotic genes were downregulated in both rods and cones. Moreover, cones exhibited downregulation of inflammatory response genes, including those in TNF-alpha signaling via NFKB and Interferon Alpha responses. This suggests that the downregulation of both inflammatory and non-inflammatory cell death pathways could be involved in enhancing cone survival (Figure S16).

We adopted a similar approach for CS-KI-2, examining genes differentially expressed when comparing CS-KI-2-treated samples to those treated with CS-KI-2A and to low glucose controls. Consistent with the principal component analysis, CS-KI-2 treatment resulted in fewer differentially expressed genes. In cones, we identified 10 significantly upregulated genes, with *HMGCS1*, *MSMO1*, *INSIG1*, and *ACAT2I* also showing significant upregulation in rods. Additionally, *HTR2C* was found to be downregulated in cones (Figure 6E). Gene set enrichment analysis in both cones and rods indicated that genes involved in mTORC1 signaling and cholesterol homeostasis were upregulated following CS-KI-2 treatment. Furthermore, genes related to fatty acid metabolism showed upregulation in cones (Figure S16).

These findings indicate that treatment with HSP90AA1I-1, CS-KI-1, or CS-KI-2 induces highly distinct gene expression alterations in photoreceptors, each manifesting varying degrees of transcriptomic changes. Despite numerous shared expression modifications across treatments in both rods and cones, there are also notable cell type-specific effects.

### Photoreceptor-saving targets of CS-KI-1 and CS-KI-2 revealed by kinase profiling

To investigate causal targets of CS-KI-1 and CS-KI-2 for saving cones or rods, we conducted biochemical kinase profiling for these compounds and their nonfunctional analogues CS-KI-1A and CS-KI-2A. We assessed the activity of 350 human kinases after treatment with each of the four compounds, comparing their activities at 10 µM against a vehicle control (Figures 7A, 7B and Table S4).

**Figure 7:**
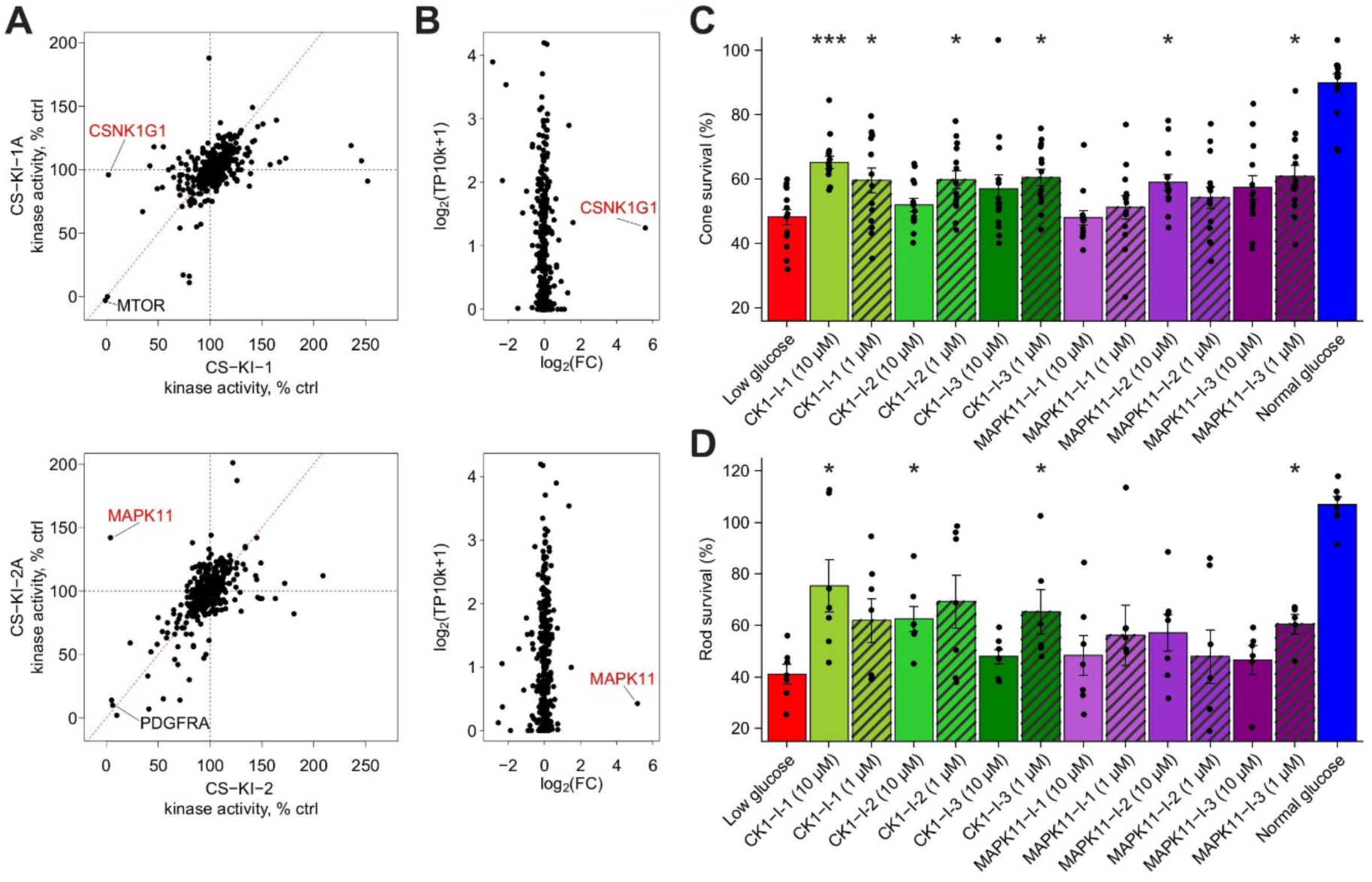
Kinase profiling and CK-1 and MAPK11 inhibition. **A:** Kinase profiling of 350 human kinases treated with CS-KI-1, CS-KI-1A, CS-KI-2 or CS-KI-2A. Percentage value of kinase activity compared to a control reaction without inhibitors. Colors indicate average expression level of the kinase gene in low glucose organoid cones. Dotted black lines, 100% kinase activity. Dotted red lines, unity line. TP10k, transcript counts per 10,000. **B:** Differential kinase activity comparing active compounds and their analogues and average expression levels in organoid cones. FC, foldchange. TP10k, transcript counts per 10,000. **C-D:** Effects of CK-1 and MAPK11 inhibitors on cone (C) and rod (D) survival under low glucose conditions. Results are shown as mean ± se with * denoting an adjusted p-value<0.05. Black dots indicate individual organoid cone (C) or rod (D) survival.

CS-KI-1 and its nonfunctional analogue CS-KI-1A had little effect on the activities of most of the 350 kinases; however, there were a few exceptions. As expected, both CS-KI-1 and CS-KI-1A induced complete inhibition of MTOR. Remarkably, the activity of only one kinase, the gamma 1 form of casein kinase 1 (CSNK1G1), was fully suppressed by CS-KI-1 but not influenced at all by CS-KI-1A (Figures 7A, 7B and Table S4). CSNK1G1 is therefore a potential target for the cone- and rod-saving effects of CS-KI-1 that is independent of MTOR inhibition.

Similarly, the activity of most kinases was not influenced by CS-KI-2 or its nonfunctional analog CS-KI-2A. Both CS-KI-2 and CS-KI-2A induced full inhibition of PDGFRA. The activity of only one kinase, mitogen-activated protein kinase 11 (MAPK11), was fully blocked by CS-KI-2 but not inhibited at all by CS-KI-2A. Indeed CS-KI-2A increased the activity of MAPK11 (Figures 7A, 7B and Table S4). MAPK11 is thus a potential target for the cone- and rod-saving effects of CS-KI-2 that is independent of PDGFRA inhibition.

Therefore, we tested three known inhibitors of casein kinase 1 (CK-1) and MAPK11 on both organoid cones and rods using the two-color organoids (GFP in cones and tdTomato in rods) described above at either 10 or 1 µM inhibitor concentrations. Strikingly, all three CK-1 inhibitors significantly improved cone survival at one or both concentrations (CK1-I-1, 10 µM, p<0.001; 1 µM, p=0.04; CK-1-I-2, 1 µM, p=0.01; CK1-I-3, 1 µM, p=0.01; n=14, Mann–Whitney U test with Benjamini Hochberg correction for multiple testing). This was also true for rods (CK1-I-1 10 µM, p=0.01; CK1-I-2 10 µM, p=0.009; CK1-I-3, 1 µM, p=0.009; n=7, Mann–Whitney U test with Benjamini Hochberg correction for multiple testing). Two out of three MAPK11 inhibitors had a significant positive effect on cone survival (MAPK11-I-2, 10 µM, n=14, p=0.01; MAPK11-I-3, 10 µM, n=13, p=0.01) and one of the MAPK11 inhibitors significantly improved rod survival (MAPK11-I-3 1µM, n=7, p=0.009; Mann–Whitney U test with Benjamini Hochberg correction for multiple testing, Figures 7C, 7D and Table S5).

*MAPK11* and *CSNK1G,* along with other casein kinase 1-encoding genes, are expressed in both organoid rods and cones, as well as in human retina rods and cones, both in the fovea and the periphery (Figure S17).

## DISCUSSION

### Large-scale compound screen in human organoids

We have performed a cell type-focused compound screen in ∼20,000 human organoids using 2,707 compounds with annotated targets. We used a modified AMASS method^22^ to produce replicable, multilayered human retinal organoids in large quantities. We induced the death of cones by lowering glucose for seven days and monitored cell death by imaging GFP that was expressed specifically in cones. We identified compounds that protected cones, compounds that damaged cones, and compounds that had no effect. The results of the primary screen, including the names and unique identifiers of all the compounds, their annotated targets, and their effect on cone survival are publicly available at https://ConeTargetedCompoundScreen.iob.ch.

Among the photoreceptors, we have focused on cones since the dysfunction of cones leads to blindness, while rod dysfunction leads to no or minor visual disabilities^25^. We labeled cones with AAVs carrying a cone-specific promoter^14^. The use of AAVs has the advantage over using organoids with genetically labeled cell types in that the target cell type can be rapidly changed across organoids with the same genetic background. Different genetic backgrounds can lead to variation in the structure and cell-type composition of retinal organoids^22^. AAV-based labeling allowed us to test some of the compounds on rods of organoids grown from the same induced pluripotent cell line, using AAVs carrying a rod-specific promoter.

To find molecules that either protect or damage cones, the screen had to satisfy several conditions. First, that we could compare the number of cones before and after the induction of death in each organoid. This is important since the number of cones varies across individual organoids. Second, that the time period in which death happens is short, within days. The reason for this is practical: it allows the testing of more organoids and compounds. Furthermore, organoids do not need a medium exchange within a few days, thus allowing live-imaging of the same organoids in the same positions. The short time period to detect cone death required that the degeneration of cones is induced synchronously and progresses rapidly within and across organoids. An alternative would have been the use of patient-derived or CRISPR-engineered human organoids with either primary or secondary cone degeneration. However, cone degeneration in such organoids has so far not been observed. If it is observed in the future, it is likely to happen over a long period of time and asynchronously. Third, that during the period in which cone death occurs, organoids should reach a state in which about half of the cones die and half remain alive, thus allowing a search for compounds that protect cones and those that damage cones. Fourth, the use of a large number of organoids. On the one hand, this allows for the screening of many compounds; on the other hand, the large numbers allow study of potentially confounding variables that affect cone death. Such a confounding variable was the number of cones before starvation: the percentage of cones dying in the low glucose condition is lower in small organoids with fewer cones. Interestingly, the fact that human retinal organoids need to grow for ca. 30 weeks to yield developed cones did not hinder the screen. Since batch-to-batch variability in cone survival was not detected, different batches of organoids could be initiated in a short time period, one after the other, and thus many organoids could be produced semi-parallel.

We developed three cell-counting algorithms for counting cones from the stack of images taken of each organoid. Each image stack was ∼260 MB and the entire dataset was ∼5 TB. We used the 3D-additive-count as the primary measure, since MIP-count loses information on the edges of the organoids, where different cones are merged in the maximum intensity projection, and since the 3D-count is computationally time consuming.

In the course of this work, we developed a promoter, ProA330, that allows specific labeling of rods in human organoids. Thus, we could mark cones and rods in the same organoid using ProA7-driven GFP and ProA330-driven tdTomato from two different AAVs. In the future, these dual color organoids will enable human retinal organoid screens focused on rods and cones simultaneously.

Since most organoids have more than one cell type and in many organs diseases affect specific cell types, the methods of and the lessons learned from the cell type-focused screen we conducted in human retinal organoids are likely to be useful for performing large-scale screens in other types of human organoids.

### Compounds that damage cones and HDAC inhibition

We identified 146 compounds that caused significant damage to cones in human retinal organoids. We found that HDAC I/II inhibitors with a wide range of targets lead to significant damage to cone photoreceptors. This damaging effect is proportional to the logarithm of the compound concentration. More selective HDAC inhibitors resulted in less damage to cones in human organoids. It was shown that inhibiting HDACs broadly in the developing mouse retina using trichostatin-A reduces the expression of transcription factors, including Otx2, Nrl, and Crx, which are important for rod development. Additionally, inhibiting HDACs in mouse retinal explant cultures resulted in a complete loss of rod photoreceptors^65^. Conversely, overexpression of Hdac4 in a mouse model of retinal degeneration extended the survival of photoreceptors^66^. Taken together, this work on cones of human retinal organoids together with the work on rods in mouse retina^65,66^ suggests that broad HDAC-inhibition is damaging to both cone and rod photoreceptors. This is particularly relevant because various HDAC inhibitors are currently being tested in clinical trials or have already been approved for cancer treatment^67^. Therefore, it will be important to monitor the structure and function of the retina in patients receiving these treatments.

Interestingly, some studies have found that broad inhibition of HDACs by trichostatin-A in mouse models of retinal degeneration can have positive effects on cones^47,48^, and another study has suggested that targeted inhibition of HDAC6 positively impacts the cones of mice^49^. Currently, it is unclear why the impact of broad HDAC inhibition on cone cells differs between humans and mice.

### Compounds that protect human cones

Considerable effort has been made to find ways to protect cones in animal models of retinal degeneration^32–46^ and some identified modifiers of cone death are currently being evaluated in clinical trials^68^. However, no treatment for protecting cones in photoreceptor diseases such as macular degeneration or retinitis pigmentosa has yet received approval for use in humans.

In this study, we adopted a complementary approach by screening compounds based on their ability to enhance human cone survival in organoids.

We induced cone death by glucose starvation not only because this triggers synchronous and rapid cone death, but also because it has been demonstrated that enhanced glucose uptake and glycolysis promote cone photoreceptor survival in animal models of photoreceptor degeneration. This suggests that cones experience starvation-induced death in some photoreceptor diseases^32,38,50–52^. The extent to which glucose starvation mirrors what occurs in degenerative diseases requires future assessment. In primary and secondary screens, we isolated four compounds that had a significant protective effect on cones after seven days of glucose starvation.

Two of the four compounds that protected cones, HSP90AA1I-1 and HSP90AA1I-2, target the same protein, HSP90. An earlier study demonstrated that a single dose of an HSP90 inhibitor improved visual function and delayed photoreceptor degeneration in a P23H transgenic rat model^69^. Despite the protective effects on photoreceptors observed in both human organoids and rats, we argue that inhibiting HSP90 is not suitable for preserving human cones. First, inhibiting HSP90 caused significant damage to cones in human organoids after 14 days of treatment. Second, HSP90 inhibition resulted in major alterations in the transcriptomes of both cone and rod photoreceptors in human organoids. HSP90 has been described as an orchestrator of transcription, a finding that aligns with our results^70–72^. Third, administering HSP90 inhibitors systemically to dogs was found to induce damage to photoreceptors and to impair vision^73^.

The other two compounds that protected cones, CS-KI-1 and CS-KI-2, are kinase inhibitors. Three pieces of evidence suggest that the currently labeled targets, including MTOR (CS-KI-1) and PDGFRA (CS-KI-2), are not responsible for the effects of CS-KI-1 and CS-KI-2 in cones. First, many inhibitors targeting the same labeled targets had no impact on the survival of cones. Second, the IC50 values of different inhibitors were not correlated with the magnitude of the protective effect on cones. Third, the targets of CS-KI-2 (PDGFRA and PDGFRB) are not or barely expressed in organoid cones.

Transcriptomic analysis of photoreceptors treated with CS-KI-1 identified differentially expressed genes relative to CS-KI-1A and low glucose that lead to the downregulation of stress responses as well as inflammatory and non-inflammatory cell death pathways. Both apoptotic and necrotic cell death have been described as potential pathways for disease-related photoreceptor degeneration^74–77^. Interestingly, CS-KI-2 elicited a distinct response characterized by the upregulation of a smaller set of genes involved in cholesterol homeostasis and mTORC1 signaling. Previous research in a mouse model of retinitis pigmentosa demonstrated that activation of the mTOR pathway can delay cone cell death^38^.

By performing a kinase screen using CS-KI-1 and CS-KI-2 and their nonfunctional chemical analogs CS-KI-1A and CS-KI-2A, we identified MTOR- and PDGFRA-independent kinase targets of these compounds: CSNK1G1 for CS-KI-1 and MAPK11 for CS-KI-2. We then showed that inhibitors of casein kinase 1 (CSNK1G1 is a member of the casein kinase 1 family) and inhibitors of MAPK11 protect both cones and rods. CK-1 inhibition has been identified before as a potential target to counteract neurodegeneration^78–80^. Identifying casein kinase 1 and MAPK11 as elements in cone-or rod-protecting pathways could pave the way to the discovery of more potent compounds acting on the same targets, as well as to other means of interfering with these targets specifically in cones or rods.

## ACKNOWLEDGMENTS

We thank V. Moreno-Juan, A. Muller, Á. Herrero-Navarro, T. M. Rodriguez, A. Fratzl, T. Dalmay and P. King for comments on the manuscript. We thank other members of the Roska laboratory for their discussions on the manuscript. The project was supported by a European Research Council advanced grant (HURET N°883781), Swiss National Science Foundation Synergia grant (CRSII3_141801), Swiss National Science Foundation grant (310030_212186), a Louis-Jeantet Foundation award, a Körber Foundation award, and the NCCR Molecular Systems Engineering grant to B.R.

## AUTHOR CONTRIBUTIONS

S.E.S. designed and conducted experiments, including the primary and secondary screens, cultured and prepared organoids, including dissociations, analyzed and interpreted data, including the screening data, and wrote the paper. V.J.A.M. conducted experiments, including the primary and secondary screens, cultured and prepared organoids, and designed illustrations. Z.R. developed the cone and rod counting algorithms and analyzed data. S.P.C. analyzed and interpreted RNA-seq data. S.C., O.G., I.G. and I.C. designed and conducted screening experiments and interpreted data. S.R. analyzed and interpreted screening data. L.U conducted experiments, cultured and prepared organoids, including dissociations. P.T.K. conducted experiments for ProA330 discovery. A.V. and J.I. produced AAVs. S.M. performed FASC-sorting. R.A.S. and V.H. conducted RNA isolation and sequencing. Y.H. performed staining and imaging on organoid sections. S.P. supervised RNA-seq experiments. M.C. analyzed data. J.J. supervised AAV production and experiments for ProA330 discovery. C.S.C. supervised RNA-seq analysis. M.D. supervised screening experiments. D.K.B. supervised screening experiments and analysis, M.R. supervised organoid production and screening experiments, V.U. supervised screening experiments, B.R supervised experiments, analyzed and interpreted data, and wrote the paper.

## DATA AVAILABILITY

The primary screen dataset summary is available at https://ConeTargetedCompoundScreen.iob.ch. Additional data supporting the findings of this study are available from the authors upon request.

## CODE AVAILABILITY

The custom code used for data analysis is available upon request from the authors.

## BIOLOGICAL MATERIALS AVAILABILITY

All unique biological materials used are readily available from the authors

## MATERIALS AND METHODS

### Generation of retinal organoids

Retinal organoids were generated as previously described^22^, with the modifications given below. All experiments in this study were performed using organoids that were 30- to 32 weeks old.

### Induced pluripotent stem cell culture

Organoids were derived from the induced pluripotent stem cell line 01F49i-N-B7^22^. The cells were cultured at 37°C and 5% CO_2_ in a humidified incubator, using mTesR1 medium (STEMCELL Technologies, #85850) on Matrigel-coated (Corning, #356230) 6-well plates (Corning, #3516). The culture medium was replaced daily and cells were passaged weekly using 0.5 mM EDTA (Invitrogen, #15575020) in PBS without CaCl_2_/MgCl_2_ applied for 3-5 min to facilitate detachment of cells as small clumps for subsequent seeding.

### Embryoid body formation and culture

Induced pluripotent stem cells were detached and a single-cell suspension was created using 0.5 mM EDTA (Invitrogen, #15575020) for 3 min, followed by a 3-min Accutase (Thermo Fisher Scientific, #00-4555-56) treatment at 37°C. Embryoid body formation took place in 256 microwell-hydrogels with ∼250-300 cells seeded per microwell. The hydrogels were generated using a MicroTissues 3D Petri Dish micro-mold (Sigma Aldrich, Z764000) and 2% agarose (Thermo Fisher Scientific, #R0491). Each hydrogel was cultured in a 12-well plate (Corning, #3513) in neural induction medium DMEM / F12 (GIBCO, #31331‒028), 1% N2 Supplement (GIBCO, #17502‒048), 1% NEAA Solution (Sigma, #M7145) and 2 mg/mL heparin (Sigma, #H3149‒50KU) for one week with daily medium exchanges. Embryoid bodies that formed in one 256 microwell-hydrogel were detached from the hydrogel and distributed evenly across three wells of a Matrigel (Corning, #356230)-coated 6-well plate (Corning, #430166).

### Early organoid culture and checkerboard scraping

Organoids in 6-well plates cultured with daily medium exchanges started to form 2D confluent structures. For the first 16 days, they were cultured in neural induction medium. The medium was subsequently changed to 3 parts DMEM (GIBCO, #10569‒010): 1 part F12 medium (GIBCO, #31765‒027) (‘3:1 medium’), supplemented with 2% B27 without vitamin A (GIBCO, #12587010), 1% NEAA Solution (MERCK, #M7145), and 1% penicillin / streptomycin (GIBCO, #15140‒122). Checkerboard scraping was performed between days 28 and 30 of culture as described previously^22^.

### 3D-organoid culture

Aggregates from four wells of a 6-well plate were transferred to one 175 cm^2^ tissue culture flask (Thermo Scientific, #159926) previously treated with an anti-adherence solution (StemCell Technologies, #07010). Flasks containing organoids were filled with 35 - 45 mL of medium, which was replaced 1-3 times per week. The flasks were maintained in 3:1 medium for 6 weeks of culture. The medium was supplemented subsequently with an additional 10% FBS (Millipore, # ES-009-B) and 100 µM Taurine (Sigma, #T0625‒25G) until week 10. Until week 14, the medium was further supplemented with 1 µM retinoic acid (Sigma, #R2625). After this period, the retinoic acid concentration was reduced to 0.5 µM and the B27 supplement was replaced with N2 supplement (GIBCO, #17502‒048) for the remaining duration of the culture. For easy access to organoids for experiments, they were transferred to round cell culture dishes (Thermo Fischer, #101VR20). Aggregates lacking neuroepithelium were removed just before the experiments.

### Organoid fixation, sectioning and staining

Organoids were fixed in paraformaldehyde for 4 h at 4°C and washed three times for 10 min each in PBS. They were then submerged in PBS containing 30% sucrose for cryopreservation. Fixed organoids were stored at -80°C.

For sectioning, the organoids were embedded in a solution of 7.5% gelatine and 10% sucrose in PBS. The embedded samples were then frozen and sectioned into 25-µm-thick slices using a cryostat (MICROM International, #HM560).

Immunostaining was carried out as described previously^22^. Briefly, slides were dried for 30 min at room temperature, then rehydrated in PBS for 5-10 min. They were then blocked with a solution containing 10% normal donkey serum (Sigma, #S30‒100ML), 1% BSA (Sigma, #05482‒25G), 0.5% Triton X-100 (Sigma, #T9284500ML), and 0.02% sodium azide (Sigma, #S2002‒25G) at room temperature for 1 h. Sections were then treated with primary antibodies (Supplementary Table 6) in a similar blocking solution but with 3% normal donkey serum for 24 h. After three washes in PBS with 0.1% TWEEN 20 (Sigma, #P9416100ML) for 10 min each, the slides were exposed to secondary antibodies (Thermo Fisher Scientific, donkey secondary antibodies conjugated to Alexa Fluor 488, 568, or 647) diluted 1:500 and Hoechst 33342 (Thermo Fisher, #62249) diluted 1:10,000 in the same buffer as the primary antibodies for 2 h. This was followed by two 10-min washes in PBS with 0.1% Tween and a 15-min wash in PBS. Slides were finalized with ProLong Gold (Thermo Fisher Scientific, #P36934) before sealing.

### Imaging stained cryosections

Images of representative regions of the organoids were captured using a spinning disc confocal microscope (Olympus IXplore SpinSR). The microscope was adjusted to either 20x or 40x magnification and images were taken across multiple Z-planes. All captured images are shown as maximum intensity projections.

### AAV production

Adherent HEK293T cells (ATCC, #CRL3216) were cultured in a 5-layer CellSTACK (3,180 cm^2^; Corning, # CLS3319) for AAV vector production. These cells were co-transfected with an AAV transgene plasmid, an AAV helper plasmid encoding the AAV Rep2 and Cap proteins for the selected AAV9-PHP.eB capsid^58^, and the pHGT1-Adeno1 helper plasmid carrying adenoviral genes (kindly provided by C. Cepko, Harvard Medical School, Boston, USA) using PEIMAX (Polyscience, #POL24765-1). Plasmids were mixed in 98 mL DMEM (Thermo Fischer, #11965-092) and incubated for 5 min. PEIMAX was then added to the DMEM-diluted DNA. After an additional 10-min incubation, the DNA-PEIMAX complex was added to the cells. After 60 h, the culture medium was supplemented with 250 mL fresh DMEM containing 1% Pen-Strep (Thermo Fischer, #15140-122). AAV vectors present in cells and the culture medium were harvested approximately 5 days post-transfection.

### Purification of AAVs

AAVs were purified either from cell culture medium alone or from both cells and cell culture medium. The cell culture supernatant was first cleared by centrifugation at 1400 x g for 15 min (5920R; Eppendorf) and then filtered through a 0.45-µm PES filter (Merck Millipore, #S2GPU02RE). The cell pellet was resuspended in 11 mL lysis buffer (150 mM NaCl, 20 mM Tris-HCl pH 8.0) and subjected to three freeze-thaw cycles. To remove cell debris, the cell lysate was centrifuged at 4,000 x g for 30 min and the resulting supernatant filtered through a 0.45-µm filter.

Both the filtered cell culture medium and the cell lysate were treated with Turbonuclease (Accelagen, #N0103L) at 50 U/mL for 1 h at 37°C. The sample was then loaded onto an affinity column (POROS CaptureSelect AAVX; ThermoFisher, #A36652) and eluted with a solution of 0.1M glycine (Merck, #1.04201.1000), 0.25M arginine (Sigma, A5006), 0.2 M NaCl (pH 2.7; Sigma, 31434), following an extensive wash with 20 times the column volume of 500 mM NaCl, 50 mM Tris at pH 7.3 (Merck, 93350), and 0.01% Pluronic F-68 (Thermo, #24040032). The eluted AAVs were immediately neutralized with 1/11 volume of 1 M Tris-HCl pH 10. The purified AAV vectors were then concentrated as needed in sterile PBS + 0.001% Pluronic F-68 using a spin filter (Amicon Ultra Centrifugal Filter Units; Millipore Sigma, UFC910096; molecular cutoff 100 kDa).

### AAV titration

Encapsidated viral DNA was quantified using TaqMan RT-PCR (Thermo Fischer, #4444557) targeting the ITR sequences (forward primer: GGAACCCCTAGTGATGGAGTT; reverse primer: CGGCCTCAGTGAGCGA; probe: [6FAM]CACTCCCTCTCTGCGCGCTCG[BHQ1]) relative to a linearized ITR-containing plasmid as a standard. Prior to quantification, AAV particles were denatured using Proteinase K (Thermo Fischer, #11501515). The titers were then calculated and expressed as genome copies per mL.

### AAV transduction of organoids in 96-well plates

For small-scale experiments (Figures 5, 6, 7, S9, S11-S15), individual organoids were transferred to a single well of an ultra-low attachment U-bottom 96-well plate (Corning, #7007) after 25 to 26 weeks of maturation. AAVs were diluted in culture medium to a concentration of 3.3 × 10^12^ genome copies per mL. The existing medium in the wells was removed from the organoids and replaced with 30 µL of the AAV solution. After an incubation period of 4-5 h, an additional 70 µL of medium was added per well. After 24 h, 100 µL of medium was added on top per well. After a further 24 h, 150 µL of medium was replaced per well. Transduced organoids were then cultured for 4 to 5 weeks at 37°C and 5% CO_2_ before the onset of further experiments. During this period, the medium (150 µL per well) was replaced 3 times per week.

### Bulk AAV transduction of organoids

For large-scale experiments (Figures 5, 6, S1-S8 and S10), organoids were transduced with AAVs in their original flask (Sigma Aldrich, #Z764000). AAVs were diluted in culture medium to a concentration of 1 × 10^13^ to 2.5 × 10^13^ genome copies per mL. The flasks were placed upright and the medium aspirated from the organoids. AAV solution was then added at 8 mL per flask and the flasks incubated in the upright position for 24 h. Following this, 32 mL of medium was added to each flask and the flasks were then laid flat. After a further 24 h, the medium was exchanged completely. The transduced organoids were cultured for an additional 4-5 weeks at 37°C and 5% CO_2_ with a medium exchanged once a week.

### Low-throughput imaging

All non-screening imaging (Figures 1, 4G-4I, 5, 6 and S9) was conducted using a spinning disk confocal microscope (Olympus IXplore SpinSR) with a 4x or a 10x objective. For live imaging, organoids were kept in a humidified chamber maintained at 37°C with 5% CO_2_. The contrast and brightness settings for images captured from the same organoid across different timepoints were the same.

### Glucose starvation

After 30 weeks of maturation and four weeks after AAV transduction, organoids were transferred into an ultra-low attachment U-bottom 96-well plate (Corning, #7007) with 150-200 μL medium. The remaining medium was then reduced to approximately 60 µL per well.

The low glucose starvation medium was composed of 3:1 medium supplemented with an additional 10% heat-inactivated FBS (Millipore, #es‒009‒b), N2 supplement (GIBCO, #17502‒ 048), 100 μM taurine (Sigma, #T0625‒25G), and 0.5 µM retinoic acid. Instead of the standard DMEM, DMEM with no glucose (Thermo Fisher, #11966025) was used. The normal glucose medium was prepared similarly but with regular DMEM containing 25 mM glucose (Thermo Fisher, #10569010). The low glucose medium still contained a small amount of glucose due to the F12 medium (GIBCO, #31765‒027) and possibly the FBS.

Prior to the addition of the respective experimental conditions, organoids including the normal glucose controls were washed twice with 120 μL of low glucose medium. This process was conducted manually or, for screening experiments, with a 96-well head Selma pipettor (Cybio, #OL7001-26-212) fitted with 60-μL tips (Cybio, #OL3800-25-735-P).

### Glucose consumption measurement

Organoids were transferred to an ultra-low attachment U-bottom 96-well plate (Corning, #7007) and subjected to glucose starvation. On each measurement day, 5 µL of medium was collected from the 180 µL of medium for each of the 10 organoids per condition. For the initial measurement, which involved only medium without organoids, three replicates were taken. The glucose concentration was subsequently determined using a Glucose Colorimetric Detection Kit (Invitrogen, #EIAGLUC). The assay results were read using a Hidex Sense Microplate Reader.

### Compound preparation, dilution, and addition

A set of 2,707 annotated compounds selected for screening on retinal organoids was sourced from the Mode-of-action (MOA) compound library^57^. The compounds, originally at a stock concentration of 10 mM in 100% DMSO, were plated in 384-well low dead volume plates (Labcyte, #LP-0200). Using the ECHO acoustic liquid handler (Labcyte, #Echo 555), 225 nL of the compounds was transferred into sterile 96-well polypropylene U-bottom microplates (Greiner, #65026). These plates were stored overnight at 4°C in a confined environment to prevent evaporation.

The following day, the plates were brought to room temperature and the compounds diluted 666 times by adding 150 µL of low glucose medium with the multidrop combi dispenser (Thermo Scientific, #5840300). Using a 96-well head Selma pipettor (Cybio, #OL7001-26-212) equipped with 60-µL tips (Cybio, #OL3800-25-735-P), 120 µL of culture medium was removed from the ultra-low attachment U-bottom 96-well assay microplates (Corning, #7007) containing the organoids. Then, using the same pipettor, 120 µL of compounds diluted in medium were pipetted from the intermediate 96-well plates into the plates containing the retinal organoids. The assay plates were then incubated (5% CO_2_; 37°C; humidified environment) in an automated incubator (Thermo, #incubator Cytomat 10 C 450) until imaging. Each compound was tested in five replicates at a final concentration of 10 µM. All vehicle controls were 0.1 % DMSO in low glucose or normal glucose medium. In the secondary screens, the compounds were tested at four different concentrations (10; 1; 0.1 and 0.01 µM) using the compound transfer process as in the primary screen.

HSP90AA1I-1, CS-KI-1 and CS-KI-1A (Figures 4G-4H, 5C-5D, 6 and S9, S12-S15) were newly synthesized by Enamine. CS-KI-2 and CS-KI-2 were purchased from MolPort (Figures 4I, 5D, 6, S9, S12-S15). These were dissolved in 90% DMSO prior to manual dilution in low glucose medium. The vehicle controls for these experiments involved 0.09% DMSO. If not indicated otherwise HSP90AA1I-1 was added at a concentration of 1 µM and CS-KI-1, CS-KI-1A, CS-KI-2 and CS-KI-2A at a concentration of 10 µM. All CK-1 and MAPK11 inhibitors were purchased from MedChemExpress or StemCell Technologies.

### Automated imaging

Confocal images for all screening experiments were captured using a 4x objective lens (Olympus UPLSAPO, NA=0.16) on an automated spinning disk confocal microscope equipped with a sCMOS camera (Yokogawa, CV7000). The samples were maintained in a 5% CO_2_ and 37°C environment during acquisition. Images were acquired at 24 different confocal planes, each separated by a 34-µm interval, to cover the entire organoid. This was followed by the acquisition of a stack of seven brightfield images at 100-µm intervals. After image acquisition, each plate was incubated (5% CO_2_; 37°C; humidified environment) for seven days and then re-imaged using the same procedure.

### Promoter ProA330

The general design and the testing of ProA series promoters have been described previously^14^. The AAV serotypes used were AAV9-PHP.eB^58^ for human retinal organoids and AAV8-BP2^81^ for mouse injections. This new promoter has the following sequence:

AACCCAAGAAATTACAGGCTGAAACCAGAAAAGAACACATTAAAGCACCAAGAGAAAGTTGGAGTGGGTTGAAGGGAAACAGATTTTTAAAGTTAAGGCTCTGTGAAATGGGTAGAATTAACTACAGGTTAAAAATAAAATGTTAACTAAAGGTTGCCTCTGAGTAACAGGATTATGGGTGATTTTAATTGTCTTCTTTGTGTATGTTCAACAGTGACTATAATATGTATTACTTTTGGAATAAAGGAAAACCTGAAAGGTGTGTTGTTTTATAAGGGCCCTTAGGTTGCCAAAATTAGAGTCATTGAAATCTAAAGCTGATAAAAACTTTAGTGCAAAGATTGTGACATGGGAGACTACACATACCAGATCCATAATGTACATGAGGACAGTAGGCCGAGGGGCCCTGCACATTGAAAGCCCACATGGGAGAAGCCCTTGGGAAGGGGAGTGGAAGGATGAGGCAAGGGGCCGGGGGGATGCAGAGGCTGGCAGGCAGTCATTTCTCAGCTTCAGCCATTCCCGCCATGGGGGAATGTGGACAGAGAAGCCAAACAAATCTCCTAAACAGTAAATGTCAGTCTTCTGTGTCAGATATTTAAGAAAACTAACAGAGGTCAGAGAAGACACACCTACAGCAAGTAGACTGTCCCTGTGCTGCCTTTTTGCAACCCCTGCTTTGGCAGTGCTCAAGCCCACCTCCTGCTCTGTGCAGACATCTCTTCTTTGCTCTTACTAGACCAAGGTGAAAGAAAACTCTCACCTTCTCCCATCTGGCCCCACAGCATCTGGAACACACTGATCCTCATAATCCTTGTTCTTGAGAAATATTAATGACTTAATCTCCCAAGCTTGCTCCCTCTCCTGTGCAGGCCATCTCAGTATGTTTTGCAGACAAGACCCAGAGAAGTCCAGACTGGACTTGTTGCAGACTGCAAAACTGCCATTGGAAGGCCTCCGTCCCAGTCCTTCTACAGAGTAGCCAGTGGGATTCCCAGCC

### Organoid dissociation and FACS-sorting for RNA-seq

For bulk RNA sequencing, organoids at week 26 were co-transduced with a ProA7-GFP^14^ construct (cone-specific promoter driving GFP expression) and ProA330-tdTomato (rod-specific promoter driving tdTomato expression). Four weeks post-transduction, organoids were subjected to their corresponding treatments (untreated, normal glucose, low glucose, and low glucose with either HSP90AA1I-1, CS-KI-1, CS-KI-1A, CS-KI-2 and CS-KI-2A) in 96-well plates. After seven days of treatment, individual organoids were dissociated using the reagents from the Neural Tissue Dissociation Kit (P) (Miltenyi Biotec, #130-092-628).

Each organoid was transferred to a 1.5-mL Eppendorf tube, washed once with 1 mL PBS, and then with 1 mL of provided Buffer X. Subsequently, 25 µL of provided Enzyme P solution was diluted in 1 mL Buffer X and added to the organoids. Organoids were then incubated in the enzyme solution at 37°C with agitation at 900 rpm for 25 min. During this incubation period, the organoids were pipetted up and down using a 1-mL pipette every 5 min to assist dissociation. Then 5 µL of Enzyme A together with 10 µL of Buffer Y were added to the partially dissociated organoids, followed by a 15-min incubation at 37°C without shaking. Thereafter the cells from the fully dissociated organoids were handled on ice. The dissociated cells were centrifuged at 300 x g for 5 min at 4°C to pellet the cells and remove residual enzyme solution. The cell pellet was then resuspended in 250 µL PBS and passed through a 70-µm filter (pluriSelect, 43-10070). Prior to FACS-sorting, Hoechst 33342 (Thermo Fisher, 62249) was added to the cell suspension at a 1:10,000 dilution. This was done to allow exclusion of debris from nucleated cells during FACS.

FACS sorting was performed using a FACSAria (BD Biosciences). Up to 500 fluorescent cones and rods were sorted directly into guanidine lysis buffer (0.25 M GuHCl, 24 μM dNTPs, 1.8 μM oligo-dT, 1.2 μM DTT, 1 M Betaine) for subsequent RNA extraction and immediately frozen at - 70 °C for storage.

### Bulk RNA-sequencing

Cell lysates were processed following the bulk FLASH-seq protocol^82,83^. Briefly, RNA was converted to cDNA fragments using Superscript IV (Thermo Fisher Scientific, #18090200) and amplified with KAPA HiFi HotStart (Roche, #KK2602,). The cDNA was then cleaned using a 0.8x ratio of homebrew SeraMag beads in 18% PEG (CytiviaTm, #GE24152105050250). cDNA concentration and quality were measured using Qubit (Thermo Fisher Scientific, #Q33231) and an Agilent Bioanalyzer (Agilent, #5067-4626). The cDNA was normalized to 200 pg/μL before tagmentation using 0.2 μM of homemade Tn5. Tn5 transposase was produced by the EPFL Protein Facility (Lausanne, Switzerland). The reaction was halted with 0.2% SDS. An indexing PCR was performed to add Nextera index adapters (1 μM, Integrated DNA Technology) using KAPA HiFi reagents (Roche, #KK2102). Libraries were pooled in equal volumes and a final 0.8x cleanup performed with homebrew SeraMag beads before measuring sample concentration and quality. The library pool was normalized and sequenced on Illumina NextSeq MO flowcell (75-8-8-75) at approximately 1 million reads/sample. Basecalling and demultiplexing were performed with bcl2fastq (v2.20, Illumina Inc.).

### Kinase profiling

CS-KI-1, CS-KI-1A, CS-KI-2, and CS-KI-2A were subjected to *in vitro* kinase profiling against a panel of 350 human wild-type kinases. This profiling was conducted by the Reaction Biology Corporation using their PanQinase assay method, a plate-based assay that measures kinase activity through the transfer of radioactively labeled ATP onto a specific substrate. An analogues assay has been described before^84^. Briefly, kinase/substrate pairs and cofactors were prepared in a specific buffer, introducing the compounds at a concentration of 10 μM, followed by the addition of a mix of ATP and 33P ATP after approximately 20 min. For detailed assay conditions, refer to Supplementary Table 5. The reactions, maintained at 25 °C for two hours, were then applied to P81 ion exchange filter papers. Subsequent washing removed unbound phosphate and kinase activity was quantified by comparing the activity in test samples against vehicle (DMSO) controls, adjusted for background from inactive enzyme controls. Detailed experimental conditions are provided in Supplementary Table 7.

### Quantification and statistical analysis

All quantifications, statistical analyses, and plots were executed using R, Python, ImageJ or GraphpadPrism. All illustrations were created using Adobe Illustrator, while chemical structures were rendered with ChemDraw.

If not stated otherwise, ‘n’ always refers to the number of organoids per condition. The p-values depicted in the figures are not corrected for multiple testing If not indicated otherwise. A summary of the primary screen dataset can be found at https://ConeTargetedCompoundScreen.iob.ch.

### Promoter specificity and efficacy analysis

Promoter specificity and efficacy quantification was performed using ImageJ. Maximum intensity projections were calculated and cells were then manually counted using the ImageJ plugin, Cell Counter.

For the ProA7-GFP construct (Figure 1E), five different organoids were analyzed. The specificity was determined by calculating the percentage of all GFP-positive cells that were also ARR3-positive. Efficacy was determined by calculating the percentage of all ARR3-positive cells that were also GFP-positive.

For the ProA330-GFP construct (Figure 5B), three different organoids were analyzed using the methods used for ProA7-GFP. Specificity was determined by calculating the ratio of all GFP-positive cells located in the outer nuclear layer that were not ARR3-positive. Efficacy was determined by calculating the percentage of GFP-positive and ARR3-negative cells among all cells of the outer nuclear layer counted by Hoechst staining. The specificity and efficacy in mice were evaluated in a similar manner, using three retinas from two mice (Figure S10). Specificities and efficacies in the results section are displayed as mean ± sd.

### Cell counting algorithms

To assess cone survival in organoids, an algorithm was designed that locates and counts local maxima in pixel intensity values corresponding to GFP-expressing cells. Three distinct counting approaches were employed: counting was done image-by-image from a 3D confocal stack (referred to as ‘3D-additive-count’), from the entire 3D stack (referred to as ‘3D-count’), and from the maximum intensity projection of the 3D stack (referred to as ‘MIP-count’).

Initially, Gaussian filtering was applied to each image to minimize background noise. Following this, local maxima in pixel intensity were identified within each image using the peak_local_max function from the skicit-image package in Python. Any detected local maxima that fell below 1.25 times the frame’s average pixel value were disregarded. This was performed in 3D for the 3D-count. These detected local maxima were then subjected to a three-step filtering process to ensure they accurately represented cone cells.

In the first step of filtering, local maxima of low contrast were removed by applying Otsu thresholding to a local Region of Interest (ROI) around the local maximum. If active pixels were detected at the ROI edges, the window size was expanded. This iterative process continued until only inactive pixels were found at the ROI edges. Local maxima corresponding to ROIs exceeding a size of 70 x 70 pixels (113.75 x 113.75 µm) were excluded.

The second filtering step aimed to separate objects that were closely situated. Objects with a diameter ranging from 8.1 to 65 µm and with a perimeter-to-area ratio between 4 and 6.5 were selected for a process known as binary erosion, which effectively separated such adjacent or touching objects.

In the final filtering step, attributes like object diameter, perimeter-to-area ratio, and the contrast between object foreground and background were analyzed. Only objects with diameters between 6 and 100 µm and a perimeter-to-area ratio between 0.1 and 4 were retained. Low-contrast objects, defined as those for which the foreground was no more than 1.2 times brighter than the background, were also excluded.

All filtering steps for the 3D-count were done in 2D on the z-plane where each local maximum was identified.

In some experiments where noise levels were high, cell candidates where all pixels were below 150 were excluded (Figures 1H, 4G-4I, 7C and S1). The cell counts obtained were normalized to the initial cell count yielding relative cone survival values.

For counting rod photoreceptors (Figures 5D-5F and 7D), slight modifications were made to the parameters of the cone-counting algorithm. For rod photoreceptors, the maximum intensity projections were quantified. The image resolution was enhanced fourfold via cubic interpolation. The minimum allowable diameter for cell candidates was also reduced from 4 µm to 2 µm, and any candidates where all pixels were below an intensity of 200 were excluded.

If not stated otherwise cone-survival was calculated using the counts from the 3D-additive-count algorithm.

### Target categorization

Categorizer software^85^ was used to categorize the targets of the MOA compound library. Target categories were assigned to their respective compounds. If a compound had multiple targets, the most common category found among the targets was assigned.

### Cell count thresholding

For the primary screen, thresholds were determined after visually inspecting images with the lowest reported D0 counts, ensuring the inclusion of as many data points as possible. These thresholds were set uniquely for each quantification algorithm (Figure S2). The threshold for the secondary screens were the same as for the primary screen (Figures S4 and S6).

Similarly, thresholding on rod data was conducted following a visual inspection of images (Figure S10).

### Adjusted cone survival

To compensate for the effect of initial cone counts on the survival of glucose deprivation, we calculated an adjusted version of the cone survival. For this, a linear model was fitted using the complete primary screen dataset to explain the cone survival with the logarithm of the scaled cone counts at D0. The obtained regression coefficient was then multiplied with the scaled logarithm of the D0 cone counts. Subtracting this term from the original cone survival yielded the adjusted values. This was done separately for all three cone counting algorithms (Figures 2E and S2).

This adjustment sometimes led to values higher than 100% and very rarely to values lower than 0%.

Since the cone-damaging secondary screen was done in normal glucose, we analyzed separately the relationship between cone survival and D0 counts in a newly calculated linear model. While we found a linear model that significantly explains this relationship, it only accounted for a marginal amount of the explained variation for all three quantification algorithms (n=711, R^2^=0.007-0.018, p-values<0.001, Figure S6). Therefore, we did not generate adjusted cone survival values for this dataset.

For the cone-saving secondary screen dataset, we used the linear model originally generated from the primary screen to compute the adjusted values. This primary screen model accurately predicted the D0 to cone survival relationship in the secondary screen (n=673-681, R^2^=0.17-0.22, p<0.001, Figure S6).

### Well position bias analysis

To account for any potential influence on the results of well position within the 96-well plate, an analysis was performed on the mean adjusted cone survival for each well position, based on the primary screen dataset with the 3D-additive-count. This procedure assumes that most of the compounds under study do not exert a significant effect on cone survival. These mean values were then compared. If the differences between the means were found to fall within the range of the minimum standard deviation observed for the least variable well, it was determined that the well position did not have a significant impact on the outcome. Excluding the positions of the normal glucose control wells, cone survival was not influenced by any of the well positions (Figure S2).

### Analysis of primary screen data

To compare cone survival across various compound conditions while controlling for the initial count at D0, an Analysis of Covariance (ANCOVA) was conducted. In this analysis, the dependent variable was the unadjusted cone survival, and the independent variable was the compound condition (low glucose control vs. compound 1 vs. compound 2, etc.). The raw count at D0 served as the covariate in the model. P-values were calculated for a two-sided test comparing each compound to the low glucose controls. To identify significant hits from the primary screen, a statistical threshold of p<0.05 was set. The Benjamini-Hochberg correction was employed to account for multiple testing. Consequently, only compounds with an adjusted p-value less than 0.05 were considered significant. The compounds selected for secondary screening were determined based on their p-values and after visual inspection of images.

### Mode of action names

The mode of action for each compound was sourced from Canham et al.^57^. For compounds subjected to secondary screens, a concise version of their modes of action was manually generated (Figures 3, 4, S4-S6 and S7).

### Analysis of the secondary screen cone-damaging dataset

To assess the impact of different conditions compared to the control, an Analysis of Variance (ANOVA) was conducted that compared all different compound conditions to the normal glucose control. The resulting p-values were subsequently adjusted for multiple comparisons using Benjamini-Hochberg correction for all compounds and concentrations. P-values were calculated for a one-sided test. Compounds and concentrations with adjusted p-values less than 0.05 were deemed statistically significant. Controls are also displayed in Figure S4.

### Clustering cone-damaging compounds from the secondary screen

To categorize compounds from the secondary screen based on cone survivals at four different concentrations, hierarchical clustering was performed using medians of the cone survival values of all significant cone-damaging compounds. The Elbow Method was employed to identify the optimal number of clusters, involving a plot of the total within-cluster sum of squares (WSS) against the number of potential clusters. WSS values were computed for all possible solutions, ranging from 1 to 34 clusters, and an elbow in the curve was observed at four clusters. The resulting clusters were then assigned to all compounds, as depicted in Figure 3C. One outlier was removed from cluster 3 and one from cluster 4 in Figure 3C.

### Definition of target classes

HDAC1, HDAC2, HDAC3, HDAC4, HDAC5, HDAC6, HDAC7, HDAC8, HDAC9, HDAC10 and HDAC11 were categorized as HDAC I/IIs. SIRT1, SIRT2, SIRT3 and SIRT 6 were categorized as HDAC IIIs. TUBA1A, TUBA1B, TUBA1C, TUBA3C, TUBA3D, TUBA3E, TUBA4A, TUBA8, TUBB, TUBB1, TUBB2A, TUBB2B, TUBB3, TUBB4A, TUBB4B, TUBB6, TUBB8, TUBD1, TUBG1, and TUBG2 were categorized as tubulins. These target classes were used in Figures 3E, 3F and S4.

### Analysis of the secondary screen cone-saving dataset

The secondary screen cone-saving dataset was analyzed in the same way as the primary dataset using an ANCOVA with the D0 count as covariate, comparing compound effects to the low glucose controls. The resulting p-values were subsequently adjusted using a Benjamini-Hochberg correction for multiple testing for all compounds and concentrations (Figure S6). The p-values were calculated for a one-sided test. Controls are shown in Figure S6.

### Analysis of target effects in the primary screen dataset

In the analysis of the impact of compound targets on cone survival, targets with three or more listed compounds were initially selected (Figure 2C). For each of these targets, the average of the median adjusted cone survival across all targeting compounds was determined. This average was then compared to a distribution generated by randomly drawing an equal number of compounds and calculating their mean of the median adjusted cone survival.

This process of random drawing was performed 10,000 times initially to create a distribution of mean values. In the analysis of cone-saving targets, the number of these randomly generated means that were higher than the observed mean was determined (Figure 4E and S8). Conversely, for the cone-damaging targets, the number that were lower was determined (Figure 3F).

If less than 10 of the random means were found to be higher (or lower, depending on the analysis), the process was repeated with 100,000 random draws to ensure robustness. The p-value for each target was then estimated as the fraction of random means that were found to be higher (or lower) than the observed mean, plus one, divided by the total number of random draws.

Finally, to account for multiple comparisons, these p-values were corrected using the Benjamini-Hochberg method and a significance threshold set at 0.05. However, in Figures 3F, 4E and S6 the unadjusted p-values are displayed.

### Analysis of IC50

IC50 values for all compounds in the library targeting HSP90AA1, HSP90AB1, MTOR, PIK3CA, PIK3CB, PIK3CD, PDGFRA, and PDGFRB, were sourced from the ChEMBL database^86^. This dataset encompassed reported IC50s, even for compound-target pairs not present in the MOA library. In instances where multiple IC50 values were noted for a specific compound-target combination, the median of these values was used for subsequent analysis. Spearman correlation coefficients were determined by correlating the median-adjusted cone survival with median IC50 values.

### Analysis of transcriptomes

Sequencing reads were processed into gene counts using Snakemake (v7.21.0), a workflow management system. The workflow consisted of two main steps: read alignment and differential gene expression analysis. Reads were aligned against the GRCh38 (Ensembl release 109) reference genome using STAR (v2.7.10b). Both the number of reads per gene (--quantMode GeneCounts) and alignments translated into transcripts coordinates (--quantMode TranscriptomeSAM) were set as outputs. The reference genome was augmented to include sequences from two transgenes (ProA7-GFP and ProA330-tdTomato) used in cell sorting. Read counts per gene and per sample were aggregated using custom Python scripts, and genes expressed in fewer than 5% of the samples were filtered out. DESeq2 (v1.38) was then employed to identify differentially expressed genes, using a log_2_ fold change threshold of 1 and a 5% significance level after Benjamini-Hochberg correction for multiple hypothesis testing. Principal component analysis was performed using scikit-learn (v.1.2.2) on normalized and standardized gene counts. GSEApy pre-rank was employed to perform gene set enrichment analysis using gene ontology terms (biological process, molecular function and cellular component ontologies) as well as hallmark gene sets from the Human Molecular Signatures Database (MSigDB). For this purpose, genes were ranked based on Wald test statistics provided by DESeq2. Transcripts from ribosomal-protein coding genes were excluded from this analysis, as these genes can be variable or highly expressed irrespective of the tested conditions. Data were normalized to transcript counts per 10,000 adjusted for non-overlapping exon lengths (TP10k), where lengths were estimated using the R package GenomicFeatures (v1.50.2). Marker genes were identified based on an available adult human peripheral retina atlas (https://data.iob.ch) using scanpy’s rank_genes_groups function.

### Analysis of kinase profiling

Kinase profiling data and the main analysis were provided by Reaction Biology Corporation. Gene expression values were derived from the average gene expression in low glucose cones.

## SUPPLEMENTARY FIGURES AND SUPPLEMENTARY FIGURE LEGENDS

**Figure S1:**
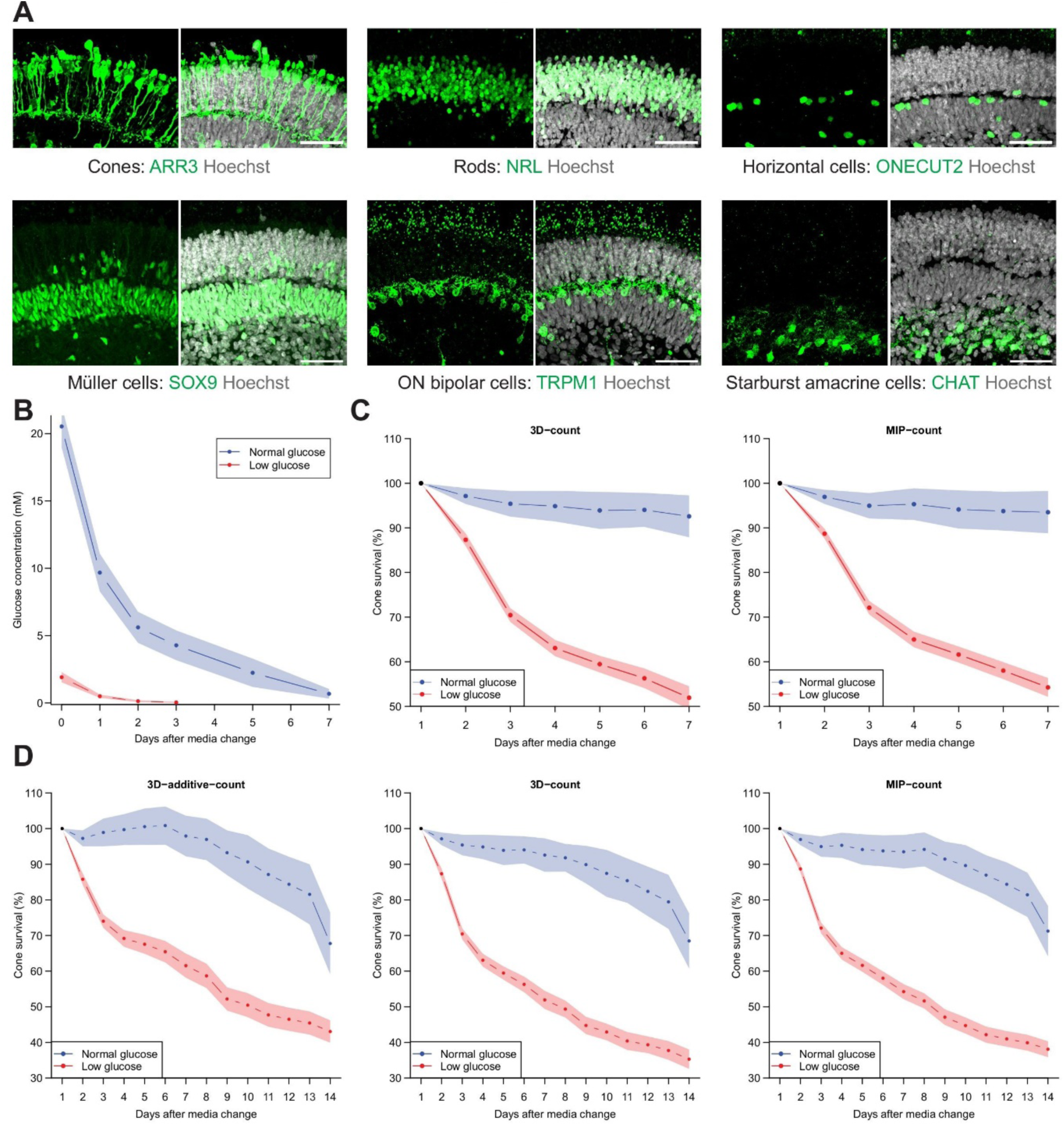
Cell types of mature human retinal organoids and glucose starvation. **A:** Marker gene expression of cell types in human retinal organoids. Confocal images of sectioned and stained human retinal organoids at week 30 of maturation with indicated staining (scale bar, 25 µm). **B:** Glucose consumption assay of human retinal organoids. Mean ± se from 10 human retinal organoids. **C:** Quantification of cone survival in human retinal organoids in normal and low glucose over seven days. Quantification with either 3D-count (left) or MIP-count (right). Mean ± se. Low glucose, n=26; normal glucose, n=6. **D:** Same as (C) but for 14 days using all three quantification algorithms (3D-additive-count, 3D-count, and MIP-count).

**Figure S2:**
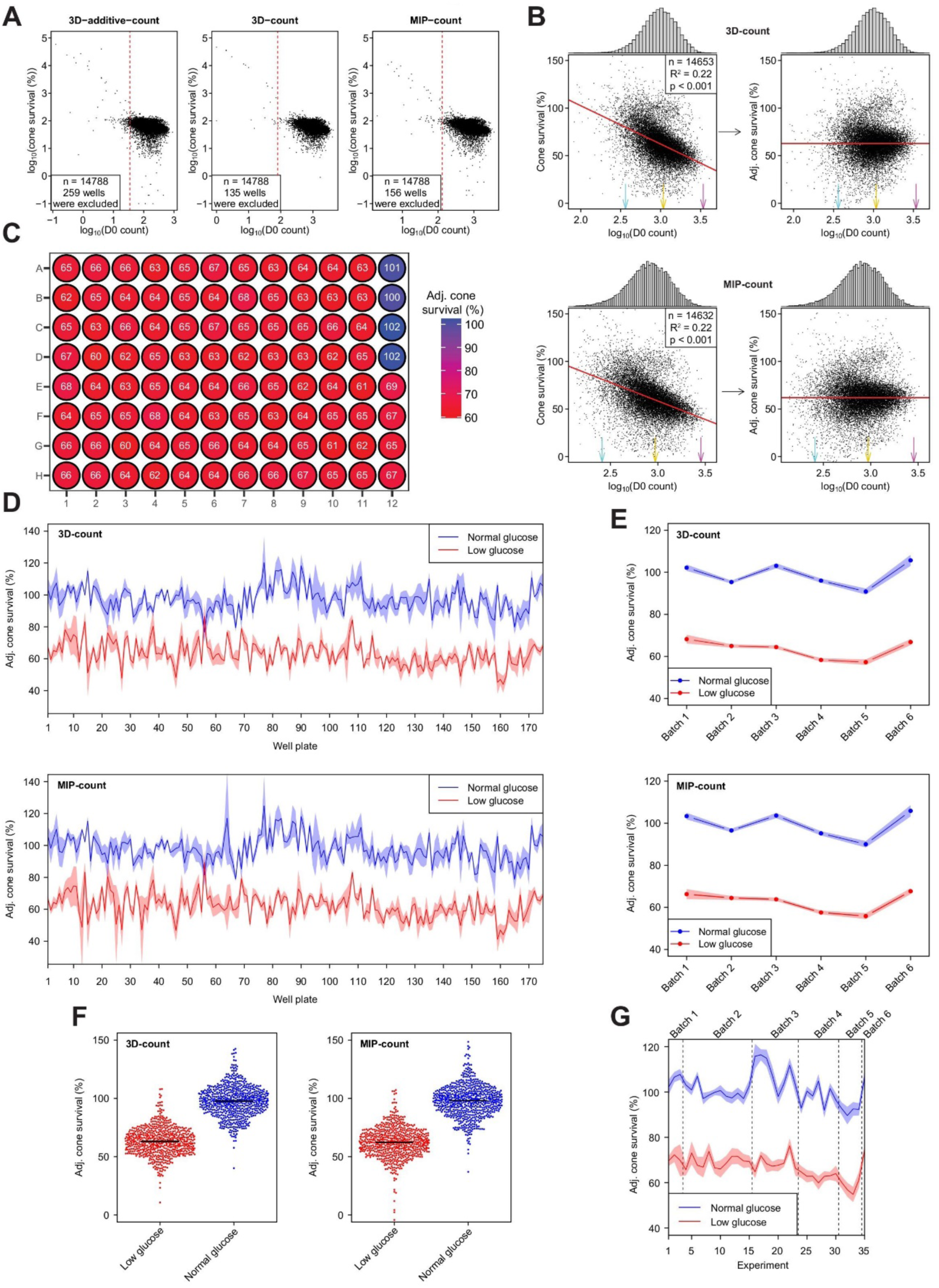
Primary screen: analysis. **A:** Data thresholding based on counts at D0 for all quantification algorithms. The dotted red line indicates the threshold beyond which wells are included in the analysis. **B:** Adjustment of cone survival values based on the D0 count. Top: 3D-count. Bottom: MIP-count. Left: cone survival and the logarithm of the cone count at D0, fitted with a linear model. Regression line, red; summary statistics, top right corner of each panel. Right: adjusted cone survival and the logarithm of the cone count at D0 with the transformed regression line. Colored arrows correspond to D0 counts of example images in Figure 2E. **C:** Well position bias analysis. The means of adjusted cone survival for all wells of the primary screen data are depicted. **D-E**: Adjusted cone survival of normal and low glucose controls for individual well plates (D) and organoid batches (E), mean ± se. Top: 3D-count. Bottom: MIP-count. **F:** Adjusted cone survival of all normal and low glucose controls. Left: 3D-count. Right: MIP-count. **G:** Adjusted cone survival of normal and low glucose controls for individual experiments (groups of 5 well plates containing all replicates for compounds) using the 3D-additive-count algorithm, mean ± se.

**Figure S3:**
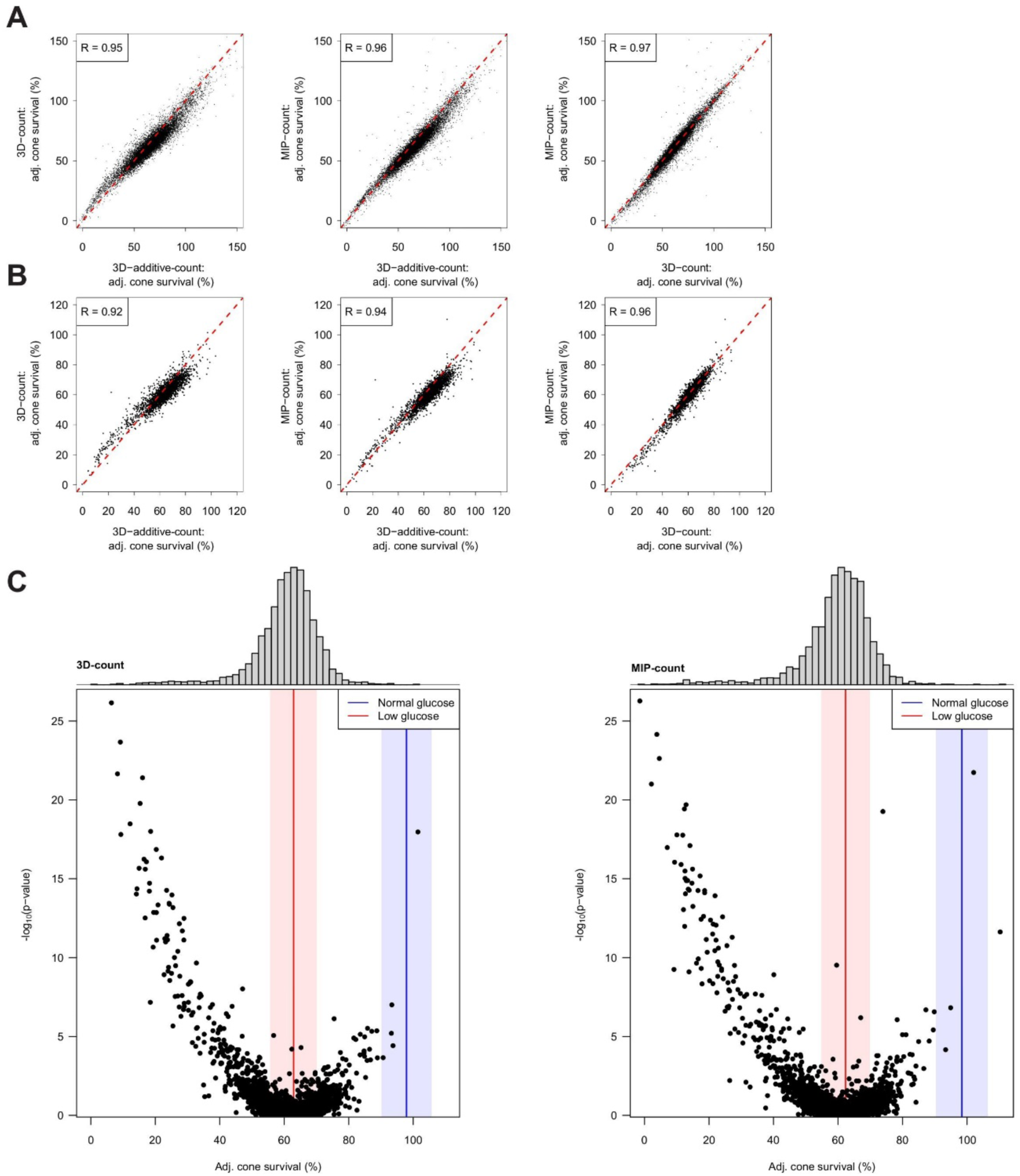
Primary screen: comparison of different quantifications. **A-B:** Comparison of different quantification methods. Comparing either adjusted cone survival values of individual organoids (A) or the median of adjusted cone survival values for individual compounds (B). The dotted red line represents the unity line and ‘R’ denotes the Pearson’s correlation coefficient. **C:** Effect of compounds on cone survival in the primary screen. Left: 3D-count. Right: MIP-count. Each dot corresponds to the effect of one compound, with the median of the adjusted cone survival of five human retinal organoids on the x-axis and the p-value comparing the cone survival in the compound and in low glucose on the y-axis. The median (line) and the interquartile range (shaded area) of normal (blue) and low (red) glucose controls are indicated. Top: the distribution of median adjusted cone survival for the compounds.

**Figure S4:**
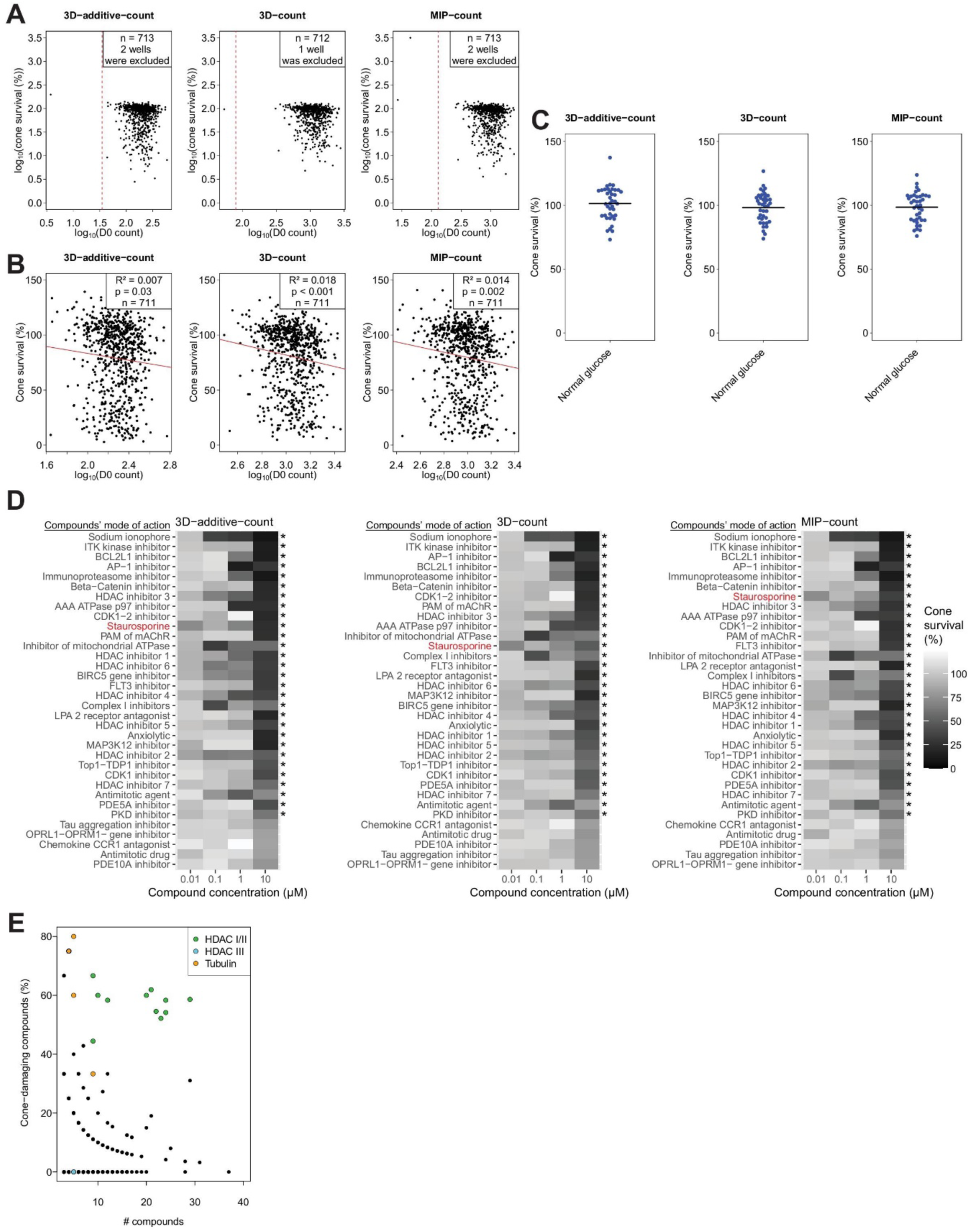
Secondary screen for cone-damaging compounds: analysis. **A:** Thresholding of data for the secondary screen of cone-damaging compounds based on counts at D0 for all quantification methods. The dotted red line indicates the threshold beyond which wells are included in the analysis. **B:** Effect of D0 count. Cone survival and the logarithm of the cone count at D0, fitted with a linear model. The regression line is shown in red and the summary statistics are displayed at top right. **C:** Normal glucose controls with indicated means for all three quantification methods. **D:** Summary of the effects of all cone-damaging compounds tested in the secondary screen. The positive control Staurosporine is indicated in red. Compounds are sorted by the minimum p-value across all concentrations, with the smallest p-value at the top. Significant compounds (at any concentration) are marked with *, denoting a p-value<0.05 (after Benjamini Hochberg correction for multiple testing). **E:** Target analysis showing the number of compounds against the fraction of compounds having a significant cone-damaging effect (p-values<0.05, after Benjamini Hochberg correction for multiple testing) for all targets with more than two compounds.

**Figure S5:**
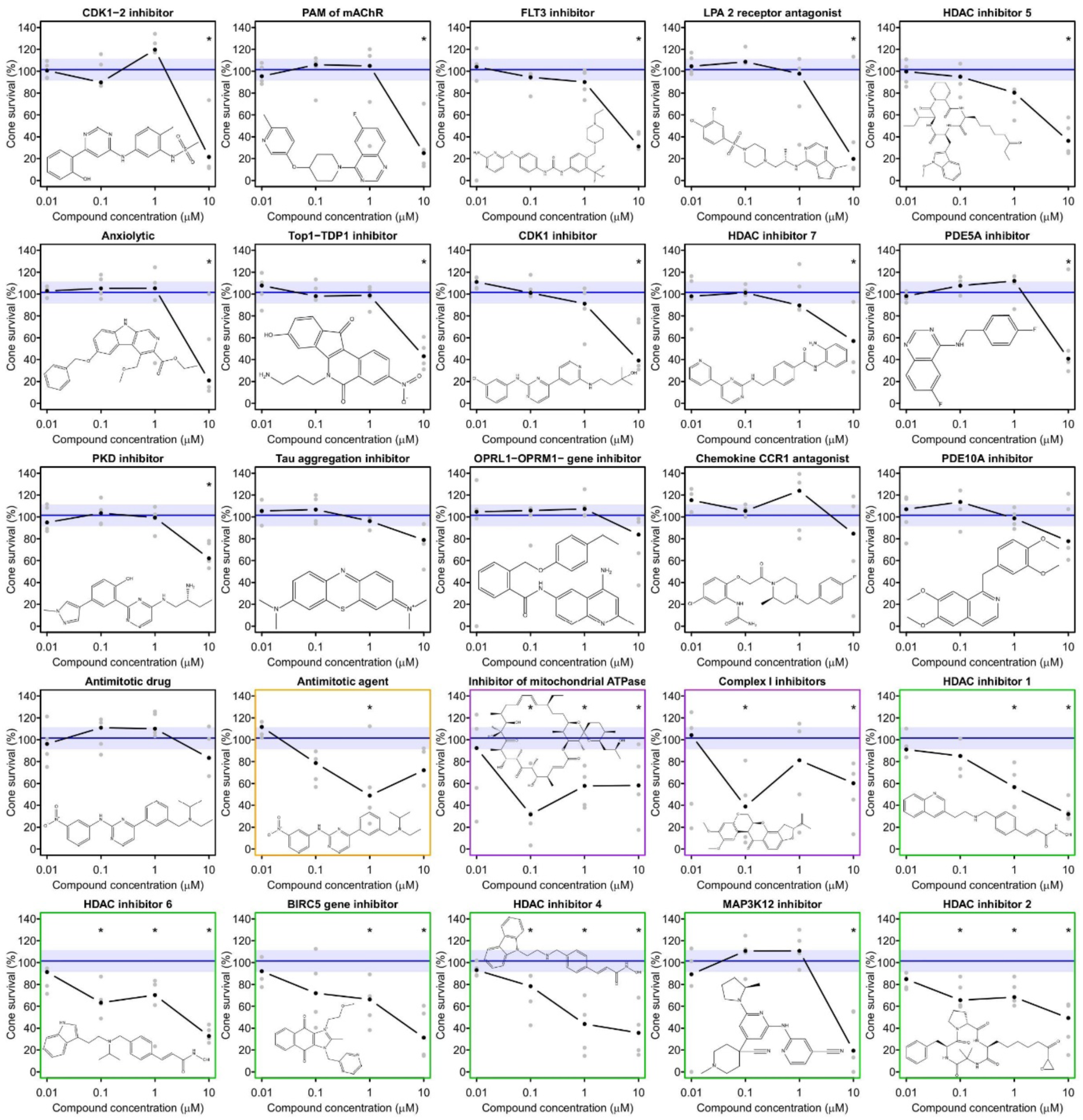
Secondary screen for cone-damaging compounds: dose-response curves. Dose-response curves of all secondary screen cone-damaging compounds, excluding those depicted in Figure 3D. The compounds are ordered based on their respective clusters from Figure 3D and minimum p-values. Black dots denote median cone survival, gray dots represent individual values. The blue line and area indicate median and interquartile range of the normal glucose controls, respectively. The colored frame corresponds to the clusters from Figure 3C. The structure of each compound is illustrated within the plot. Significant compounds and concentrations are marked with * denoting a p-value<0.05 (after Benjamini Hochberg correction for multiple testing).

**Figure S6:**
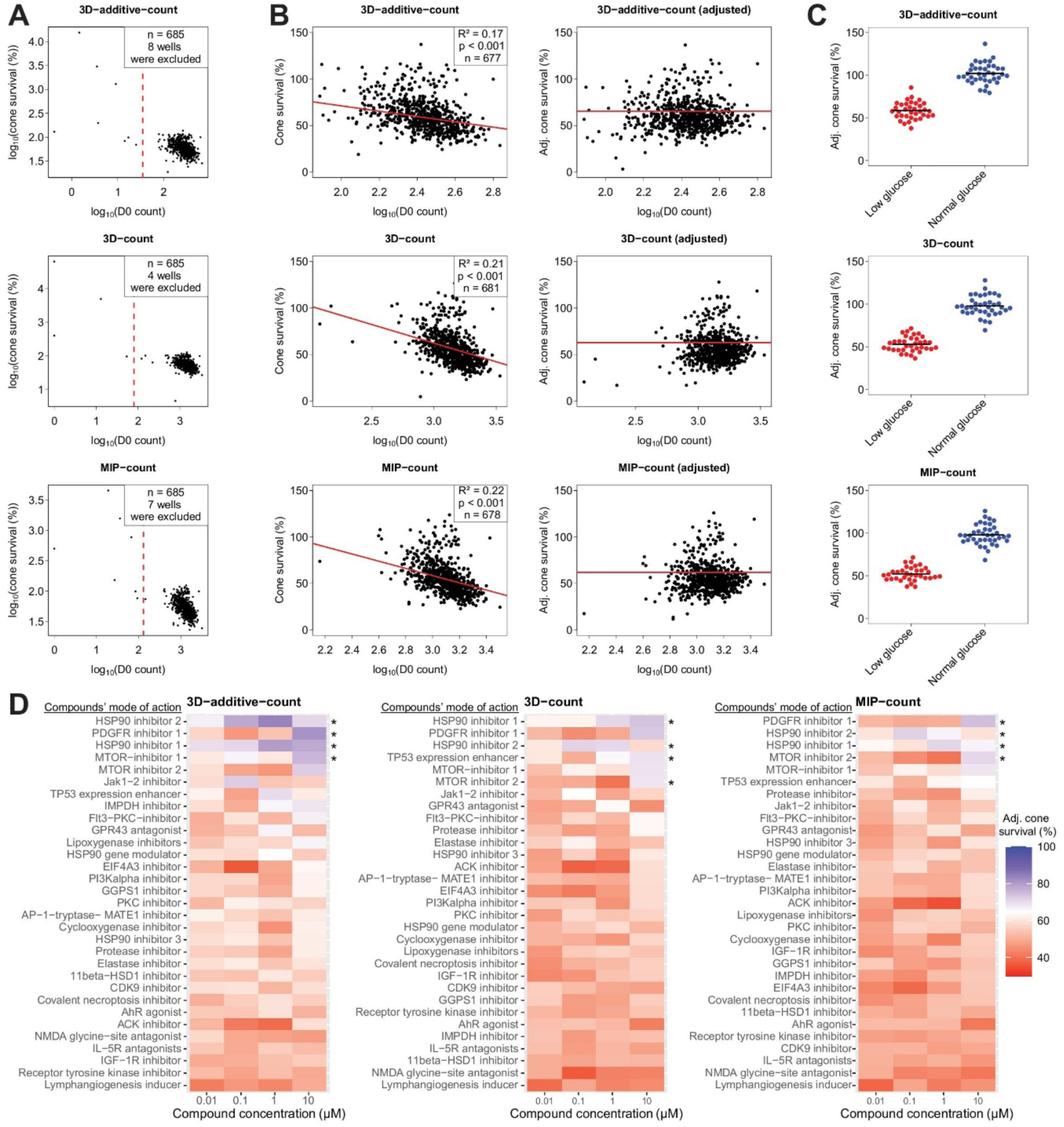
Secondary screen for cone-saving compounds: analysis. **A:** Thresholding of data for the secondary screen of cone-saving compounds based on counts at D0 for all quantification methods. The dotted red line indicates the threshold beyond which wells are included in the analysis. **B:** Adjustment of cone survival values based on D0 counts for all quantification methods. Left: cone survival and the logarithm of the cone counts at D0, fitted with a linear model. Regression line, red; summary statistics, top right corner of each panel. Right: adjusted cone survival and the logarithm of the cone counts at D0 with the transformed regression line. **C:** Adjusted cone survival of all normal and low glucose controls for all quantification methods. **D:** Summary the effects of cone-saving compounds. Compounds are sorted by the maximum median adjusted cone survival. Results are shown for all three quantification methods. Significant compounds are marked with *, denoting a p-value<0.05 (after Benjamini Hochberg correction for multiple testing).

**Figure S7:**
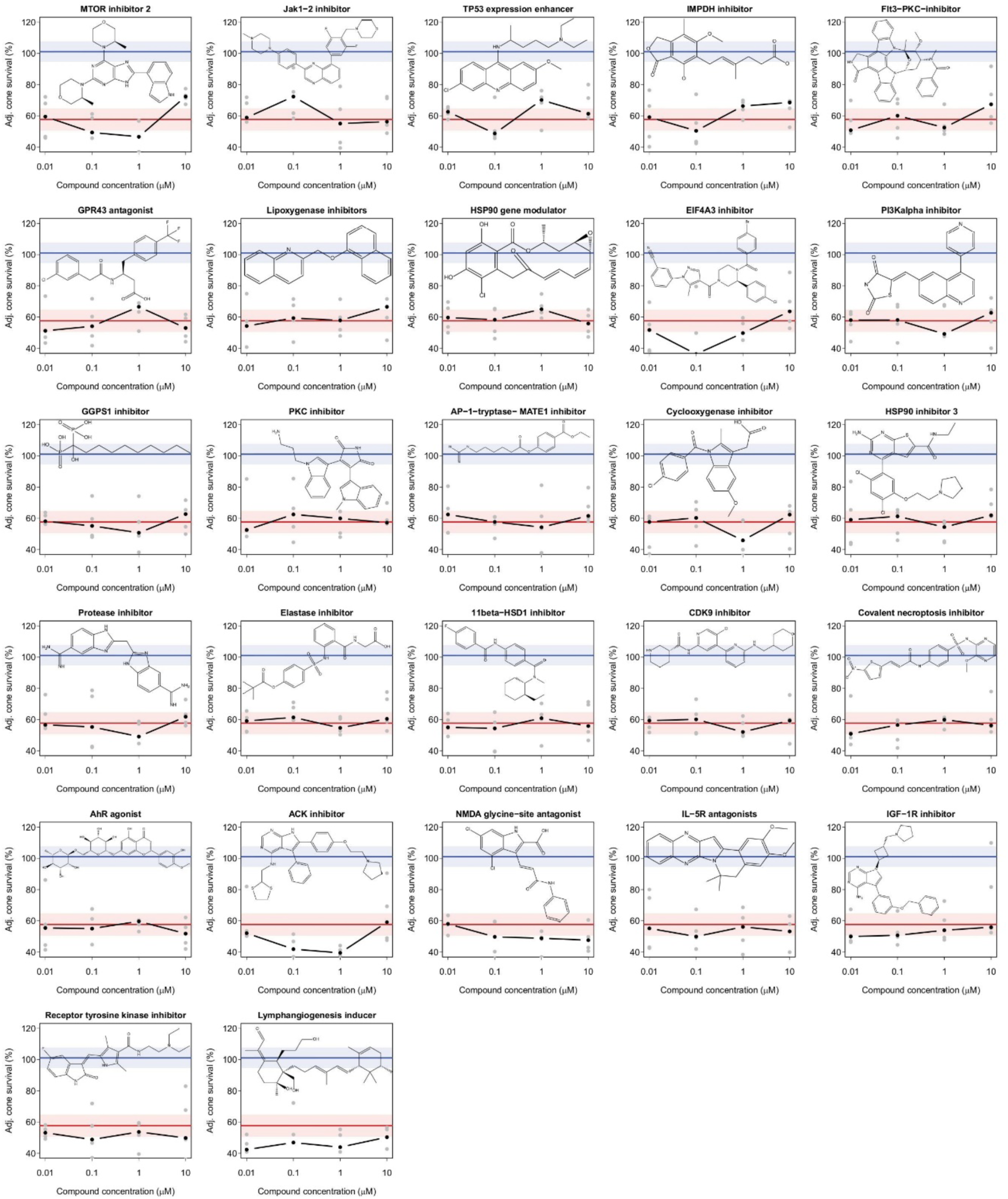
Secondary screen for cone-saving compounds: dose-response curves. Dose-response curves of all secondary screen cone-saving compounds, excluding those depicted in Figure 4B. Compounds are ordered by their maximum adjusted cone survival (most effective concentration). Black dots denote median cone survival, gray dots represent individual values. The median (line) and the interquartile range (shaded area) of normal (blue) and low (red) glucose controls are indicated. The structure of each compound is illustrated within the plot.

**Figure S8:**
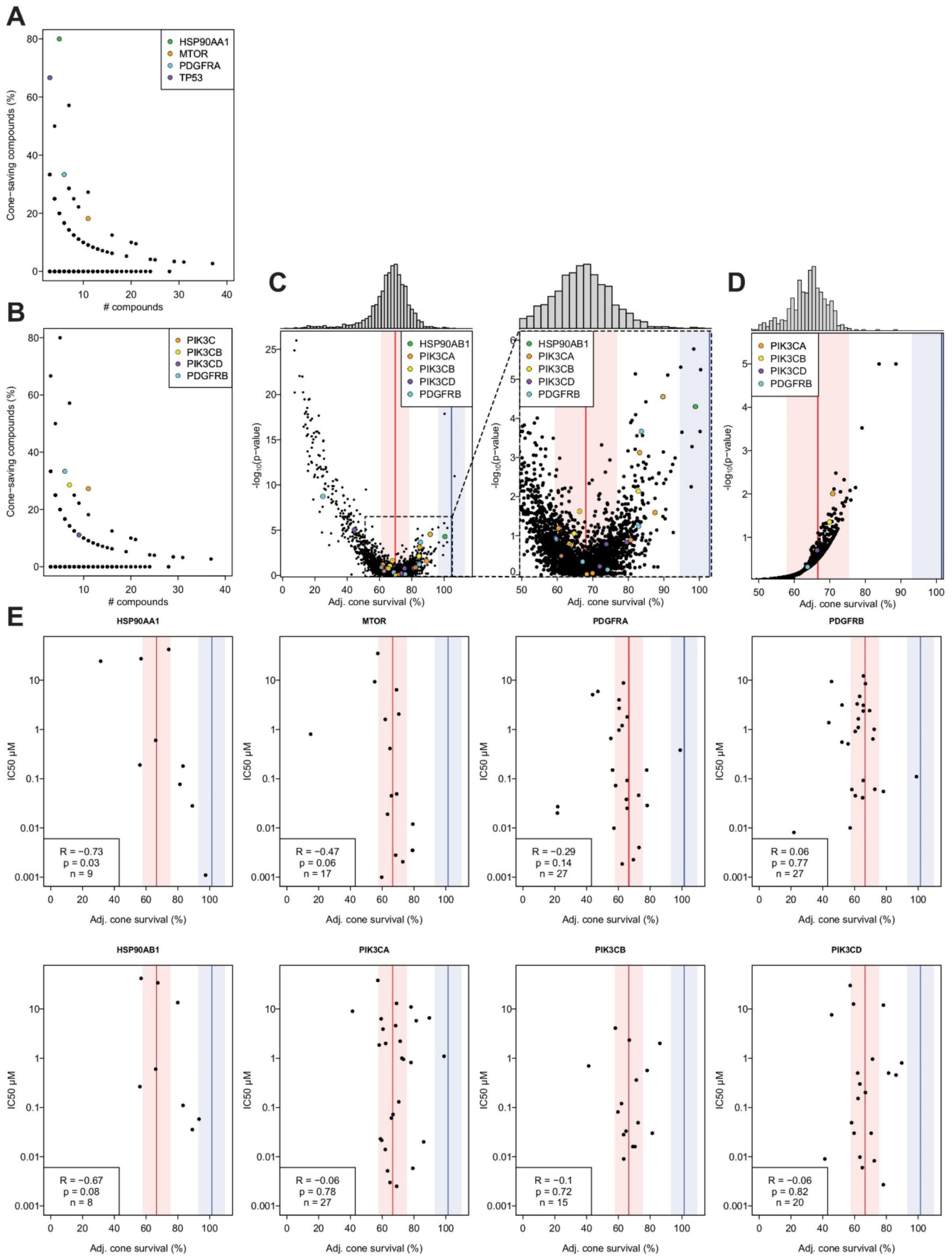
Target analysis and IC50s. **A-B**: Number of compounds and fractions of compounds having a median adjusted cone survival >80% for all targets (with more than two compounds). Indicated targets colored. **C:** Left: adjusted cone survival and p-values in the primary screen. Compound targets are indicated and colored. The median (line) and the interquartile range (shaded area) of normal (blue) and low (red) glucose controls are indicated. Right: zoom-in of plot on the left. **D:** Target analysis of primary screen. The means of the median adjusted cone survival for each target are shown, along with their p-values. Specific targets are indicated. The median and the interquartile range are labeled as in (C). **E:** Median IC50 values for compounds that bind the annotated targets (labeled on top) of cone-saving compounds and their median adjusted cone survival. The median and the interquartile range are labeled as in (C).

**Figure S9:**
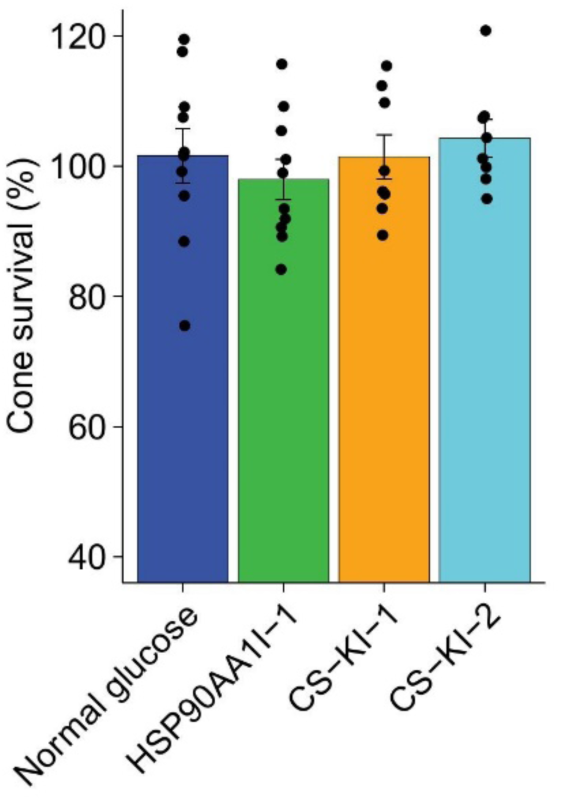
Compound effects in normal glucose. Cone survival of compound-treated organoids in normal glucose after seven days. HSP90AA1I-1 was administered at 1 µM, while CS-KI-1 and CS-KI2 were tested at 10 µM. Results are shown as mean ± se. Black dots indicate individual organoid cone survival.

**Figure S10:**
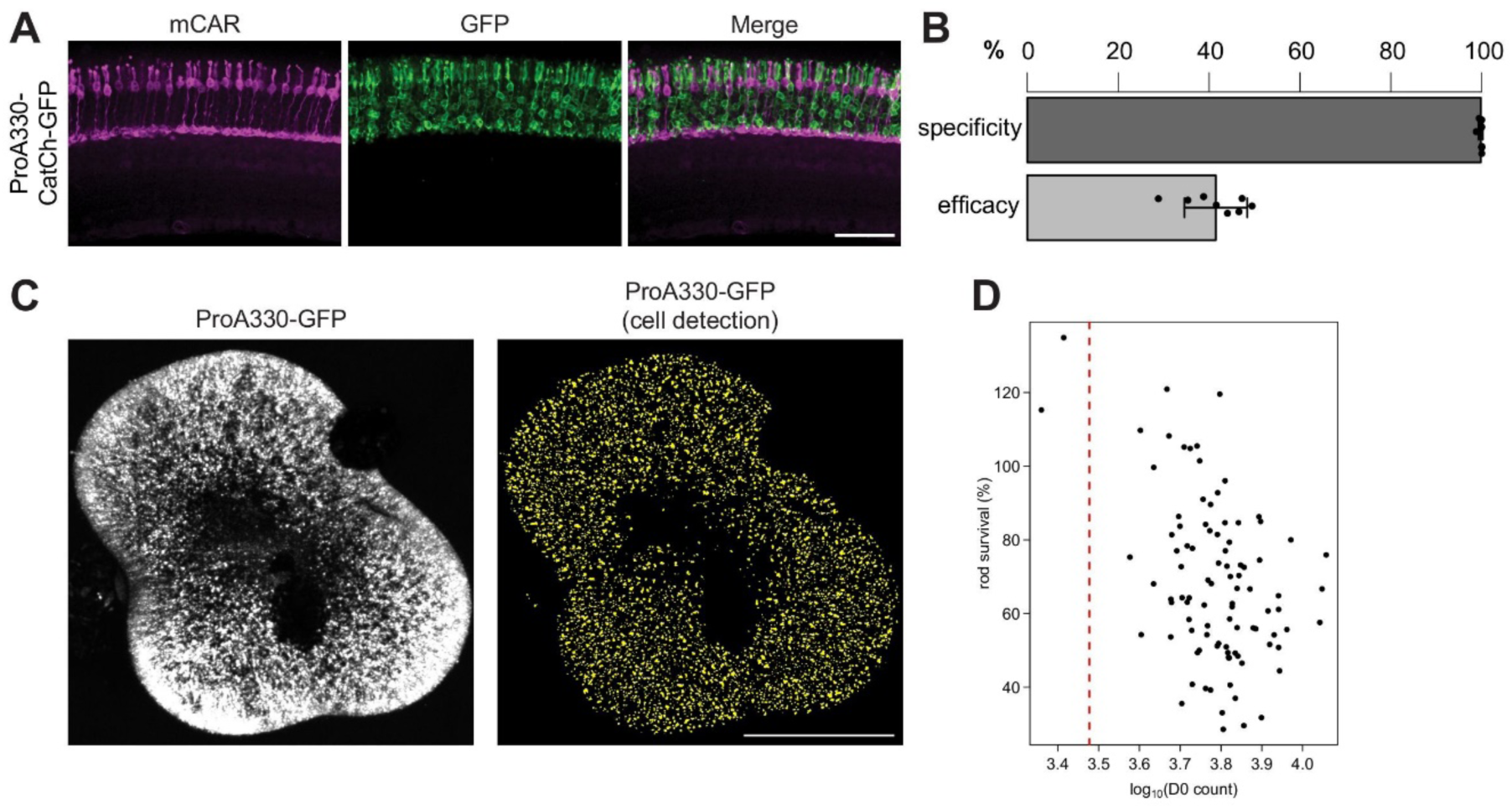
Rod promoter. **A:** Confocal image of sectioned and stained transduced mouse retina (scale bar, 50 µm). mCAR, magenta; CatCh-GFP, green. **B:** Quantification of the specificity and efficacy of rod labeling in mice by ProA330-CatCh-GFP AAV. Rods were identified as being present in the photoreceptor layer but negative for the cone-marker mCAR. Results are shown as mean ± sd. **C:** Left: example image of a ProA330-GFP AAV-transduced human retinal organoid. GFP, white. Right: Detected rods of an example human retinal organoid. Detected cells, yellow. **D:** Data thresholding based on rod counts at D0. The dotted red line indicates the threshold beyond which wells are included in the analysis.

**Figure S11:**
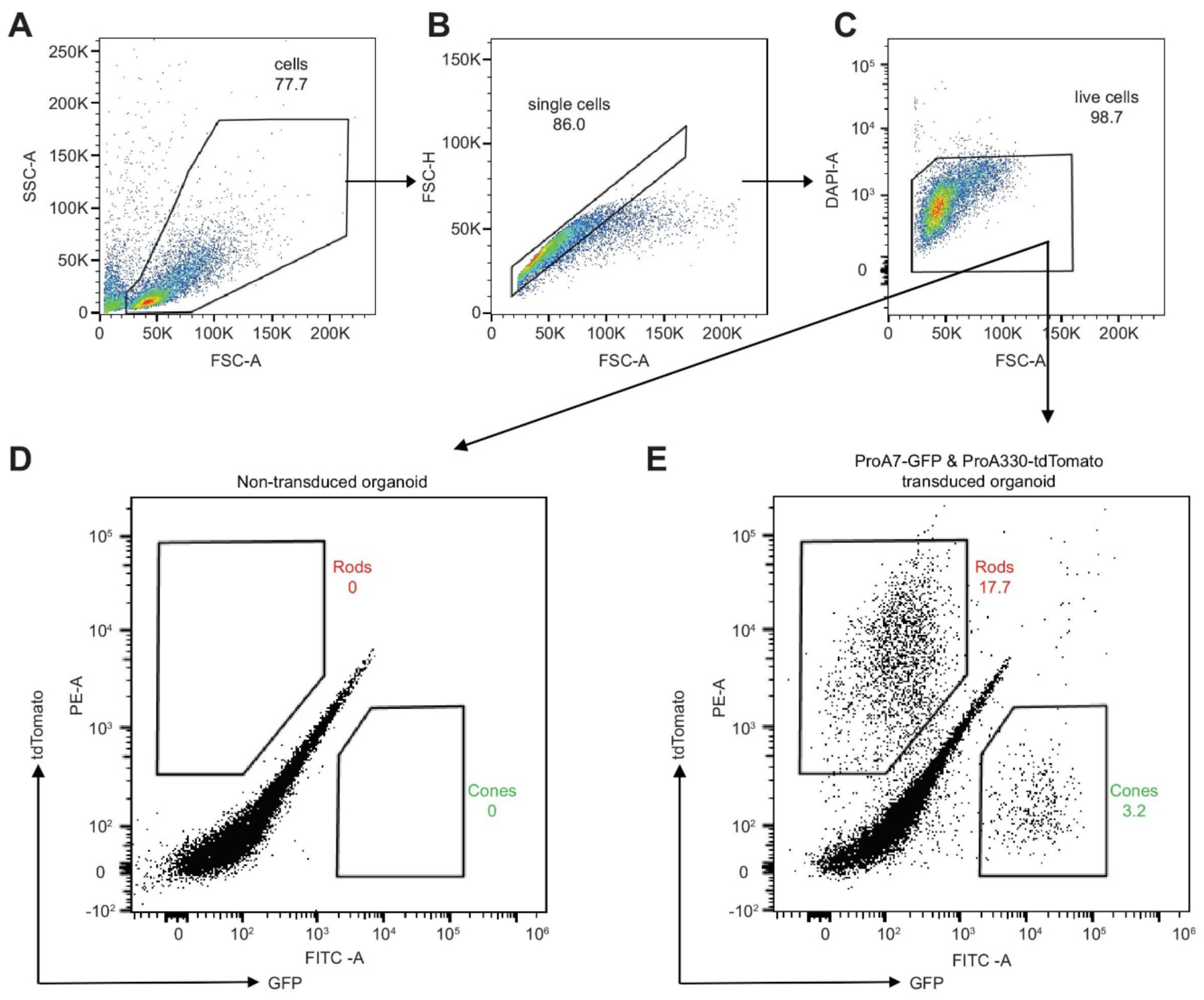
Fluorescence activated cell sorting (FACS) of photoreceptors. Representative FACS density plots from a human retinal organoid co-transduced with ProA7-GFP and ProA330-tdTomato. All numbers are percentages of gated cells. **A:** Forward scatter area (FSC-A) and side scatter area (SCS-A) to filter cells from debris. **B:** FSC-A and forward scatter height (FSC-H) to filter single cells form aggregates. **C:** FSC-A and Hoechst channel (DAPI-A) to sort out living cells. **D-E:** GFP (cone) and tdTomato (rod) positive cells from either non-transduced (D) or co-transduced (E) human retinal organoids.

**Figure S12:**
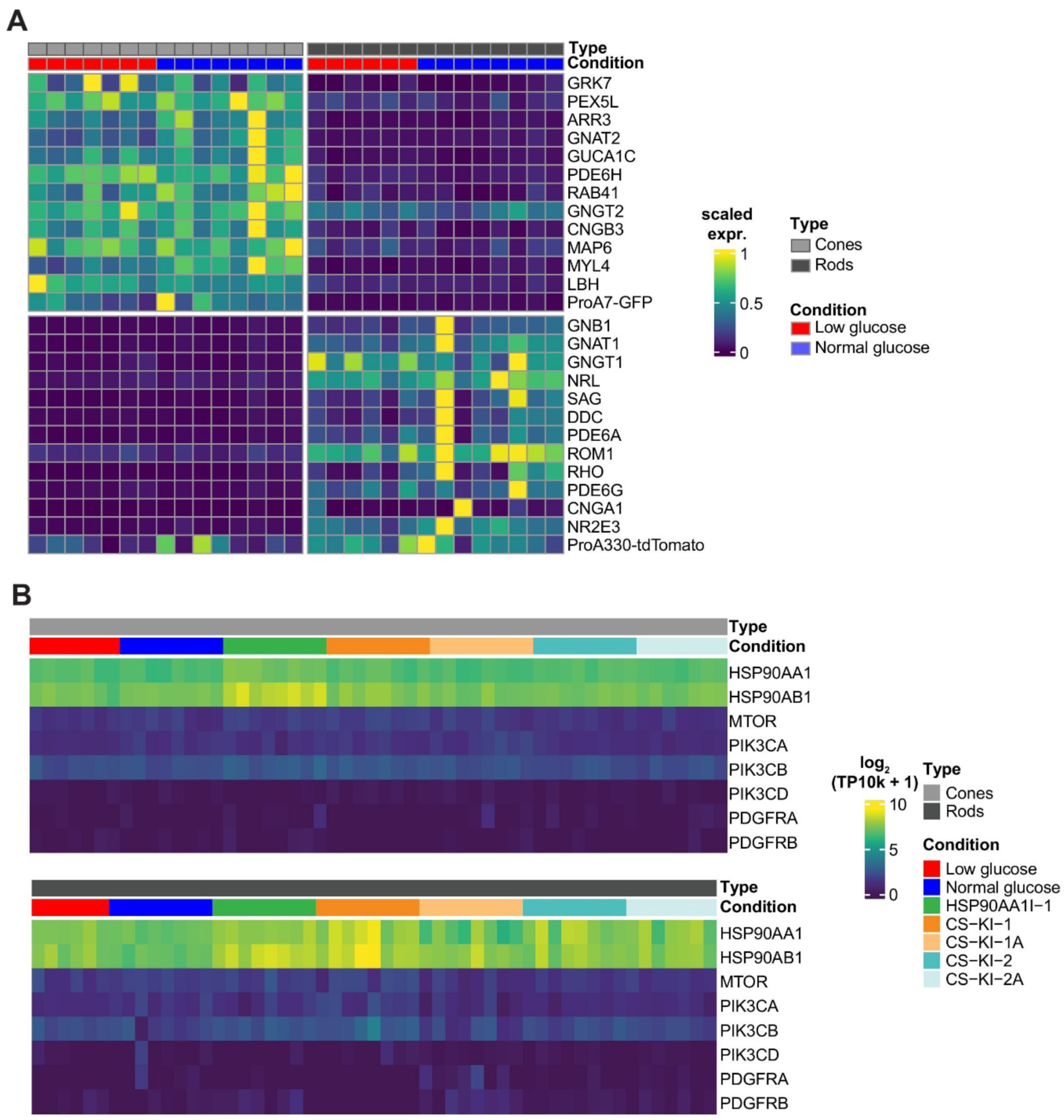
Compound target and marker gene expression. **A:** Expression of cone (top) and rod (bottom) marker genes. Heatmap colors correspond to gene expression normalized to the row-wise maximum. Normal and low glucose conditions are indicated at the top. **B**: Expression of genes encoding annotated compound targets. Colors at the top indicate cell types and conditions. Color scale indicates gene expression levels. TP10k, transcript counts per 10,000.

**Figure S13:**
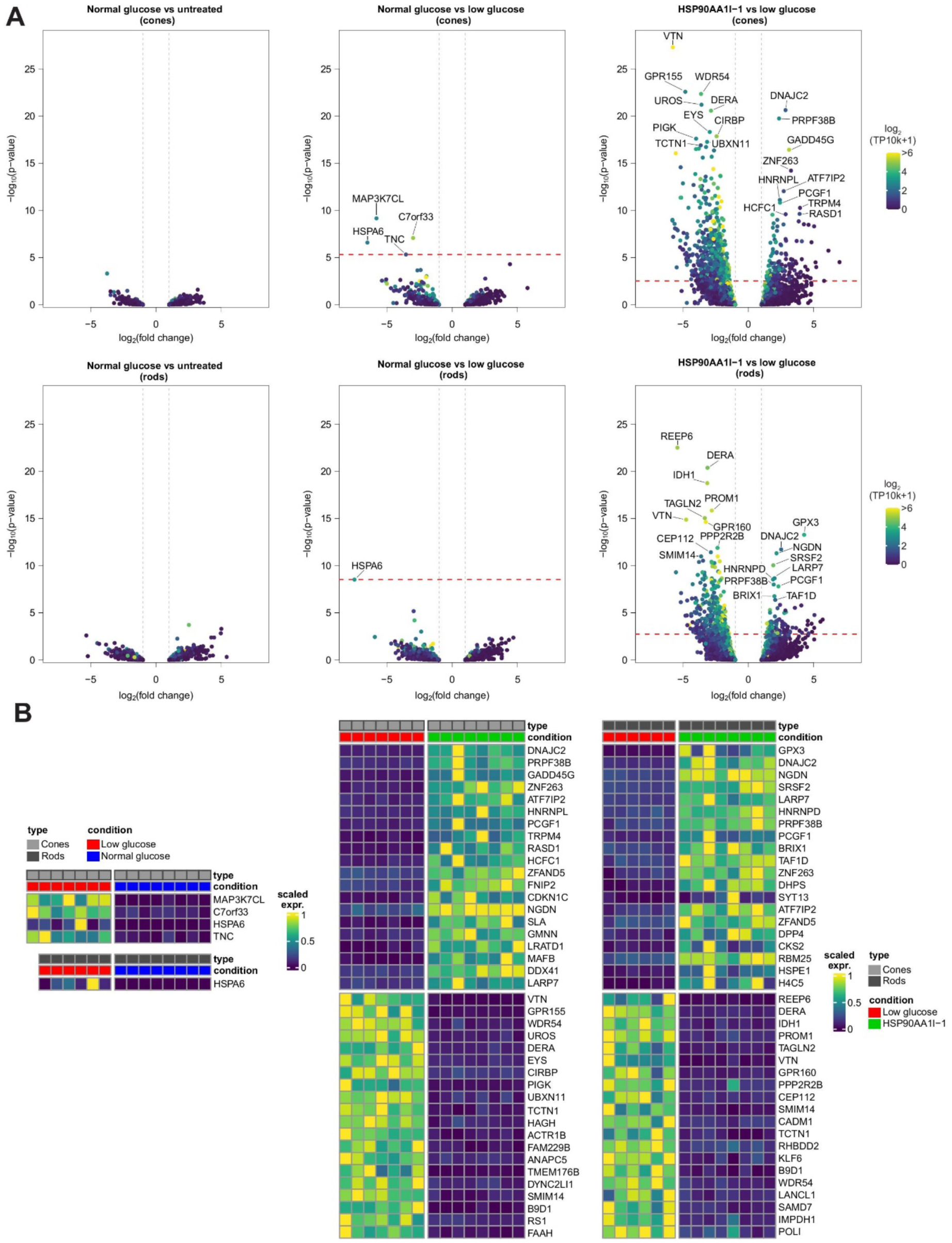
Differential gene expression analysis of controls and HSP90AA1I-1. **A:** Differential gene expression comparing human retinal organoid cone (top) and rod (bottom) transcriptomes in normal glucose with untreated controls (left), normal glucose to low glucose (middle), and HSP90AA1I-1 treatment in low glucose to low glucose (right). Gray dotted lines indicate foldchange threshold beyond which genes were included in analysis. Red dotted line indicates the p-value beyond which genes passed the significant threshold of p<0.05 after Benjamini-Hochberg correction for multiple testing. Colors indicate average gene expression in low glucose cones (top) or rods (bottom). The 10 most significantly up- or downregulated genes are labeled. TP10k, transcript counts per 10,000. **B:** Heatmaps displaying gene expression levels of differentially expressed genes across samples under specified conditions. Up to 20 up- and downregulated genes are included and ordered by p-values with the most significant p-value at the top. Heatmap colors correspond to gene expression normalized to the row-wise maximum.

**Figure S14:**
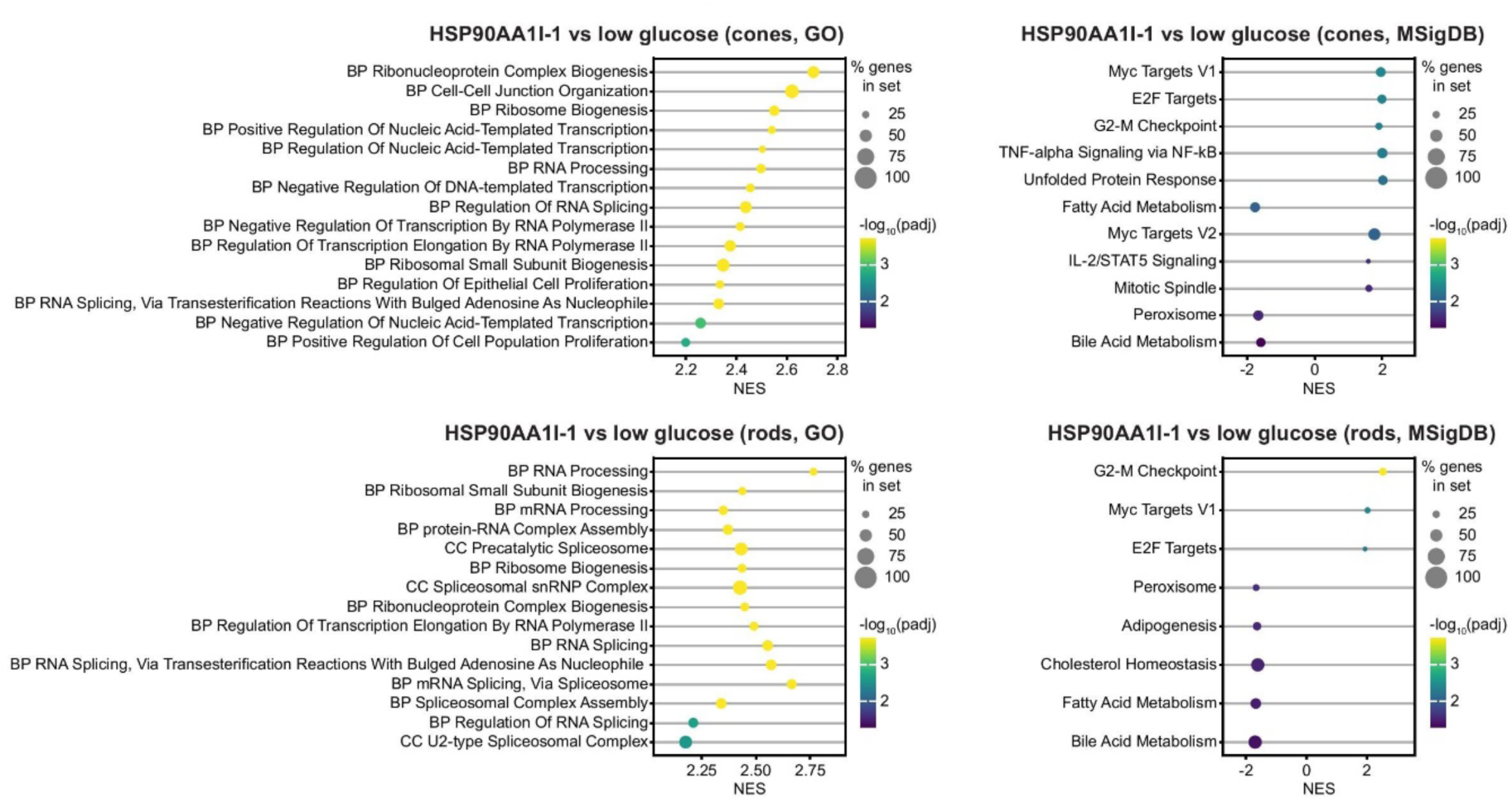
Geneset enrichment analysis for HSP90AA1I-1. Gene Ontology terms (GO, left) and Molecular Signatures Database Terms (MSigDB, right) enrichment analysis of HSP90AA1I-1-treated samples compared to low glucose in both cones (top) and rods (bottom). Dots represent normalized enrichment scores (NES) with their size indicating the percentage of differentially expressed genes within each gene set, and the color reflecting the adjusted p-value corrected for multiple testing (padj). All or the top 15 GO-terms are shown ordered by p-values, whereas all enriched MSigDB-terms are shown. BP: biological process, CC: cellular component.

**Figure S15:**
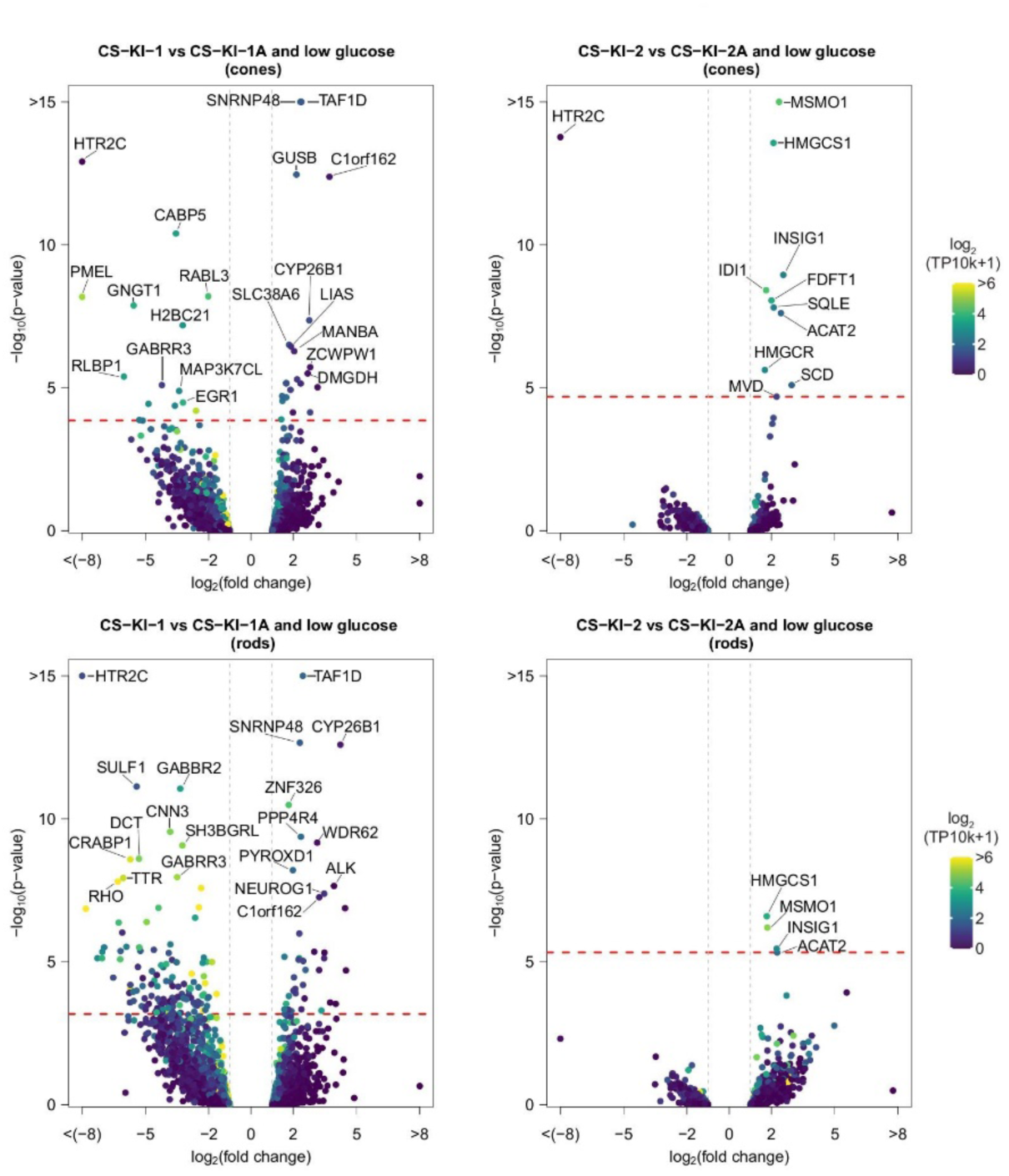
Differential gene expression analysis for cone-saving compounds. Differential gene expression comparing human retinal organoid cones (top) and rods (bottom) transcriptomes treated with CS-KI-1 compared to CS-KI-1A treatment and low glucose controls (left), and CS-KI-2 treatment compared to CS-KI-2A treatment and low glucose controls (right). Gray dotted lines indicate the threshold beyond which genes were included in analysis. Red dotted line indicates the p-value beyond which genes passed the significant threshold of p<0.05 after Benjamini-Hochberg correction for multiple testing. Colors indicate average gene expression in low glucose cones (top) or rods (bottom). Up to 10 most significantly up- or downregulated genes are labeled. TP10k, transcript counts per 10,000.

**Figure S16:**
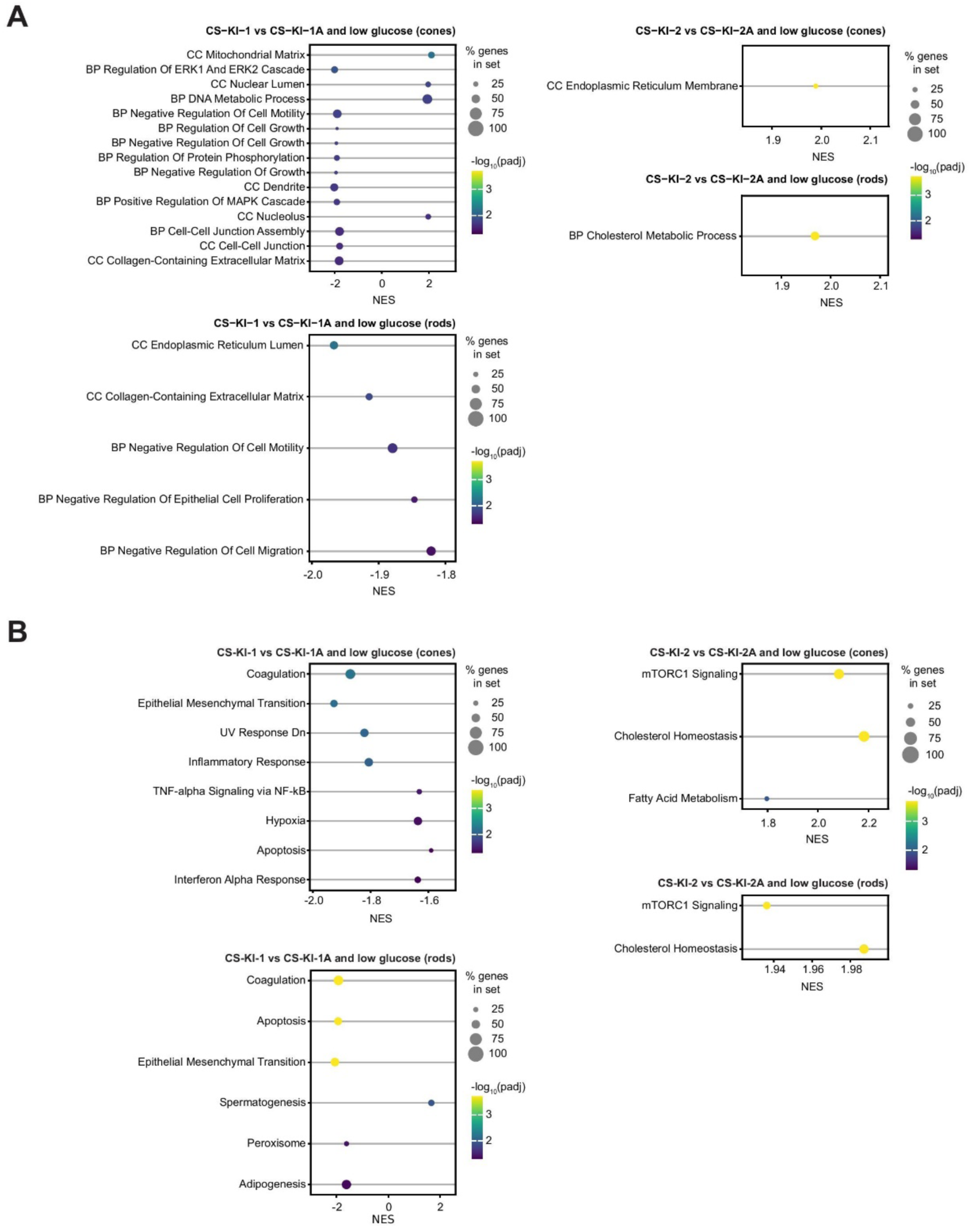
Geneset enrichment analysis for cone-saving compounds. **A-B:** Gene Ontology (GO) terms enrichment analysis (A) and Molecular Signatures Database (MSigDB) terms enrichment analysis (B) for specified comparisons in both cones (top) and rods (bottom). Dots represent normalized enrichment scores (NES) with their size indicating the percentage of differentially expressed genes within each gene set, and the color reflecting the adjusted p-value corrected for multiple testing (padj). All or the top 15 GO terms (A) or MSigDB terms (B) with the most significant p-values are displayed ordered by p-values.

**Figure S17:**
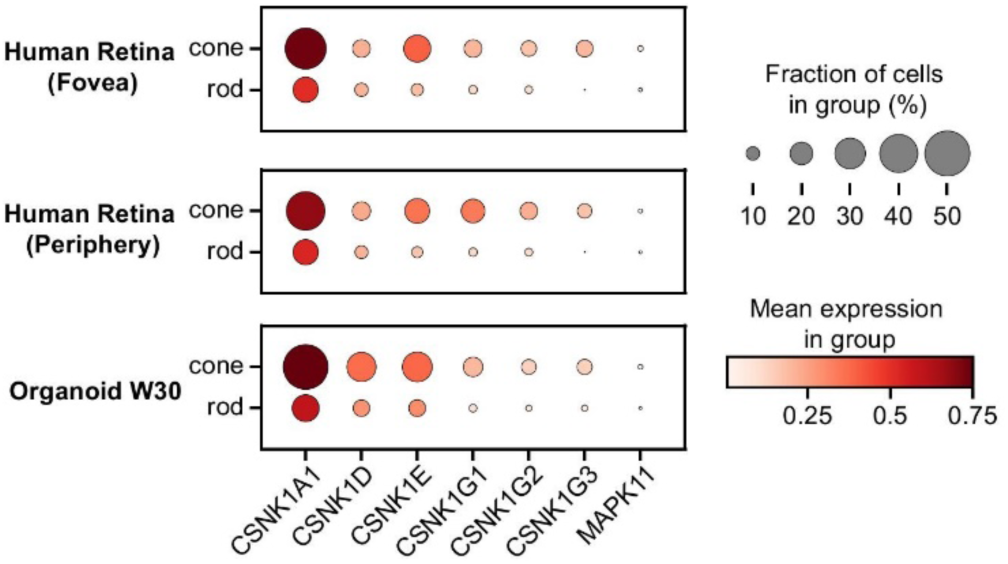
Expression of casein kinase 1 genes and MAPK11 in human retina and organoids. CK-1 and MAPK11 gene expression in human retina fovea and periphery and human retinal organoids at week 30 (W30) from published single cell atlases^22^. Color indicates mean expression across all cones or rods in UMI (unique molecular identifier) counts per 10000. Dot size indicates the fraction of cones or rods expressing the indicated gene.

