## Supplementary material for "Cell type-focused compound screen in human organoids reveals CK1 and MAPK11 inhibition protects cone photoreceptors from death": Supplemetary Tables

### Table S1

| Inchi-key | SMILES | MOA | GeneIDs | Median adj. cone survival (%) |
|---|---|---|---|---|
| BWDQBBCUWLSASG-MDZDMXLPSA-N | OCCN(CCC1=CNc2ccccc12)Cc3ccc(/C=C/C(=O)NO)cc3 | HDAC 1/2/3/6/8/10/11;Antimitotic Drugs;Apoptosis Inducers;Histone Deacetylase (HDAC) Inhibitors | HDAC9,HDAC6,HDAC5,HDAC7,HDAC8,HDAC11,HDAC10 | 0 |
| ATSUJKLLCSXMLV-BUHFOSPRSA-N | ONC(=O)/C=C/c4ccc(CNCCN3c1ccccc1c2ccccc23)cc4 | Histone Deacetylase (HDAC) Inhibitors | HDAC9,HDAC5,HDAC7,HDAC8,HDAC11,HDAC10 | 5 |
| NLKQTNGJKCBXMU-MDZDMXLPSA-N | ONC(=O)/C=C/c3ccc(CNCCc2cnc1ccccc1c2)cc3 | Histone Deacetylase (HDAC) Inhibitors | HDAC9,HDAC5,HDAC7,HDAC8,HDAC11,HDAC10 | 4 |
| HPTMTYKYKUKYCQ-UHFFFAOYSA-N | Cc1ccc(cc1NS(C)(=O)=O)Nc2cc(ncn2)c3ccccc3O | CDK1/2 inhibitor | CCNA1 | 7 |
| PIROHBFQHMKAPE-UHFFFAOYSA-N | CNc4nc1ccccc1c(NCc2ccc(cc2)NC(=O)c3ccc(Cl)nc3)n4 | beta-Catenin Inhibitors | CTNNB1,TCF4 | 9 |
| QPKKAIMODKGHRH-UHFFFAOYSA-N | CCN(Cc1cccc(c1)c3ccnc(Nc2cccc(c2)[N+]([O-])=O)n3)C(C)C | Antimitotic Drugs;Inhibitors of Signal Transduction Pathways;Aurora-A (ARK1) Kinase Inhibitors;Glycogen Synthase Kinase 3 beta (GSK-3beta\| tau Protein Kinase I) Inhibitors;Cyclin-Dependent Kinase Inhibitors | GSK3B,AURKA,CDKL5 | 10 |
| MUAICZWSFWUFNA-INIZCTEOSA-N | C[C@H](NCc1ccc4c(c1)N=C(NC(=O)C2=CC=C(S2)C3C=NNC=3)N4CC(C)(C)O)C(C)(C)C | ITK (EMT) Kinase Inhibitors;IL-4 Production Inhibitors;IL-2 Production Inhibitors | IL2,IL4,ITK | 4 |
| RLFKILXOLJVUNF-UHFFFAOYSA-N | COCc2c(ncc3Nc1ccc(cc1c23)OCc4ccccc4)C(=O)OC(C)C | GABA(A) BZ Site Receptor Partial Agonists | GABRA1,GABRA2,GABRA3,GABRA5,GABRB3,GABRG2 | 14 |
| PRDFBSVERLRRMY-UHFFFAOYSA-N | CCOc1ccc(cc1)C3Nc2ccc(cc2N=3)C5Nc4ccc(cc4N=5)N6CCN(C)CC6 | BCL2L1 gene inhibitor | BCL2L1 | 10 |
| JWOGUUIOCYMBPV-GMFLJSBRSA-N | CCC(=O)CCCCC[C@@H]4NC(=O)[C@H]1CCCCN1C(=O)[C@@H](NC(=O)[C@H](CC2=CN(OC)c3ccccc23)NC4=O)[C@@H](C)CC | HDAC inhibitor | HDAC3,HDAC4 | 13 |
| SMYUEWYXIKCMEA-UHFFFAOYSA-N | Cc1ccc(cn1)OC2CCN(CC2)c4ncnc3ccc(F)cc34 | Positive allosteric modulators (PAMs) of the muscarinic acetylcholine | CHRM4 | 12 |
| VTGBZWHPJFMTKS-UHFFFAOYSA-N | CCc1ccc(cc1)OCc2ccccc2C(=O)Nc4ccc3nc(C)cc(N)c3c4 | OPRL1 gene inhibitor;OPRM1 gene inhibitor | OPRL1,OPRM1 | 11 |
| SETVRSKZJJWOPA-FLDGXQSCSA-N | C/C(/C=C/[C@@]1(C)[C@H](C)CCC(=O)[C@@H]1C)=C\Cc2c(O)c(Cl)c(C)c(C=O)c2O | "AP-1 Inhibitors;Electron Transport Chain Inhibitors";AP-1 Inhibitors;Electron Transport Chain Inhibitors | JUN,UQCRC1 | 11 |
| RBTBFTRPCNLSDE-UHFFFAOYSA-N | CN(C)c3ccc2N=C1C=CC(C=C1Sc2c3)=[N+](C)C | Nitric Oxide Production Inhibitors;Tau Aggregation Inhibitors | MAPT | 13 |
| JUVIOZPCNVVQFO-HBGVWJBISA-N | COc5cc4OC[C@H]3Oc1c(ccc2O[C@H](Cc12)C(C)=C)C(=O)[C@H]3c4cc5OC | Electron transport chain inhibitor;Apoptosis Inducers;NADH-Ubiquinone Oxidoreductase (Complex I) Inhibitors;Non-Steroidal Antiinflammatory Drugs;Electron Transport Chain Inhibitors;"Electron transport chain inhibitor;Apoptosis Inducers;NADH-Ubiquinone Oxidoreductase (Complex I) Inhibitors;Non-Steroidal Antiinflammatory Drugs;Electron Transport Chain Inhibitors" | NDUFS1,NOX4,NOX5 | 19 |
| ZLUZDKXBTNQWOL-MDZDMXLPSA-N | ONC(=O)/C=C/c3ccc(CNCCC1=CNc2ccccc12)cc3 | HDAC inhibitor | HDAC9,HDAC6,HDAC5,HDAC7,HDAC8,HDAC11,HDAC10 | 13 |
| ILAZEUQGANOTAY-VAWYXSNFSA-N | CC(C)N(CCC1=CNc2ccccc12)Cc3ccc(/C=C/C(=O)NO)cc3 | Histone Deacetylase (HDAC) Inhibitors | HDAC9,HDAC5,HDAC7,HDAC8,HDAC11,HDAC10 | 15 |
| OTSOOHRUMBRSHZ-UHFFFAOYSA-N | COCC[N+]4=C(C)N(Cc1cnccn1)C3C(=O)c2ccccc2C(=O)C=34 | BIRC5 gene inhibitor | BIRC5 | 14 |

| Inchi-key | SMILES | MOA | GeneIDs | Median adj. cone survival (%) |
|---|---|---|---|---|
| XQYZDYMELSJDRZ-UHFFFAOYSA-N | COc3ccc(Cc1nccc2cc(OC)c(cc12)OC)cc3OC | Phosphodiesterase PDE10A Inhibitors | PDE4C,PDE10A,ENPP6 | 11 |
| ALBKMJDFBZVHAK-UHFFFAOYSA-N | CCOC(=O)c3ncc2Nc1ccc(cc1c2c3COC)OCc4ccccc4 | Anxiolytics | GABRA1,GABRA2,GABRA3,GABRB3,GABRG2 | 10 |
| NUUSUAWULNXMGF-UHFFFAOYSA-N | O=C(Nc1ccc(Cl)c(Cl)c1)Nc3ccc2N=NSc2c3 | Eukaryotic Translation Initiation Factor 2-alpha Kinase 1 (HRI) Activators | EIF2AK1 | 12 |
| FUGQNAUKABUDQI-UHFFFAOYSA-N | CC(C)C2=Cc1c(ncnc1S2)N3CCN(CC3)C4=NCC(C)(C)S4 | Inhibitor of MEN1 (Menin);MEN1 inhibitor | MEN1 | 18 |
| SCJXQZZYGYLKJG-CQSZACIVSA-N | CC[C@@H](N)CNc1ccnc(n1)c2cc(ccc2O)C3C=NN(C)C=3 | protein kinase d inhibitor | PKD1 | 18 |
| RDALZZCKQFLGJP-UHFFFAOYSA-N | CC2=Cc1c(cccc1N2c5nc3CCOCc3c(NCc4ccccc4)n5)C(N)=O | p97 AAA ATPase inhibitor | VCP | 16 |
| IYVIAGDMQOYYQC-OUKQBFOZSA-N | ONC(=O)/C=C/c2ccc(CNCCCc1ccccc1)cc2 | Histone Deacetylase (HDAC) Inhibitors | HDAC9,HDAC5,HDAC7,HDAC8,HDAC11,HDAC10 | 19 |
| VQQRBBFRJRBWPF-UHFFFAOYSA-N | CCN4CCN(Cc1ccc(cc1C(F)(F)F)NC(=O)Nc2ccc(cc2)Oc3ccnc(N)n3)CC4 | Flt3 (FLK2/STK1) Inhibitors | FLT3 | 13 |
| SKBXAZQJXDJNFI-RMKNXTFCSA-N | COc3ccc2NC=C(CCNCc1ccc(/C=C/C(=O)NO)cc1)c2c3 | Histone Deacetylase (HDAC) Inhibitors | HDAC9,HDAC5,HDAC7,HDAC8,HDAC11,HDAC10 | 12 |
| XQYASZNUFDVMFH-CQSZACIVSA-N | C[C@@H]2CN(Cc1ccc(F)cc1)CCN2C(=O)COc3ccc(Cl)cc3NC(N)=O | Chemokine CCR1 Antagonists | CCR1 | 15 |
| YJGVMLPVUAXIQN-XVVDYKMHSA-N | COc1cc(cc(OC)c1OC)[C@@H]4c3cc2OCOc2cc3[C@H](O)[C@H]5COC(=O)[C@H]45 |  | TUBA4A,TUBB2A,TUBA1A,TUBA1B,TUBB3,TUBB4A,TUBB4B,TUBB1,TUBB6,TUBA1C,TUBA3E,TUBA3D,TUBB,TUBB8,TUBB2B,TUBA3C,TUBG1,TUBG2,TUBD1,TUBA8 | 17 |
| OKJDLRMQHZRYOZ-UHFFFAOYSA-N | NCCCN4C(=O)c1cc(ccc1C3C(=O)c2ccc(O)cc2C=34)[N+]([O-])=O | Dual Top1, TDP1 inhibitors | TOP1,TDP1 | 16 |
| BPRNMVDTWIHULJ-AWEZNQCLSA-N | C[C@@H](CN1CCN(CC1)S(=O)(=O)c2ccc(Cl)c(Cl)c2)Nc3ncnc4C(C)=CSc34 | Lysophosphatidate-2 receptor antagonist | LPAR2 | 22 |
| BPNUQXPIQBZCMR-IBGZPJMESA-N | CC1=NNc2ccc(cc12)c3cncc(c3)OC[C@@H](N)Cc4ccccc4 | cAMP-Dependent Protein Kinase (PKA) Inhibitors;Inhibitors of Signal Transduction Pathways;PKB alpha/Akt1 Inhibitors | PRKACA,PRKACB,PRKACG | 22 |
| XVOOCQSWCCRVDY-UHFFFAOYSA-N | COc4cc(Br)c(NC1=NC(=CS1)C2=C(C)N=C3N=CC=CN23)c(Br)c4 | Polycomb Complex Protein BMI-1 Inhibitors | BMI1 | 16 |
| AWIVHRPYFSSVOG-UHFFFAOYSA-N | Fc3ccc(CNc1ncnc2ccc(F)cc12)cc3 | Autophagy Agonist;Phosphodiesterase V (PDE5A) Inhibitors | PDE5A | 15 |
| WOFGKOQKJKFJLB-UHFFFAOYSA-N | COCCCc1cc(ccn1)C2=CNc3cc(ccc23)C(=O)c4ccc(Cl)c(c4)S(N)(=O)=O | MMP-13 (Collagenase 3) Inhibitors;MMP-2 (Gelatinase A) Inhibitors | MMP2,MMP13 | 22 |
| SXHHURZUIGDYAR-QGZVFWFLSA-N | C[C@@H]1CCCN1c3cc(cc(Nc2cc(C#N)ccn2)n3)C4(C#N)CCN(C)CC4 | MAP3K12 (DLK) Inhibitors | MAP3K12 | 12 |
| SGYJGGKDGBXCNY-ZDIDWYTNSA-N | CC4(C)NC(=O)[C@H](CCCCCC(=O)C1CO1)NC(=O)[C@H]2CCCN2C(=O)[C@H](Cc3ccccc3)NC4=O | Reported to be a general HDAC inhibitor | HDAC1,HDAC2,HDAC3,HDAC4,HDAC6,HDAC5 | 19 |
| GAKKPKBEHVIOLH-UHFFFAOYSA-N | CNC1=NC(C)=C(S1)c4ccnc(Nc2ccc(cc2)N3CCNCC3)n4 | CDK Inhibitor;Pan Kinase Inhibitor | CDK2,CCNE2 | 20 |
| CHILCFMQWMQVAL-UHFFFAOYSA-N | Oc1ccc(Cl)cc1C(=O)Nc2cc(cc(c2)C(F)(F)F)C(F)(F)F | IKK-2 (IKK-beta) Inhibitors;NF-kappaB (NFKB) Activation Inhibitors | IKBKB | 14 |
| KPBNHDGDUADAGP-VAWYXSNFSA-N | O=C(/C=C/c1cccnc1)NCCCCC2CCN(CC2)C(=O)c3ccccc3 | Angiogenesis Inhibitors;Apoptosis Inducers;Nicotinamide Phosphoribosyltransferase (NMPRTase) Inhibitors | NAMPT | 23 |

| Inchi-key | SMILES | MOA | GeneIDs | Median adj. cone survival (%) |
|---|---|---|---|---|
| VJXBYUITQBTTQM-DSRNDQRRSA-N | CC[C@H](C)[C@H]2NC(=O)[C@H]1CSSCCC=C[C@H](CC(=O)N[C@H](CCSC)C(=O)N1)OC(=O)C[C@@H]2O | HDAC inhibitor (no hydroxamic acid) | HDAC1,HDAC2,HDAC3,HDAC6,HDAC8 | 15 |
| IYNDTACKOAXKBJ-UHFFFAOYSA-N | OCCCNc1cc(ccn1)c3ccnc(Nc2cccc(Cl)c2)n3 | Protein Kinase C (PKC) Inhibitors;CDK1 Inhibitors;CDK2 Inhibitors | CDK1,CDK2 | 15 |
| IEENMDADGAZPAM-UHFFFAOYSA-N | CCC(C)C6C(=O)N1NCCCC1C(=O)N2NCCCC2C(=O)N(C)C(Cc3ccccc3)C(=O)N4CCCC4C(=O)NC(Cc5ccccc5)C(=O)N6O | Vasopressin (AVP) V1a Antagonists;Vasopressin (AVP) V2 Antagonists | AVPR1A,AVPR2 | 13 |
| VFBGXTUGODTSPK-BAQGIRSFSA-N | O=C3Nc2ccc1N=CSc1c2/C/3=C/C4=CNC=N4 | PKR Inhibitor | EIF2AK2 | 18 |
| HCAQGQIHBFVVIX-LYXAAFRTSA-N | C/C(=N\NC(N)=N)/c1ccc(cc1)NC(=O)Nc2ccc(cc2)/C(/C)=N/NC(N)=N | Ribonuclease P Inhibitors;Checkpoint Kinase 2 (Chk2) Inhibitors | CHEK2 | 20 |
| BFXLAXBXCXOWNH-UHFFFAOYSA-N | [O-][N+](=O)c1cc(ccc1Cl)S(=O)(=O)Nc2ccc(Cl)cc2C(=O)Nc3ccc(Cl)cc3 | Phosphopantetheine Adenylyltransferase (PPAT) Inhibitors | COASY | 24 |
| BBLGCDSLCDDALX-LKGBESRRSA-N | C/C=C(\C)/[C@H](O)[C@H](C)/C=C(\C)/C=C/C/C(/C)=C/Cc1nc(OC)c(OC)c(O)c1C | NADH oxidase inhibitors;Electron transport chain inhibitor | NDUFAB1,NDUFS1,NDUFV1 | 19 |
| FUSNMLFNXJSCDI-UHFFFAOYSA-N | Cc1cccc(c1)N(C)C(=S)Oc3ccc2ccccc2c3 | Fungal Squalene Monooxygenase Inhibitors | SQLE | 24 |
| OWBFCJROIKNMGD-BQYQJAHWSA-N | COc2cc(OC)c(/C=C/S(=O)(=O)Cc1ccc(OC)c(c1)NCC(O)=O)c(c2)OC | Antagonist of Raf-Ras interaction | BRAF,RAF1,ZHX2 | 24 |
| HRNLUBSXIHFDHP-UHFFFAOYSA-N | Nc1ccccc1NC(=O)c4ccc(CNc2nccc(n2)c3cccnc3)cc4 | HDAC1/2;Apoptosis Inducers;Histone Deacetylase 1 (HDAC1) Inhibitors | HDAC1,HDAC2,HDAC3 | 21 |
| WQAVPPWWLLVGFK-VTNASVEKSA-N | COc4ccc(C[C@H](NC(=O)[C@H](C)NC(=O)CN1CCOCC1)C(=O)N[C@@H](Cc2ccccc2)C(=O)[C@@]3(C)CO3)cc4 | Immunoproteasome Inhibitors | PSMB8 | 18 |
| DNODJHQYSZVNMH-UHFFFAOYSA-N | Nc2ccc(O)c3C(=O)N(C1CCC(=O)NC1=O)C(=O)c23 | CRBN neomorph;E3 ligase inhibitor | CRBN | 22 |
| IAKHMKGGTNLKSZ-INIZCTEOSA-N | COc3cc2CC[C@H](NC(C)=O)C1=CC(=O)C(=CC=C1c2c(OC)c3OC)OC | Tubulin Polymerase Inhibitors | TUBA4A,TUBB2A,TUBA1A,TUBA1B,TUBB3,TUBB4A,TUBB4B,TUBB1,TUBB6,TUBA1C,TUBA3E,TUBA3D,TUBB,TUBB8,TUBB2B,TUBA3C,TUBG1,TUBG2,TUBD1,TUBA8 | 13 |
| WQBLEMAGSGUUGW-UHFFFAOYSA-N | Nc1cc(ccn1)c3ccnc(Nc2cccc(Cl)c2)n3 | CDK1 Inhibitors | CDK1 | 24 |
| YJGVMLPVUAXIQN-HAEOHBJNSA-N | COc1cc(cc(OC)c1OC)[C@@H]4c3cc2OCOc2cc3[C@H](O)[C@H]5COC(=O)[C@@H]45 | CASP3 activator;apoptosis inhibitor;IGF1R inhibitor;IGF-1R Inhibitors;Caspase 3 Activators;Apoptosis Inducers | CASP3,IGF1R | 17 |
| ZCURBFFRNVPIAS-CYBMUJFWSA-N | CCCCCC(=O)N3C[C@@H](CCl)c2c1ccccc1c(O)cc23 | covalent inhibitor of ALDH1A1 | ALDH1A1 | 22 |
| GVIFMXAPEOVXKO-UHFFFAOYSA-N | COc5cc(CN4N=C(CCN1CCN(CC1)c2cccc(Cl)c2C)c3cc(OC)c(cc34)OC)ccc5OC | Calmodulin Antagonists | CALM1 | 21 |
| LUTPUCJIKBULCQ-UHFFFAOYSA-N | CC2CCc1cc(F)ccc1N2C(=O)COc4cccc3cnccc34 | Bile Acid Responsive TGR5 Receptors (GPBAR1, AXOR 109, GPCR19) Agonists | GPBAR1 | 17 |
| QLJDJSOFNVSHNG-VMPITWQZSA-N | ONC(=O)/C=C/c3ccc(CNCCC1=CNc2ccc(F)cc12)cc3 | Histone Deacetylase (HDAC) Inhibitors | HDAC9,HDAC5,HDAC7,HDAC8,HDAC11,HDAC10 | 19 |

| Inchi-key | SMILES | MOA | GeneIDs | Median adj. cone survival (%) |
|---|---|---|---|---|
| UIFFUZWRFRDZJC-SBOOETFBSA-N | CCCCCC[C@H]2C(=O)O[C@H](C)[C@H](NC(=O)c1cccc(NC=O)c1O)C(=O)O[C@@H](C)[C@@H]2OC(=O)CC(C)C | Electron transport chain inhibitor;Cytochrome c reductase;Electron Transport Chain Inhibitors | UQCRC1 | 30 |
| SDABUVHLPSIZEG-NBMRYCAZSA-N | CN4C(=O)[C@@]35C[C@](C)(C#N)[C@H](c2ccc1OCOc1c2)N3C(=O)[C@]4(C)SS5 | induces concomitant H3K9me3 downregulation | SUV39H1 | 31 |
| XWQVQSXLXAXOPJ-NJDAHSKKSA-N | COC[C@@H](C)N[C@@H]1CC[C@H](CC1)Nc2cc(c(Cl)cn2)c4cccc(NCC3(C#N)CCOCC3)n4 | CDK9/Cyclin T1 Inhibitors | CCNT1,CDK9 | 28 |
| JGDCRWYOMWSTFC-AZGSIFHYSA-N | C[C@]12CC[C@H](O)C[C@H]1CC[C@@H]3[C@@H]2[C@H](O)C(=O)[C@]4(C)[C@H](CC[C@]34O)C5C=CC(=O)OC=5 | Arenobufagin, SMUT | PSMB1,PSMB2,PSMB5,ATP1B4 | 23 |
| QSYLKMKIVWJAAK-UHFFFAOYSA-N | Cc5cc(Nc1ccc(cc1)NC(=O)c2ccc(cc2)Nc4ccnc3ccccc34)nc(N)n5 | DNA Methyltransferase I Inhibitors | DNMT1 | 23 |
| OHAXNCGNVGGWSO-UHFFFAOYSA-N | Oc2cc1ccccc1cc2C(=O)Nc3ccc(Cl)cc3 | Cyclic AMP Response Element-Binding Protein (CREB) Inhibitors | CREB1 | 28 |
| DYLJVOXRWLXDIG-UHFFFAOYSA-N | COc1cccc(OC)c1C2=CC(=NN2c3ccnc4cc(Cl)ccc34)C(=O)NC6(C(O)=O)C5CC7CC(C5)CC6C7 | Carboxypeptidase A Inhibitors;Neurotensin NTS1 (NT1) Receptor Antagonists | NTSR1 | 29 |
| XUSKJHCMMWAAHV-SANMLTNESA-N | CC[C@@]1(O)C(=O)OCc2c1cc3n(Cc4c(c5cc(O)ccc5nc43)[Si](C)(C)C(C)(C)C)c2=O | DNA-Intercalating Drugs;DNA Topoisomerase I Inhibitors | TOP1 | 25 |
| ZGLXUQQMLLIKAN-SVIJTADQSA-N | COc1cc(cc(OC)c1OC)[C@@H]4c3cc2OCOc2cc3C[C@H]5COC(=O)[C@H]45 | Cyclooxygenase-2 Inhibitors;Non-Steroidal Antiinflammatory Drugs;Angiogenesis Inhibitors | CASP3,PTGS2 | 34 |
| ULXXDDBFHOBEHA-CWDCEQMOSA-N | CN(C)C/C=C/C(=O)Nc3cc2c(Nc1ccc(F)c(Cl)c1)ncnc2cc3O[C@H]4CCOC4 | Irreversible EGFR (HER1;erbB1) Inhibitors;HER2 (erbB2) Inhibitors;Inhibitors of Signal Transduction Pathways | EGFR,ERBB2,ERBB4 | 23 |
| KQNZDYYTLMIZCT-KQPMLPITSA-N | C[C@H]2CCC/C=C/[C@@H]1C[C@H](O)C[C@H]1[C@H](O)/C=C/C(=O)O2 | Apoptosis Inducers;Caspase 3 Activators;Autophagy inducer;"Apoptosis Inducers;Caspase 3 Activators;Autophagy inducer" | ARF1,GBF1,CYTH2,ARFGEF2,ARFGEF1,CYTH1 | 28 |
| CDOVNWNANFFLFJ-UHFFFAOYSA-N | C1CN(CCN1)c2ccc(cc2)C4C=NC3=C(C=NN3C=4)c6ccnc5ccccc56 | Activin Receptor Like Kinase 3 (ALK3 BMPR-IA) Inhibitors;Activin Receptor Like Kinase 2 (ALK2 ActR-IA) Inhibitors | ACVR1,BMPR1A,BMPR1B | 49 |
| KAKPGJJRYRYSTP-UHFFFAOYSA-N | COc1cc(cc(c1)N2CCN(CC2)C(=O)Nc4nc3cc(F)ccc3nc4OC)OC | phosphorylated-p68 RNA helicase inhibitor | DDX5 | 24 |
| GUGBOAVFOFNJFG-VAWYXSNFSA-N | ONC(=O)/C=C/c3ccc(CNCCCC1=CNc2ccccc12)cc3 | Histone Deacetylase (HDAC) Inhibitors | HDAC9,HDAC5,HDAC7,HDAC8,HDAC11,HDAC10 | 30 |
| UTOXGQNLFXWCMS-QGZVFWFLSA-N | CS(=O)(=O)N1CCC(CC1)C2=NOC(=N2)[C@H](CCCC3CCCCC3)CC(=O)NO | Procollagen C-Proteinase Inhibitors | BMP1 | 28 |
| INVTYAOGFAGBOE-UHFFFAOYSA-N | Nc1ccccc1NC(=O)c3ccc(CNC(=O)OCc2cccnc2)cc3 | HDAC1/3;Wnt pathway agonist;Histone deacetylase-1 inhibitor;Histone deacetylase-2 inhibitor | HDAC1,HDAC2,HDAC3 | 35 |
| QGQFNQWZHHSZIC-UHFFFAOYSA-N | CC1NC(CO)=C(C(=O)C=1Cl)c2ccc(cc2)Oc3ccc(cc3)OC(F)(F)F | "P. falciparum Cytochrome b-c1 Complex (Complex III subunit 3) Inhibitors;Electron Transport Chain Inhibitors";P. falciparum Cytochrome b-c1 Complex (Complex III subunit 3) Inhibitors;Electron Transport Chain Inhibitors | UQCRC1 | 35 |
| ODPGGGTTYSGTGO-UHFFFAOYSA-N | CCN4CCN(Cc1ccc(cc1C(F)(F)F)NC(=O)Nc2ccc(cc2)Oc3cc(NC)ncn3)CC4 | Flt3 (FLK2/STK1) Inhibitors;Inhibitors of Signal Transduction Pathways | MAPK14,FLT3,KIT,PDGFRB,MAPK8,MAPK10 | 22 |
| QJZRFPJCWMNVAV-HHHXNRCGSA-N | CC(C)[C@H](C2=Nc1cc(Cl)ccc1C(=O)N2Cc3ccccc3)N(CCCN)C(=O)c4ccc(C)cc4 | Antimitotic Drugs;Kinesin-Like Spindle Protein KIF11 (KSP, Eg5) Inhibitors | KIF11 | 25 |
| ITFBYYCNYVFPKD-FMIDTUQUSA-N | CC5(C)CC[C@@]4(CC[C@]3(C)[C@H](C(=O)C=C2[C@@]1(C)C=C(C#N) | Apoptosis Inducers;Nitric Oxide Production Inhibitors;Nuclear Factor, Erythroid Derived 2, Like 2 (Nrf2) Activators;Antiinflammatory Drugs;PPARgamma | NFE2L2,PPARG,KEAP1 | 37 |

| Inchi-key | SMILES | MOA | GeneIDs | Median adj. cone survival (%) |
|---|---|---|---|---|
| | C(=O)C(C)(C)[C@@H]1CC[C@]23C)[C@@H]4C5)C(=O)N6C=CN=C6 | Agonists;"Apoptosis Inducers;Nitric Oxide Production Inhibitors;Nuclear Factor, Erythroid Derived 2, Like 2 (Nrf2) Activators;Antiinflammatory Drugs;PPARgamma Agonists" | | |
| XFMZYCHKCPPZQW-HXUWFJFHSA-N | CN(C)C[C@@H](OC(=O)N3CC2=C(NC(=O)c1ccc(F)cc1)NN=C2C3(C)C)c4ccccc4 | PAK4 gene inhibitor | PAK4 | 27 |
| MRAMUVZVDBOPEH-JOCHJYFZSA-N | O=C(CCl)N([C@@H]1CCCN(C1)C(=O)c2ccc(cc2)N3CCOCC3)c4ccccc4 | covalent modifier of catalytic cysteine of pro-CASP8 | CASP8 | 16 |
| WPTTVJLTNAWYAO-KPOXMGGZSA-N | COC(=O)[C@]15CCC(C)(C)C[C@H]1[C@H]4C(=O)C=C3[C@@]2(C)C=C(C#N)C(=O)C(C)(C)[C@@H]2CC[C@@]3(C)[C@]4(C)CC5 | Angiogenesis Inhibitors;Antiinflammatory Drugs;Apoptosis Inducers;Bcl-2 Inhibitors;Glutathione Reductase (NADPH) Activators;Heme Oxygenase Activators;IKK-1 (IKK-alpha) Inhibitors;NF-kappaB (NFKB) Activation Inhibitors;Nitric Oxide Production Inhibitors;Nuclear Factor, Erythroid Derived 2, Like 2 (Nrf2) Activators;PPARgamma Agonists | KEAP1,IKBKB | 41 |
| QAIPRVGONGVQAS-DUXPYHPUSA-N | OC(=O)/C=C/c1ccc(O)c(O)c1 | 5-Lipoxygenase Inhibitors;HIV Integrase Inhibitors;Antioxidants | ALOX5 | 30 |
| JDJGAAQTPZJIDZ-UHFFFAOYSA-P | C[n+]2cccc1cc(ccc12)NC(=O)c3cccc(c3)C(=O)Nc5ccc4c(ccc[n+]4C)c5 | Telomerase Inhibitors;DNA G-quadruplex (G4) Ligands | TERT | 28 |
| DFBIRQPKNDILPW-KTGKZQHOSA-N | CC(C)[C@]17O[C@H]1[C@@H]2O[C@]26[C@]3(O[C@H]3CC5C4COC(=O)C=4CC[C@@]56C)[C@@H]7O | ERCC3 (TFIIH subunit) | ERCC3 | 35 |
| FJHBVJOVLFPMQE-QFIPXVFZSA-N | CCc4c1cc(O)ccc1nc5C3=CC2=C(COC(=O)[C@]2(O)CC)C(=O)N3Cc45 | Apoptosis Inducers;DNA Topoisomerase I Inhibitors | TOP1 | 27 |
| NSMRMZWAHUBRPG-UHFFFAOYSA-N | NC(=O)CCNc1cc(ccn1)c3ccnc(Nc2cccc(Cl)c2)n3 | CDK1 Inhibitors | CDK1 | 30 |
| MYKJVLTXPNIGOV-KTKRTIGZSA-N | CCN(C/C=C\c2ccc(C1CCCCC1)c(Cl)c2)C3CCCCC3 | sigma1 Receptor Ligands;sigma2 Receptor Ligands | SIGMAR1 | 30 |
| NDDAHWYSQHTHNT-UHFFFAOYSA-N | CC2Cc1ccccc1N2NC(=O)c3ccc(Cl)c(c3)S(N)(=O)=O | Carbonic Anhydrase Type VII Inhibitors | CA4,CA5A,CA7,CA12,CA5B | 38 |
| KXMZDGSRSGHMMK-VWLOTQADSA-N | NC4=NC(Nc2ccc1CC[C@@H](CCc1c2)N3CCCC3)=NN4c7cc6CCCc5ccccc5c6nn7 | Axl tyrosine kinase receptor inhibitor | AXL | 22 |
| JADDQZYHOWSFJD-FLNNQWSLSA-N | CCNC(=O)[C@H]1O[C@H]([C@H](O)[C@@H]1O)N3C=Nc2c(N)ncnc23 | Adenosine Receptor Agonists | ADORA2A | 31 |
| BCWCEHMHCDCJAD-UHFFFAOYSA-N | Cc1ccc(cc1)C(=O)C(=O)c2ccc(C)cc2 | Carboxylesterase Inhibitors | CES1,CES2,CES3,CES1P1,CES5A | 35 |
| PCFKMZRPZATASD-UHFFFAOYSA-N | CS(=O)(=O)N5CC1(CCN(CC1)C(=O)Nc2cnc(cn2)c3ccccc3)c4ccccc45 | Neuropeptide Y5 (NPY Y5) Antagonists | NPY5R | 43 |
| AUVVAXYIELKVAI-CKBKHPSWSA-N | CC[C@H]3CN2CCc1cc(OC)c(cc1[C@@H]2C[C@@H]3C[C@H]4NCCc5cc(OC)c(cc45)OC)OC | Translation inhibitor | RPS14,RPS20 | 34 |
| TXJZRSRTYPUYRW-NQIIRXRSSA-N | COC(=O)c1ccc(cc1)[C@H]4C3Nc2ccccc2C=3C[C@H](C(=O)OC)N4C(=O)CCl | GPX4 covalent inhibitor | GPX4 | 20 |
| DZIUPOCVDSYPSY-UHFFFAOYSA-N | Cc1c(ccc2NC=Cc12)Nc3c(C#N)cncc3C6=Cc5cc(CN4CCN(C)CC4)ccc5O6 | Protein Kinase PKC theta Inhibitors | PRKCQ | 39 |
| RDONXGFGWSSFMY-UHFFFAOYSA-N | CCS(=O)(=O)Nc2ccc(Oc1ccc(F)cc1F)c(c2)C3=CN(C)C(=O)C4NC=CC3=4 | Bromodomain-Containing Protein 4 (Brd4, HUNK1) Inhibitors | BRD4 | 33 |
| CEGSUKYESLWKJP-UHFFFAOYSA-N | C(CC1=CNc2ccccc12)Nc3ccc(cc3)Nc4ccncc4 | MDM2 (hdm2) Inhibitors | MDM2 | 34 |
| FJDDSMSDZHURBJ-UHFFFAOYSA-N | COc2ccc1NC(I)=C(CCNC(C)=O)c1c2 | MTNR1A agonist;MTNR1B agonist | MTNR1A,MTNR1B | 38 |
| YKJYKKNCCRKFSL-BFHYXJOUSA-N | COc2ccc(C[C@@H]1NC[C@@H](O)[C@@H]1OC(C)=O)cc2 | translation inhibitor | RPL3,RPL8,RPL11,RPL15,RPL19,RPL23A,RPL | 45 |

| Inchi-key | SMILES | MOA | GeneIDs | Median adj. cone survival (%) |
|---|---|---|---|---|
| | | | 37,RPL23,RPL13A,RPL 26L1,RSL24D1,RPL10L | |
| JRWROCIMSDXGOZ-UHFFFAOYSA-N | CC(C)(C)c1ccc(cc1)S(=O)(=O)Nc2ccc(Cl)cc2C(=O)c3cc[n+]([O-])cc3 | CCR9 chemokine antagonist | CCR9 | 39 |
| RVKFQAJIXCZXQY-CBZIJGRNSA-N | C[C@]34CC[C@@H]2c1ccc(cc1CC[C@H]2[C@@H]3CCC4=O)OS(N)(=O)=O | Estrogen Receptor (ER) Agonists;Steryl Sulfatase Inhibitors | STS | 41 |
| RPDFDSQFBCJTDY-GAQXSTBRSA-N | CC(C)Oc6cc(CC(=O)N4CCC[C@@](CC[N+]12CCC(CC1)(CC2)c3ccccc3)(C4)c5cc(Cl)c(Cl)cc5)ccc6 | Tachykinin NK1 Antagonists | TACR1 | 38 |
| FDWQSLRDIBRKEI-UHFFFAOYSA-N | CC(C)(C)C3=CN=C(CSC2=CN=C(NC(=O)Cc1ccc(CNC(CO)CO)cc1)S2)O3 | CDK2/Cyclin E Inhibitors;CDK1 Inhibitors;CDK4 Inhibitors | CCNE1,CDK1,CDK2,CD K4,CCNE2 | 33 |
| ULKIYKLTKXRBOD-UHFFFAOYSA-N | OC(=O)C(CP(O)(O)=O)c1ccccc1 | Glutamate Carboxypeptidase II (NAALADase;NAAG Peptidase, FOLH1, PSMA) Inhibitors | FOLH1 | 34 |
| NICHJJOSEXYBED-AMGIVPHBSA-N | CC(C)C[C@H](NC(=O)CCN(C)C)c1cc(Cl)ccc1N2CCN(CC2)C(=O)[C@H](C)Cc3ccc(Cl)cc3 | Melanocortin MC4 Receptor Antagonists | MC4R | 37 |
| HUNGUWOZPQBXGX-UHFFFAOYSA-N | O=C(Cc1ccc(cn1)c2ccc(cc2)OCCN3CCOCC3)NCc4ccccc4 | c-src allosteric inihitor;allosteric Src inhibitor;Src Kinase Inhibitors;Antimitotic Drugs;Inhibitors of Signal Transduction Pathways;Tubulin polymerization inhibitors | SRC,TUBG2 | 38 |
| IRGAIDAWHGYOKD-UHFFFAOYSA-N | COc1cc(F)c(cc1OC)C4Nc3ncc(Cl)c(NCC2CCNCC2)c3N=4 | JAK1 selective inhibitors | CDK2,JAK1,AURKB | 36 |
| NCNRHFGMJRPRSK-MDZDMXLPSA-N | ONC(=O)/C=C/c1cccc(c1)S(=O)(=O)Nc2ccccc2 | HDACs;Apoptosis Inducers;Histone Deacetylase 1 (HDAC1) Inhibitors;Histone Deacetylase 2 (HDAC2) Inhibitors;Angiogenesis Inhibitors | HDAC1,HDAC2,HDAC9, HDAC5,HDAC7,HDAC8, HDAC11,HDAC10 | 30 |
| RGXYYAZGELLKDA-UHFFFAOYSA-N | [O-][N+](=O)C1=CC=C(SCCCCCCO)C2=NON=C12 | Glutathione-S-Transferase P1 (GSTP1) Inhibitors | GSTM2,GSTP1 | 36 |
| RZKDEGZIFSJVNA-IBGZPJMESA-N | C[C@H]4CN(Cc3ccc(CC(=O)N1CCC(CC1)Nc2cccc(F)c2)cc3)CCN4 | Motilin Receptor Agonists | MLNR | 34 |
| YUJFUSDUQKTNNX-UHFFFAOYSA-N | Cc4cccc5C(CC1C(=O)N(CCCCCN)C(=O)N1S(=O)(=O)c2cc(C(=O)N(C)C)c(cc2)Oc3cc(Cl)c(O)cc3)=CNc45 | SSTR2 agonist | SSTR2 | 37 |
| UKMJWGFHXMGRNG-VZUYHUTRSA-O | CCCC[P+](CCCC)(CCCC)Cc1ccc(cc1)NC(=O)[C@@H](Cc3ccc2ccccc2c3)N/C(/NC4CCCCC4)=N/C5CCCCC5 | BDKRB2 antagonist | BDKRB2 | 35 |
| WRWCAQNPEXYGJK-PKNBQFBNSA-N | CC2(C)CCC(C)(C)c1cc(ccc12)C3CCCC4OC(/C=C/C(O)=O)=CC3=4 | Retinoid RXRalpha Agonists | RXRA | 38 |
| SBOKKVUBLNZTCT-OUKQBFOZSA-N | Oc3nc1ccc(Cl)cc1c(c2ccccc2)c3C(=O)/C=C/C5Nc4ccccc4N=5 | Apoptosis Inhibitors;PKB alpha/Akt1 Inhibitors | AKT1 | 39 |
| FIVPIPIDMRVLAY-RBJBARPLSA-N | CN3C(=O)[C@]24CC1=CC=C[C@H](O)[C@H]1N2C(=O)[C@@]3(CO)SS4 | NF-kappaB (NFKB) Activation Inhibitors | SUV39H1 | 37 |
| MPUQHZXIXSTTDU-QXGSTGNESA-N | NS(=O)(=O)OC[C@@H]1C[C@H](C[C@@H]1O)N5C=Cc4c(N[C@H]2CCc3ccccc23)ncnc45 | NEDD8-Activating Enzyme (NAE) Inhibitors | NAE1,UBA3 | 41 |
| UBPYILGKFZZVDX-UHFFFAOYSA-N | COc4cc(Nc2c1cc(OC)c(cc1ncc2C#N)OCCCN3CCN(C)CC3)c(Cl)cc4Cl | Abl Kinase Inhibitors;Apoptosis Inducers;Bcr-Abl Kinase Inhibitors;Inhibitors of Signal Transduction Pathways;Src Kinase Inhibitors;STAT-5 Inhibitors | FYN | 44 |
| OJCKRNPLOZHAOU-MNKIFKDHSA-N | C[C@H]1C[C@@H](C)C[C@H](C)[C@@H](O)/C(/C#N)=C/C=C\C[C@H](OC(=O)C[C@H](O)[C@@H](C)C1)[C@@H]2CCC[C@H]2C(O)=O | Translation inhibitor;TARS inhibitor;angiogenesis inhibitor | TARS1 | 39 |
| VRQMAABPASPXMW-HDICACEKSA-N | COc4cc(CCC3C=C(NC(=O)c1ccc(cc1)N2C[C@@H](C)N[C@@H](C)C2)NN=3)cc(c4)OC | FGFR1 gene inhibitor | FGFR1 | 37 |

| Inchi-key | SMILES | MOA | GeneIDs | Median adj. cone survival (%) |
|---|---|---|---|---|
| VUVUVNZRUGEAHB-CYBMUJFWSA-N | COc1cc4c(cc1C2C(C)=NOC=2C)ncc5NC(=O)N([C@H](C)c3ccccn3)c45 | BRD2/3/4/T BET familiy inhibitor;BRD4 gene inhibitor | BRD4 | 39 |
| GXJABQQUPOEUTA-RDJZCZTQSA-N | CC(C)C[C@H](NC(=O)[C@H](Cc1ccccc1)NC(=O)c2cnccn2)B(O)O | Proteasome inhibitor;Proteasome Inhibitors;Apoptosis Inducers;Caspase 3 Activators;NF-kappaB (NFKB) Activation Inhibitors | CASP3,CTRB1,PSMA1,PSMA2,PSMA3,PSMA4,PSMA5,PSMA6,PSMA7,PSMB1,PSMB2,PSMB3,PSMB4,PSMB5,PSMB6,PSMB7,PSMB8,PSMB9,PSMB10,PSMC3,PSMC5,PSMD1,PSMD2,PSMD3,PSMD4,PSMD7,PSMD8,PSMD11,PSMD13,CTRC,PSMB11,PSMA8 | 39 |
| RGFKZORXYSZRTQ-UHFFFAOYSA-N | Cc2nc(NC(N)=N)nc3ccc1ccccc1c23 | | NPY1R,NPFFR1 | 34 |
| HVXBOLULGPECHP-WAYWQWQTSA-N | COc2ccc(/C=C\c1cc(OC)c(OC)c(c1)OC)cc2O | Microtubule Polymerization Inhibitors | TUBA4A,TUBB2A,TUBA1A,TUBA1B,TUBB3,TUBB4A,TUBB4B,TUBB1,TUBB6,TUBA1C,TUBA3E,TUBA3D,TUBB8,TUBB2B,TUBA3C,TUBG1,TUBG2,TUBD1,TUBA8 | 39 |
| JCTUPGHJPIIZOK-UHFFFAOYSA-N | O=C(Nc1ccccc1)Nc2ccc(cc2)OC(F)(F)F | P2Y1 antagonist | P2RY1 | 34 |
| OGHNVEJMJSYVRP-UHFFFAOYSA-N | COc1ccccc1OCCNCC(O)COc3cccc4Nc2ccccc2c34 | beta1-Adrenoceptor Antagonists | ADRB1 | 38 |
| WWVANQJRLPIHNS-ZKWXMUAHSA-N | N=C2N[C@H]1CS[C@@H](CCCCC(O)=O)[C@H]1N2 | Inducible Nitric Oxide Synthase (NOS-2) Inhibitors;Neuronal Nitric Oxide Synthase Inhibitors | NOS1,NOS2 | 52 |
| AECDBHGVIIRMOI-UHFFFAOYSA-N | Nc1ncnc2c1C(=CN2C4CC(CN3CCC3)C4)c5cccc(c5)OCc6ccccc6 | MTH1 inhibitor;IGF1R inhibitor | IGF1R,NUDT1 | 42 |
| LGGNGJGQVBUVOE-UHFFFAOYSA-N | C#CCOCCOCCOCCOCCOc1ccc(cc1)c2ccc(cc2)OS(=O)(=O)F | CRABP2 inhibitor | CRABP2 | 38 |
| JQUJTHQSFVWZEH-UHFFFAOYSA-N | CSC2SC(C1C=CN(N=1)C(C)=O)=C3CC(C)(C)CC(=O)C=23 | | GABRA5 | 35 |
| WAEXFXRVDQXREF-UHFFFAOYSA-N | ONC(=O)CCCCCCC(=O)Nc1ccccc1 | HDACs;Apoptosis Inducers;Histone Deacetylase 1 (HDAC1) Inhibitors;Histone Deacetylase 2 (HDAC2) Inhibitors;Histone Deacetylase 6 (HDAC6) Inhibitors;Histone Deacetylase 3 (HDAC3) Inhibitors;Inhibitor;"HDACs;Apoptosis Inducers;Histone Deacetylase 1 (HDAC1) Inhibitors;Histone Deacetylase 2 (HDAC2) Inhibitors;Histone Deacetylase 6 (HDAC6) Inhibitors;Histone Deacetylase 3 (HDAC3) Inhibitors;Inhibitor" | HDAC1,HDAC2,HDAC3,HDAC9,HDAC6,HDAC5,HDAC7,HDAC8,HDAC11,HDAC10,STAT3 | 35 |
| GWHSPAGSUKYSEY-UHFFFAOYSA-N | OC(=O)c1ccc(cc1)C2=NC(=CS2)c3ccc(c3)C(F)(F)F | Retinoid RXRalpha Agonists;Retinoid RARalpha Agonists | RARA,RXRA | 34 |
| PTJGLFIIZFVFJV-UHFFFAOYSA-N | ONC(=O)CCCCCCC(=O)Nc1cccnc1 | Histone Deacetylase (HDAC) Inhibitors | HDAC9,HDAC5,HDAC7,HDAC8,HDAC11,HDAC10 | 43 |
| WDPFJWLDPVQCAJ-UHFFFAOYSA-N | CCN(CC)CCN(Cc1ccc(cc1)c2ccc(cc2)C(F)(F)F)C(=O)CN4C(=NC(=O)C3CCCC=34)SCc5ccc(F)cc5 | Lipoprotein Associated Phospholipase A2 (Lp-PLA2) Inhibitors | PLA2G7,OPTN | 48 |
| TYLTZPAGUBOPCU-UHFFFAOYSA-N | Cc1ccccc1N3C(=O)c2c(cccc2C)N=C3CN6N=C(c4ccc(O)c(F)c4)c5c(N)ncnc56 | Phosphatidylinositol 3-Kinase delta (PI3Kdelta) Inhibitors;Phosphatidylinositol 3-Kinase gamma (PI3Kgamma) Inhibitors | PIK3CD,PIK3CG | 41 |
| AJFGLTPLWPTALJ-SSDOTTSWSA-N | N[C@@](CF)(CC1=CNC=N1)C(O)=O | Histidine Decarboxylase Inhibitors | HDC | 37 |

| Inchi-key | SMILES | MOA | GeneIDs | Median adj. cone survival (%) |
|---|---|---|---|---|
| RPGDCRNUJYFGLT-UHFFFAOYSA-N | CC(CN)C1=CNc2ccc(O)cc12 | | HTR1F | 34 |
| LCNDUGHNYMJGIW-UHFFFAOYSA-N | CN2C(=O)C1C(=NOC=1C=C2c3ccccc3)c4ccccc4 | mgluR7 Antagonists | GRM7 | 42 |
| XQVVPGYIWAGRNI-JOCHJYFZSA-N | CC[C@@H]4C(=O)N(C)c3cnc(Nc1ccc(cc1OC)C(=O)NC2CCN(C)CC2)nc3N4C5CCCC5 | Antimitotic Drugs;Polo-like Kinase-1 (Plk-1) Inhibitors | PLK1,BRD4 | 46 |
| HHOVRZGUSBMKKU-ZDUSSCGKSA-N | CNc1nccc(n1)C2=CC=C(S2)C(=O)N[C@H](CN)Cc3ccc(Cl)cc3Cl | PKA and AKT (a.k.a. PKB) | AKT1 | 36 |
| MRBBFOWSPXHYQT-FCXRPNKRSA-N | COc3cc(/C=C/C2=CC(\C=C\c1ccc(O)c(c1)OC)=NO2)ccc3O | beta-Amyloid (Abeta) Aggregation Inhibitors;Cyclooxygenase-1 Inhibitors;Cyclooxygenase-2 Inhibitors;Free Radical Scavengers | APP,PTGS1,PTGS2 | 45 |
| PPLNRTPNYCWODC-UHFFFAOYSA-N | Oc3cc2Nc1ccccc1c2cc3C(=O)Nc4ccc(Cl)cc4 | Carboxylesterase Inhibitors | CES2 | 36 |
| QMCXVSJLYBVIHV-VXGBXAGGSA-N | C[C@@H]1CC(C[C@@H](C)N1)Nc2nccc(n2)C3=CNc4c(F)cccc34 | IKBKB inhibitor | IKBKB | 41 |
| VRYZCEONIWEUAV-UHFFFAOYSA-N | Cc1cc(C)cc(c1)C(=O)NOCCCCCC(=O)NO | HDAC 4/5 | HDAC4,HDAC5 | 40 |
| RTKIYFITIVXBLE-WKWSCTOISA-N | C/C(/C=C/C(=O)NO)=C\C(C)C(=O)c1ccc(cc1)N(C)C | HDAC inhibitor | HDAC1,HDAC3,HDAC4,HDAC6,HDAC10 | 21 |
| TZZISDZOBASIEO-UHFFFAOYSA-N | CC(C)N4C=C(c2ccnc(Nc1cc(ccc1C)C(N)=O)n2)c3ccncc34 | NFkappaB-inducing kinase (NIK;MAP3K14) Inhibitors | MAP3K14 | 44 |

***Table S1:*** Compound summary of all significant cone-damaging compounds of the primary screen.

### Table S2

| MOA-name | Structure | Inchi-key | Median cone survival (%) | | | | Significant at any concentration |
|---|---|---|---|---|---|---|---|
| | | | 0.01 µM | 0.1 µM | 1 µM | 10 µM | |
| **Sodium ionophore** | 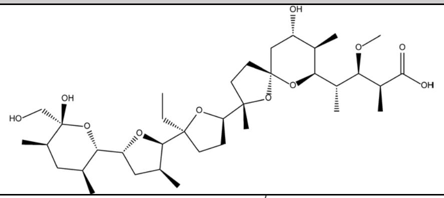 | GAOZTHIDHYLHMS-KEOBGNEYSA-N | 99 | 34 | 31 | 8 | * |
| **ITK Kinase inhibitor** | 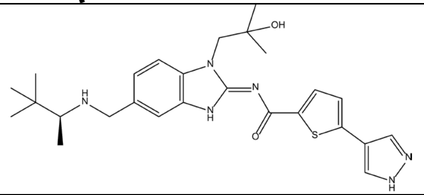 | MUAICZWSFWUFNA-INIZCTEOSA-N | 100 | 92 | 91 | 14 | * |
| **BCL2L1 inhibitor** | 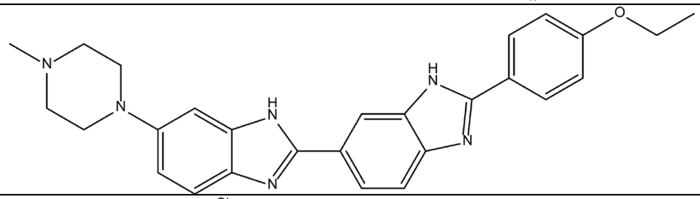 | PRDFBSVERLRRMY-UHFFFAOYSA-N | 104 | 111 | 64 | 14 | * |
| **AP-1 inhibitor** | 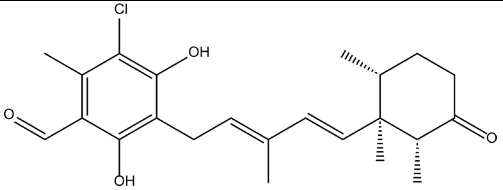 | SETVRSKZJJWOPA-FLDGXQSCSA-N | 105 | 111 | 14 | 29 | * |
| **Immunoproteasome inhibitor** | 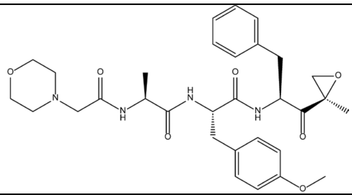 | WQAVPPWWLLVGFK-VTNASVEKSA-N | 101 | 106 | 53 | 11 | * |
| **Beta-Catenin inhibitor** | 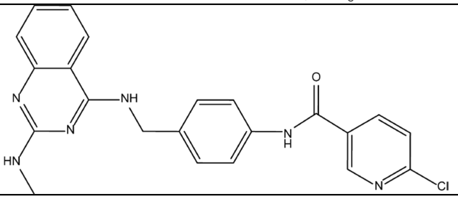 | PIROHBFQHMKAPE-UHFFFAOYSA-N | 106 | 92 | 98 | 21 | * |

| MOA-name | Structure | Inchi-key | Median cone survival (%) | | | | Significant at any concentration |
|---|---|---|---|---|---|---|---|
| | | | 0.01 µM | 0.1 µM | 1 µM | 10 µM | |
| **HDAC inhibitor 3** | 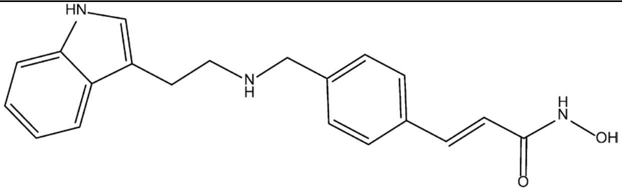 | ZLUZDKXBTNQWOL-MDZDMXLPSA-N | 93 | 68 | 52 | 22 | * |
| **AAA ATPase p97 inhibitor** | 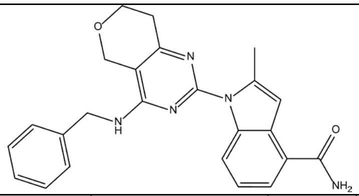 | RDALZZCKQFLGJP-UHFFFAOYSA-N | 104 | 102 | 25 | 26 | * |
| **CDK1-2 inhibitor** | 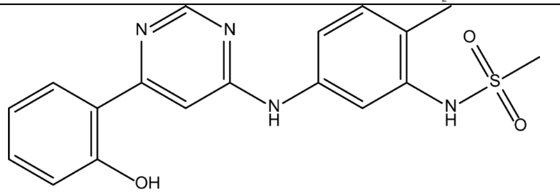 | HPTMTYKYKUKYCQ-UHFFFAOYSA-N | 101 | 90 | 120 | 22 | * |
| **Staurosporine** | 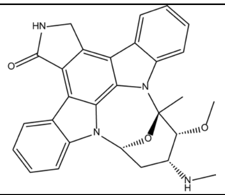 | HKSZLNNOFSGOKW-FYTWVXJKSA-N | 73 | 89 | 69 | 24 | * |
| **PAM of mAChR** | 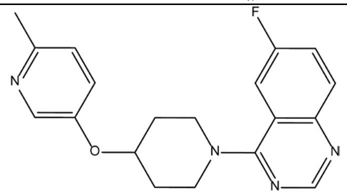 | SMYUEWYXIKCMEA-UHFFFAOYSA-N | 95 | 106 | 105 | 25 | * |
| **Inhibitor of mitochondrial ATPase** | 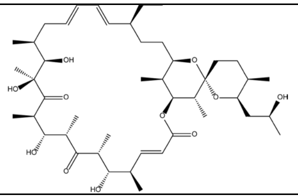 | MNULEGDCPYONBU-WMBHJXFZSA-N | 92 | 32 | 58 | 58 | * |

| MOA-name | Structure | Inchi-key | Median cone survival (%) | | | | Significant at any concentration |
|---|---|---|---|---|---|---|---|
| | | | 0.01 µM | 0.1 µM | 1 µM | 10 µM | |
| **HDAC inhibitor 1** | 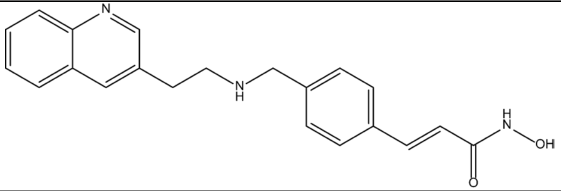 | NLKQTNGJKCBXMU-MDZDMXLPSA-N | 91 | 85 | 57 | 32 | * |
| **HDAC inhibitor 6** | 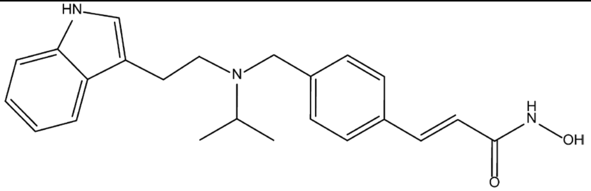 | ILAZEUQGANOTAY-VAWYXSNFSA-N | 91 | 63 | 70 | 33 | * |
| **BIRC5 gene inhibitor** | 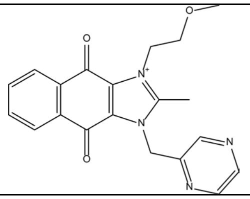 | OTSOOHRUMBRSHZ-UHFFFAOYSA-N | 92 | 72 | 66 | 31 | * |
| **FLT3 inhibitor** | 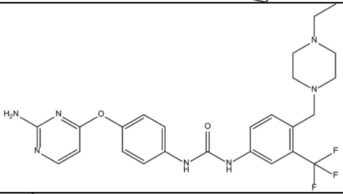 | VQQRBBFRJRBWPF-UHFFFAOYSA-N | 104 | 94 | 90 | 31 | * |
| **HDAC inhibitor 4** | 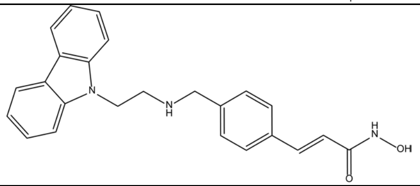 | ATSUJKLLCSXMLV-BUHFOSPRSA-N | 93 | 78 | 44 | 36 | * |
| **Complex I inhibitors** | 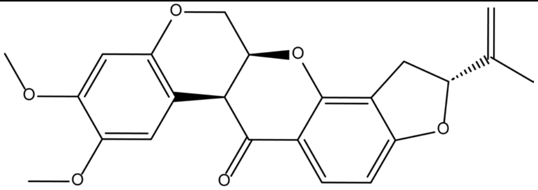 | JUVIOZPCNVVQFO-HBGVWJBISA-N | 104 | 39 | 81 | 60 | * |

| MOA-name | Structure | Inchi-key | Median cone survival (%) | | | | Significant at any concentration |
|---|---|---|---|---|---|---|---|
| | | | 0.01 µM | 0.1 µM | 1 µM | 10 µM | |
| **LPA 2 receptor antagonist** | 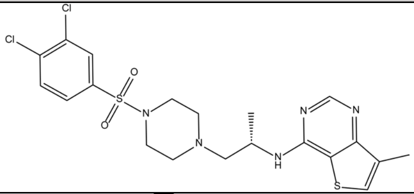 | BPRNMVDTWIHULJ-AWEZNQCLSA-N | 104 | 109 | 98 | 20 | * |
| **HDAC inhibitor 5** | 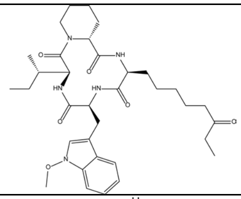 | JWOGUUIOCYMBPV-GMFLJSBRSA-N | 100 | 95 | 80 | 36 | * |
| **Anxiolytic** | 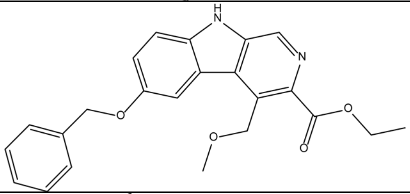 | ALBKMJDFBZVHAK-UHFFFAOYSA-N | 103 | 105 | 105 | 21 | * |
| **MAP3K12 inhibitor** | 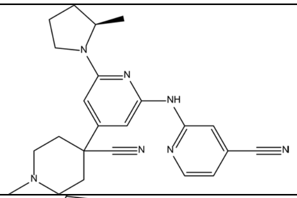 | SXHHURZUIGDYAR-QGZVFWFLSA-N | 89 | 111 | 111 | 19 | * |
| **HDAC inhibitor 2** | 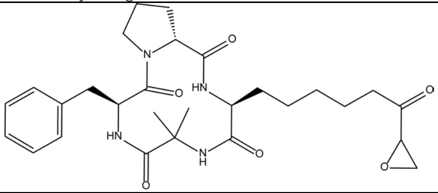 | SGYJGGKDGBXCNY-ZDIDWYTNSA-N | 85 | 66 | 68 | 49 | * |
| **Top1-TDP1 inhibitor** | 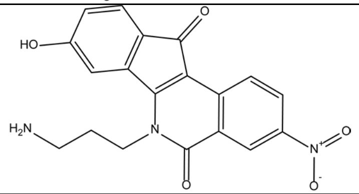 | OKJDLRMQHZRYOZ-UHFFFAOYSA-N | 108 | 98 | 99 | 43 | * |

| MOA-name | Structure | Inchi-key | Median cone survival (%) | | | | Significant at any concentration |
|---|---|---|---|---|---|---|---|
| | | | 0.01 µM | 0.1 µM | 1 µM | 10 µM | |
| **CDK1 inhibitor** | 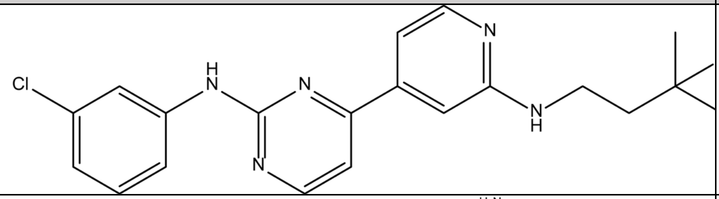 | AAAAA | 111 | 101 | 91 | 39 | * |
| **HDAC inhibitor 7** | 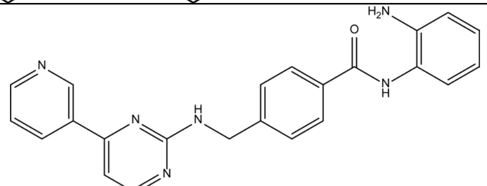 | HRNLUBSXIHFDHP-UHFFFAOYSA-N | 98 | 101 | 90 | 57 | * |
| **Antimitotic Agent** | 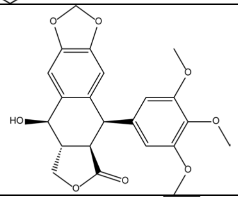 | YJGVMLPVUAXIQN-XVVDYKMHSA-N | 112 | 79 | 49 | 72 | * |
| **PDE5A inhibitor** | 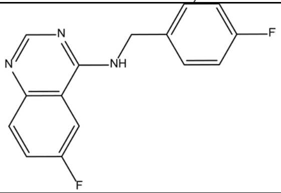 | AWIVHRPYFSSVOG-UHFFFAOYSA-N | 98 | 108 | 112 | 41 | * |
| **PKD inhibitor** | 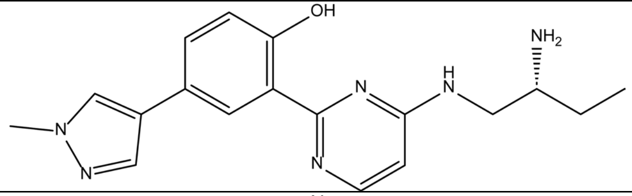 | SCJXQZZYGYLKJG-CQSZACIVSA-N | 95 | 103 | 99 | 62 | * |
| **Tau Aggregation inhibitor** | 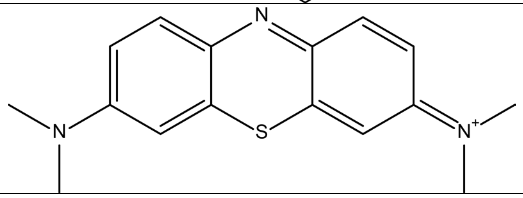 | RBTBFTRPCNLSDE-UHFFFAOYSA-N | 105 | 107 | 96 | 79 | |

| MOA-name | Structure | Inchi-key | Median cone survival (%) | | | | Significant at any concentration |
|---|---|---|---|---|---|---|---|
| | | | 0.01 µM | 0.1 µM | 1 µM | 10 µM | |
| **OPRL1-OPRM1- gene inhibitor** | 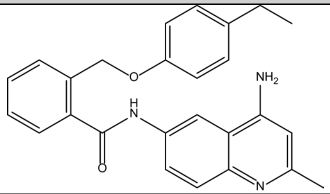 | VTGBZWHPJFMTKS-UHFFFAOYSA-N | 105 | 106 | 107 | 84 | |
| **Chemokine CCR1 antagonist** | 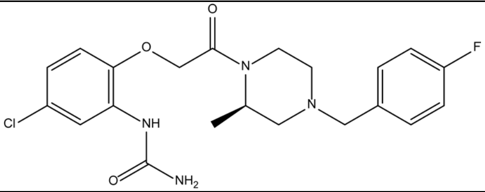 | XQYASZNUFDVMFH-CQSZACIVSA-N | 115 | 106 | 124 | 85 | |
| **Antimitotic drug** | 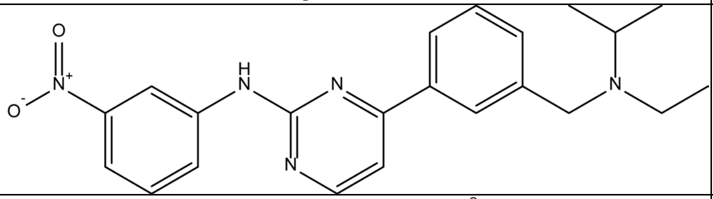 | QPKKAIMODKGHRH-UHFFFAOYSA-N | 96 | 111 | 110 | 83 | |
| **PDE10A inhibitor** | 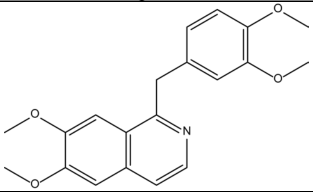 | XQYZDYMELSJDRZ-UHFFFAOYSA-N | 107 | 114 | 99 | 78 | |

***Table S2:*** Compound summary of all retested cone-damaging compounds in the secondary screen.

### Table S3

| MOA-name | Structure | Inchi-key | Median adj. cone survival (%) | | | | Significant at any concentration |
|---|---|---|---|---|---|---|---|
| | | | 0.01 µM | 0.1 µM | 1 µM | 10 µM | |
| **HSP90 inhibitor 2** | 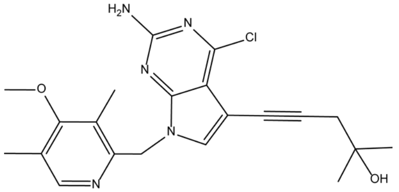 | VOASEWXFCTZRDF-UHFFFAOYSA-N | 67 | 78 | 84 | 71 | * |
| **PDGFR inhibitor 1** | 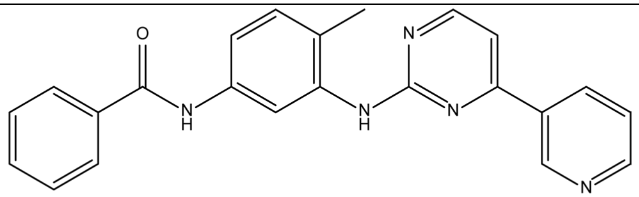 | UOEJSOXEHKCNAE-UHFFFAOYSA-N | 58 | 47 | 54 | 81 | * |
| **HSP90 inhibitor 1** | 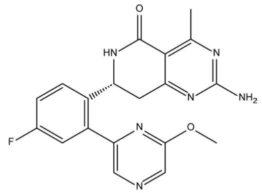 | XRFHWSYKRFEPRA-CYBMUJFWSA-N | 70 | 71 | 80 | 78 | * |
| **MTOR inhibitor 1** | 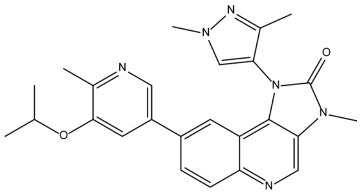 | YWDAJLXJSLKKPA-UHFFFAOYSA-N | 61 | 67 | 61 | 76 | * |
| **MTOR inhibitor 2** | 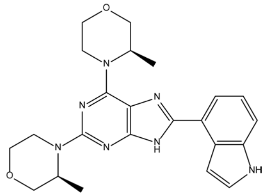 | BYPBFDASESWSQG-CABCVRRESA-N | 59 | 49 | 47 | 72 | |
| **Jak1-2 inhibitor** | 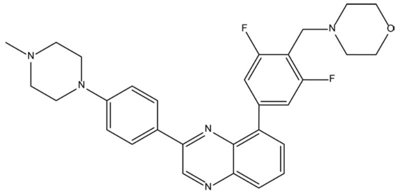 | LBZZVBOHWAZYRL-UHFFFAOYSA-N | 59 | 72 | 55 | 56 | |

| MOA-name | Structure | Inchi-key | Median adj. cone survival (%) | | | | Significant at any concentration |
|---|---|---|---|---|---|---|---|
| | | | 0.01 µM | 0.1 µM | 1 µM | 10 µM | |
| **TP53 expression enhancer** | 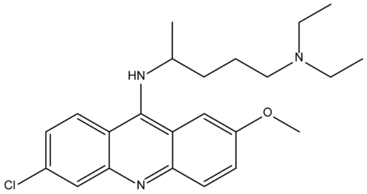 | GPKJTRJOBQGKQK-UHFFFAOYSA-N | 63 | 49 | 70 | 61 | |
| **IMPDH inhibitor** | 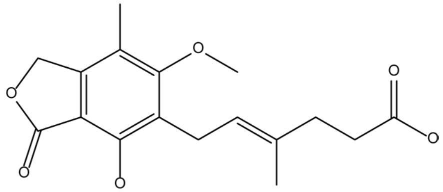 | HPNSFSBZBAHARI-RUDMXATFSA-N | 59 | 50 | 66 | 69 | |
| **Flt3-PKC-inhibitor** | 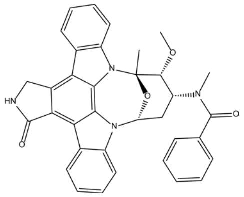 | BMGQWWVMWDBQGC-IIFHNQTCSA-N | 51 | 60 | 52 | 67 | |
| **GPR43 antagonist** | 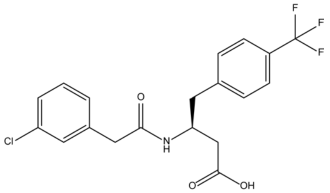 | QOSIJVVNNGXEKE-INIZCTEOSA-N | 51 | 54 | 67 | 53 | |
| **Lipoxygenase inhibitors** | 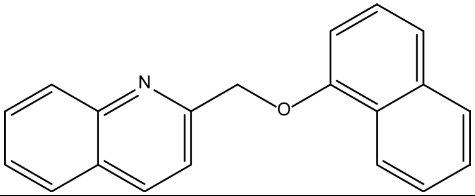 | NZLDBNPKNBCGEN-UHFFFAOYSA-N | 54 | 59 | 58 | 67 | |
| **HSP90 gene modulator** | 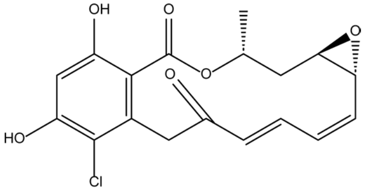 | WYZWZEOGROVVHK-GTMNPGAYSA-N | 60 | 58 | 65 | 56 | |

| MOA-name | Structure | Inchi-key | Median adj. cone survival (%) | | | | Significant at any concentration |
|---|---|---|---|---|---|---|---|
| | | | 0.01 µM | 0.1 µM | 1 µM | 10 µM | |
| **EIF4A3 inhibitor** | 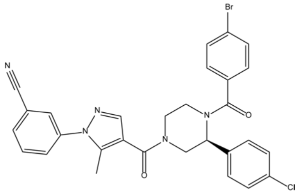 | BDGKKHWJYBQRIE-HHHXNRCGSA-N | 52 | 36 | 50 | 64 | |
| **PI3Kalpha inhibitor** | 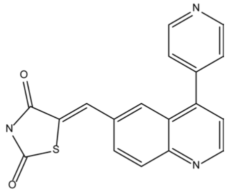 | QDITZBLZQQZVEE-YBEGLDIGSA-N | 58 | 58 | 49 | 63 | |
| **GGPS1 inhibitor** | 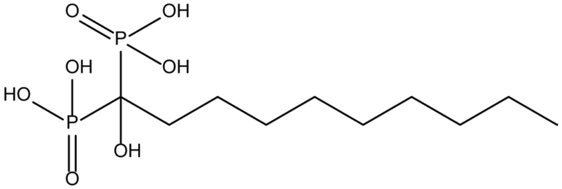 | NSCPCMXKWUFFNL-UHFFFAOYSA-N | 58 | 55 | 51 | 63 | |
| **PKC inhibitor** | 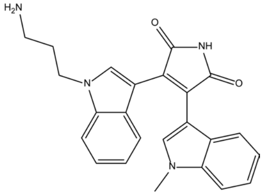 | UQHKJRCFSLMWIA-UHFFFAOYSA-N | 53 | 62 | 60 | 57 | |
| **AP-1-tryptase-MATE1 inhibitor** | 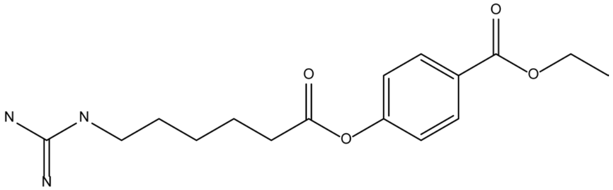 | YKGYIDJEEQRWQH-UHFFFAOYSA-N | 62 | 58 | 54 | 62 | |
| **Cyclooxygenase inhibitor** | 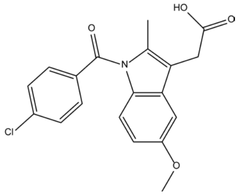 | CGIGDMFJXJATDK-UHFFFAOYSA-N | 58 | 60 | 46 | 62 | |

| MOA-name | Structure | Inchi-key | Median adj. cone survival (%) | | | | Significant at any concentration |
|---|---|---|---|---|---|---|---|
| | | | 0.01 µM | 0.1 µM | 1 µM | 10 µM | |
| **HSP90 inhibitor 3** | 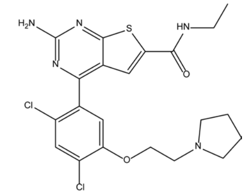 | WJUNQSYQHHIVFX-UHFFFAOYSA-N | 59 | 61 | 54 | 62 | |
| **Protease inhibitor** | 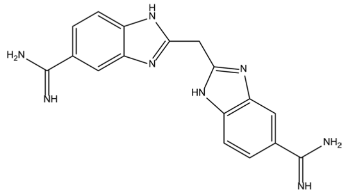 | QZKOOEFIMWKZPK-UHFFFAOYSA-N | 57 | 55 | 49 | 62 | |
| **Elastase inhibitor** | 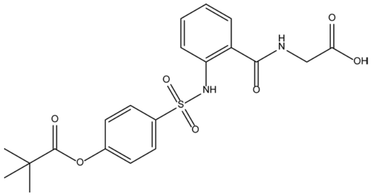 | BTGNGJJLZOIYID-UHFFFAOYSA-N | 59 | 61 | 55 | 60 | |
| **11beta-HSD1 inhibitor** | 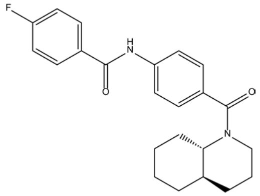 | WSUXACNDZSQZBC-IERDGZPVSA-N | 55 | 54 | 61 | 56 | |
| **CDK9 inhibitor** | 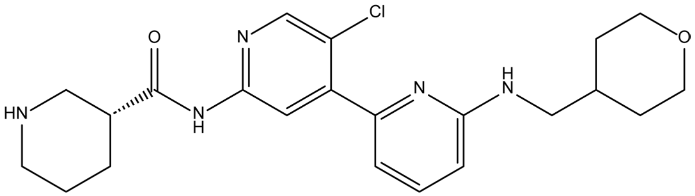 | PHESISZXAITBIM-MRXNPFEDSA-N | 59 | 60 | 52 | 59 | |
| **Covalent necroptosis inhibitor** | 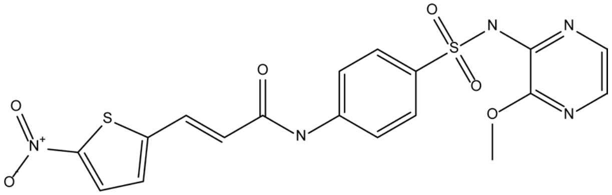 | FNPPHVLYVGMZMZ-XBXARRHUSA-N | 51 | 56 | 60 | 56 | |

| MOA-name | Structure | Inchi-key | Median adj. cone survival (%) | | | | Significant at any concentration |
|---|---|---|---|---|---|---|---|
| | | | 0.01 µM | 0.1 µM | 1 µM | 10 µM | |
| **AhR agonist** | 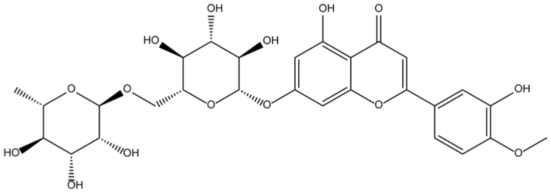 | GZSOSUNBTXMUFQ-YFAPSIMESA-N | 55 | 55 | 60 | 52 | |
| **ACK inhibitor** | 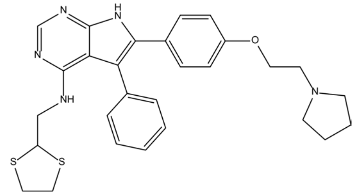 | ZJLXSIPWULUCRJ-UHFFFAOYSA-N | 52 | 42 | 39 | 59 | |
| **NMDA glycine-site antagonist** | 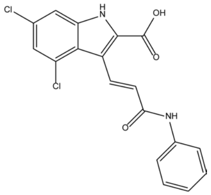 | WZBNEZWCNKUOSM-VOTSOKGWSA-N | 58 | 50 | 49 | 48 | |
| **IL-5R antagonists** | 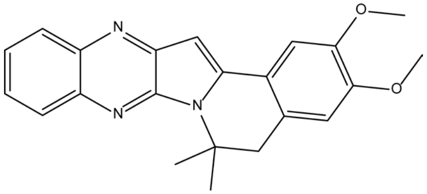 | HIXSPVQXXDULHS-UHFFFAOYSA-N | 55 | 50 | 56 | 53 | |
| **IGF-1R inhibitor** | 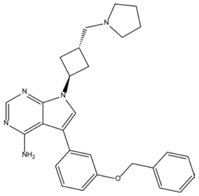 | LSFLAQVDISHMNB-AFARHQOCSA-N | 50 | 51 | 54 | 56 | |
| **Receptor tyrosine kinase inhibitor** | 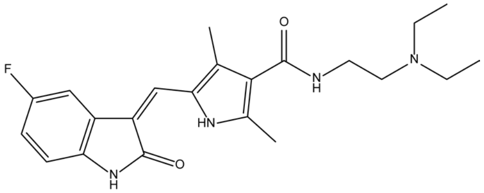 | WINHZLLDWRZWRT-ATVHPVEESA-N | 53 | 49 | 54 | 50 | |

| MOA-name | Structure | Inchi-key | Median adj. cone survival (%) | | | | Significant at any concentration |
|---|---|---|---|---|---|---|---|
| | | | 0.01 µM | 0.1 µM | 1 µM | 10 µM | |
| **Lymphangiogenesis inducer** | 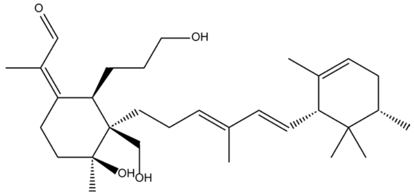 | CUWJDZXEDIUEEW-HLFHJSLSSA-N | 42 | 47 | 44 | 50 | |

***Table S3:*** Compound summary of all retested cone-saving compounds in the secondary screen.

### Table S4

| Kinase Name | GeneID | kinase activity, % ctrl | | | |
|---|---|---|---|---|---|
| | | CS-KI-1 | CS-KI-1A | CS-KI-2 | CS-KI-2A |
| ABL1 | ABL1 | 101 | 104 | 71 | 14 |
| ABL2 | ABL2 | 116 | 115 | 80 | 30 |
| ACK1 | TNK2 | 91 | 100 | 90 | 104 |
| ACVR1 | ACVR1 | 97 | 83 | 92 | 95 |
| ACVR1B | ACVR1B | 104 | 101 | 96 | 120 |
| ACVR2A | ACVR2A | 90 | 86 | 94 | 109 |
| ACVR2B | ACVR2B | 138 | 114 | 83 | 118 |
| ACVRL1 | ACVRL1 | 98 | 121 | 97 | 105 |
| AKT1 | AKT1 | 116 | 113 | 106 | 123 |
| AKT2 | AKT2 | 110 | 137 | 126 | 187 |
| AKT3 | AKT3 | 236 | 119 | 112 | 125 |
| ALK | ALK | 49 | 85 | 81 | 95 |
| AMPKalpha1 | PRKAA1 | 98 | 76 | 87 | 88 |
| ARAF YDYD | ARAF | 107 | 87 | 68 | 71 |
| ARK5 | NUAK1 | 102 | 108 | 102 | 105 |
| ASK1 | MAP3K5 | 125 | 108 | 107 | 115 |
| AuroraA | AURKA | 112 | 92 | 96 | 103 |
| AuroraB | AURKB | 79 | 100 | 94 | 92 |
| AuroraC | AURKC | 102 | 94 | 80 | 89 |
| AXL | AXL | 71 | 80 | 82 | 85 |
| BLK | PTK6 | 79 | 85 | 82 | 66 |
| BMPR1A | BMPR1A | 112 | 97 | 105 | 90 |
| BMPR1B | BMPR1B | 117 | 102 | 112 | 119 |
| BMX | BMX | 106 | 104 | 92 | 94 |
| BRAF | BRAF | 126 | 91 | 59 | 67 |
| BRK | BRK | 105 | 99 | 92 | 91 |
| BRSK1 | BRSK1 | 112 | 101 | 103 | 112 |
| BRSK2 | BRSK2 | 117 | 109 | 97 | 108 |
| BTK | BTK | 89 | 94 | 83 | 80 |
| BUB1B | BUB1B | 108 | 119 | 90 | 121 |
| CAMK1D | CAMK1D | 113 | 125 | 112 | 117 |
| CAMK2A | CAMK2A | 98 | 111 | 88 | 97 |
| CAMK2B | CAMK2B | 94 | 97 | 95 | 113 |
| CAMK2D | CAMK2D | 106 | 105 | 98 | 103 |
| CAMK2G | CAMK2G | 141 | 122 | 113 | 114 |
| CAMK4 | CAMK4 | 120 | 119 | 111 | 123 |
| CAMKK1 | CAMKK1 | 79 | 93 | 88 | 105 |
| CAMKK2 | CAMKK2 | 42 | 103 | 78 | 95 |
| CDC42BPA | CDC42BPA | 143 | 123 | 119 | 128 |
| CDC42BPB | CDC42BPB | 106 | 83 | 86 | 84 |
| CDC7/DBF4 | CDC7 | 106 | 96 | 87 | 102 |
| CDK1/CycA2 | CDK1 | 117 | 107 | 113 | 92 |
| CDK1/CycB1 | CDK1 | 104 | 102 | 102 | 90 |
| CDK1/CycE1 | CDK1 | 130 | 131 | 132 | 121 |
| CDK10/CycQ | CDK10 | 65 | 94 | 88 | 106 |
| CDK11B/CycK | CDK11B | 97 | 106 | 105 | 103 |
| CDK12/CycK | CDK12 | 121 | 107 | 123 | 109 |
| CDK13/CycK | CDK13 | 84 | 87 | 85 | 84 |

| Kinase Name | GeneID | kinase activity, % ctrl | | | |
|---|---|---|---|---|---|
| | | CS-KI-1 | CS-KI-1A | CS-KI-2 | CS-KI-2A |
| CDK14/CycY | CDK14 | 113 | 122 | 104 | 115 |
| CDK15/CycA2 | CDK15 | 103 | 109 | 209 | 112 |
| CDK15/CycB1 | CDK15 | 93 | 94 | 100 | 106 |
| CDK16/CycY | CDK16 | 126 | 114 | 120 | 124 |
| CDK17/p35NCK | CDK17 | 119 | 113 | 112 | 121 |
| CDK18/CycY | CDK18 | 118 | 109 | 106 | 119 |
| CDK19/CycC | CDK19 | 64 | 102 | 77 | 66 |
| CDK2/CycA2 | CDK2 | 131 | 106 | 105 | 117 |
| CDK2/CycD1 | CDK2 | 134 | 112 | 150 | 94 |
| CDK2/CycE1 | CDK2 | 100 | 109 | 115 | 123 |
| CDK20/CycH | CDK20 | 94 | 100 | 102 | 104 |
| CDK20/CycT1 | CDK20 | 88 | 93 | 90 | 90 |
| CDK3/CycC | CDK3 | 116 | 111 | 88 | 99 |
| CDK3/CycE1 | CDK3 | 122 | 113 | 106 | 109 |
| CDK4/CycD1 | CDK4 | 127 | 100 | 101 | 110 |
| CDK4/CycD2 | CDK4 | 117 | 102 | 114 | 114 |
| CDK4/CycD3 | CDK4 | 117 | 109 | 112 | 116 |
| CDK5/p25NCK | CDK5 | 95 | 106 | 99 | 120 |
| CDK5/p35NCK | CDK5 | 96 | 104 | 109 | 104 |
| CDK6/CycD1 | CDK6 | 99 | 105 | 110 | 90 |
| CDK6/CycD2 | CDK6 | 104 | 106 | 100 | 110 |
| CDK6/CycD3 | CDK6 | 109 | 89 | 93 | 95 |
| CDK7/CycH/MAT1 | CDK7 | 108 | 98 | 113 | 106 |
| CDK8/CycC | CDK8 | 46 | 118 | 66 | 46 |
| CDK9/CycK | CDK9 | 93 | 113 | 125 | 103 |
| CDK9/CycT1 | CDK9 | 78 | 89 | 92 | 104 |
| CHK1 | CHEK1 | 109 | 102 | 105 | 100 |
| CHK2 | CHEK2 | 111 | 104 | 89 | 96 |
| CIT 1-450 | CIT | 106 | 84 | 90 | 89 |
| CK1alpha1 | CSNK1A1 | 111 | 110 | 103 | 89 |
| CK1delta | CSNK1D | 107 | 89 | 97 | 107 |
| CK1epsilon | CSNK1E | 89 | 91 | 96 | 100 |
| CK1gamma1 | CSNK1G1 | 2 | 96 | 98 | 103 |
| CK1gamma2 | CSNK1G2 | 105 | 102 | 102 | 102 |
| CK1gamma3 | CSNK1G3 | 101 | 101 | 103 | 86 |
| CK2alpha1 | CSNK2A1 | 102 | 100 | 84 | 80 |
| CK2alpha2 | CSNK2A2 | 94 | 93 | 86 | 80 |
| CLK1 | CLK1 | 80 | 11 | 50 | 79 |
| CLK2 | CLK2 | 80 | 16 | 87 | 84 |
| CLK3 | CLK3 | 91 | 81 | 65 | 82 |
| CLK4 | CLK4 | 74 | 17 | 23 | 59 |
| COT | MAP3K8 | 120 | 125 | 108 | 107 |
| CSF1R | CSF1R | 98 | 100 | 84 | 56 |
| CSK | CSK | 100 | 90 | 145 | 109 |
| DAPK1 | DAPK1 | 106 | 96 | 103 | 100 |
| DAPK2 | DAPK2 | 96 | 90 | 90 | 104 |
| DAPK3 | DAPK3 | 104 | 91 | 98 | 100 |
| DCAMKL2 | DCLK2 | 100 | 96 | 94 | 121 |
| DDR2 | DDR2 | 100 | 106 | 10 | 2 |
| DMPK | DMPK | 72 | 105 | 104 | 100 |
| DNAPK | PRKDC | 1 | 0 | 77 | 75 |

| Kinase Name | GeneID | kinase activity, % ctrl | | | |
|---|---|---|---|---|---|
| | | CS-KI-1 | CS-KI-1A | CS-KI-2 | CS-KI-2A |
| DYRK1A | DYRK1A | 90 | 89 | 87 | 96 |
| DYRK1B | DYRK1B | 114 | 104 | 99 | 100 |
| DYRK2 | DYRK2 | 140 | 106 | 94 | 119 |
| DYRK3 | DYRK3 | 86 | 105 | 129 | 99 |
| DYRK4 | DYRK4 | 113 | 109 | 114 | 101 |
| EEF2K | EEF2K | 123 | 125 | 126 | 122 |
| EGFR | EGFR | 100 | 111 | 90 | 88 |
| EIF2AK2 | EIF2AK2 | 113 | 108 | 95 | 87 |
| EIF2AK3 | EIF2AK3 | 124 | 93 | 119 | 128 |
| EIF2AK4 | EIF2AK4 | 109 | 102 | 107 | 106 |
| EPHA1 | EPHA1 | 93 | 117 | 111 | 107 |
| EPHA2 | EPHA2 | 84 | 76 | 59 | 75 |
| EPHA3 | EPHA3 | 98 | 104 | 93 | 102 |
| EPHA4 | EPHA4 | 70 | 85 | 72 | 89 |
| EPHA5 | EPHA5 | 89 | 88 | 80 | 88 |
| EPHA6 | EPHA6 | 107 | 92 | 85 | 110 |
| EPHA7 | EPHA7 | 102 | 105 | 88 | 107 |
| EPHA8 | EPHA8 | 107 | 106 | 144 | 112 |
| EPHB1 | EPHB1 | 96 | 83 | 68 | 75 |
| EPHB2 | EPHB2 | 101 | 104 | 97 | 103 |
| EPHB3 | EPHB3 | 91 | 99 | 80 | 88 |
| EPHB4 | EPHB4 | 99 | 93 | 81 | 107 |
| ERBB2 | ERBB2 | 89 | 92 | 82 | 90 |
| ERBB4 | ERBB4 | 110 | 112 | 84 | 57 |
| ERK1 | MAPK3 | 115 | 112 | 111 | 122 |
| ERK2 | MAPK1 | 111 | 125 | 119 | 122 |
| ERK5 | MAPK7 | 91 | 91 | 94 | 99 |
| ERK7 | MAPK15 | 108 | 109 | 110 | 125 |
| FAK | PTK2 | 70 | 109 | 93 | 84 |
| FER | FER | 82 | 110 | 99 | 61 |
| FES | FES | 77 | 104 | 87 | 85 |
| FGFR1 | FGFR1 | 101 | 104 | 89 | 115 |
| FGFR2 | FGFR2 | 81 | 72 | 124 | 93 |
| FGFR3 | FGFR3 | 83 | 94 | 94 | 96 |
| FGFR4 | FGFR4 | 96 | 95 | 113 | 112 |
| FGR | FGR | 103 | 100 | 97 | 82 |
| FLT3 | FLT3 | 79 | 114 | 73 | 71 |
| FRK | FRK | 96 | 88 | 70 | 78 |
| FYN | FYN | 109 | 104 | 103 | 85 |
| GRK2 | ADRBK1 | 115 | 94 | 102 | 98 |
| GRK3 | ADRBK2 | 105 | 92 | 106 | 108 |
| GRK4 | GRK4 | 121 | 96 | 102 | 97 |
| GRK5 | GRK5 | 91 | 89 | 98 | 99 |
| GRK6 | GRK6 | 108 | 127 | 146 | 95 |
| GRK7 | GRK7 | 98 | 100 | 107 | 97 |
| GSG2 | HASPIN | 60 | 100 | 94 | 90 |
| GSK3alpha | GSK3A | 87 | 55 | 98 | 99 |
| GSK3beta | GSK3B | 99 | 89 | 99 | 133 |
| HCK | HCK | 95 | 92 | 97 | 88 |
| HIPK1 | HIPK1 | 173 | 109 | 118 | 115 |
| HIPK2 | HIPK2 | 99 | 95 | 89 | 96 |

| Kinase Name | GeneID | kinase activity, % ctrl | | | |
|---|---|---|---|---|---|
| | | CS-KI-1 | CS-KI-1A | CS-KI-2 | CS-KI-2A |
| HIPK3 | HIPK3 | 129 | 129 | 111 | 126 |
| HIPK4 | HIPK4 | 96 | 95 | 69 | 42 |
| HRI | EIF2AK1 | 97 | 105 | 103 | 102 |
| IGF1R | IGF1R | 109 | 95 | 99 | 117 |
| IKKalpha | CHUK | 110 | 104 | 106 | 109 |
| IKKbeta | IKBKB | 108 | 100 | 106 | 113 |
| IKKepsilon | IKBKE | 118 | 105 | 111 | 108 |
| INSR | INSR | 103 | 116 | 115 | 91 |
| INSRR | INSRR | 110 | 111 | 106 | 96 |
| IRAK1 | IRAK1 | 130 | 100 | 90 | 98 |
| IRAK4 | IRAK4 | 139 | 97 | 115 | 99 |
| ITK | ITK | 252 | 91 | 101 | 100 |
| JAK1 | JAK1 | 98 | 112 | 95 | 81 |
| JAK2 | JAK2 | 110 | 118 | 97 | 103 |
| JAK3 | JAK3 | 109 | 125 | 112 | 108 |
| JNK1 | MAPK8 | 102 | 100 | 75 | 71 |
| JNK2 | MAPK9 | 113 | 105 | 88 | 80 |
| JNK3 | MAPK10 | 94 | 86 | 58 | 63 |
| KIT | KIT | 101 | 97 | 40 | 33 |
| LCK | LCK | 100 | 98 | 55 | 15 |
| LIMK1 | LIMK1 | 141 | 149 | 118 | 106 |
| LIMK2 | LIMK2 | 119 | 126 | 110 | 105 |
| LKB1/MO25a/STRADa | STK11 | 103 | 97 | 100 | 99 |
| LRRK2 | LRRK2 | 94 | 101 | 83 | 101 |
| LTK | LTK | 70 | 87 | 89 | 93 |
| LYN | LYN | 106 | 102 | 94 | 47 |
| MAP3K1 | MAP3K1 | 115 | 111 | 108 | 111 |
| MAP3K10 | MAP3K10 | 108 | 108 | 104 | 113 |
| MAP3K11 | MAP3K11 | 88 | 109 | 91 | 97 |
| MAP3K7/MAP3K7IP1 | MAP3K7 | 92 | 85 | 90 | 86 |
| MAP3K9 | MAP3K9 | 96 | 102 | 89 | 101 |
| MAP4K1 | MAP4K1 | 95 | 75 | 98 | 102 |
| MAP4K2 | MAP4K2 | 78 | 95 | 88 | 95 |
| MAP4K4 | MAP4K4 | 89 | 92 | 97 | 101 |
| MAP4K5 | MAP4K5 | 93 | 91 | 92 | 111 |
| MAPKAPK2 | MAPKAPK2 | 95 | 98 | 95 | 100 |
| MAPKAPK3 | MAPKAPK3 | 97 | 92 | 89 | 98 |
| MAPKAPK5 | MAPKAPK5 | 81 | 82 | 84 | 89 |
| MARK1 | MARK1 | 100 | 97 | 101 | 98 |
| MARK2 | MARK2 | 106 | 104 | 102 | 101 |
| MARK3 | MARK3 | 86 | 102 | 92 | 108 |
| MARK4 | MARK4 | 120 | 124 | 99 | 126 |
| MASTL | MASTL | 92 | 88 | 90 | 100 |
| MATK | MATK | 105 | 88 | 92 | 104 |
| MEK1 | MAP2K1 | 138 | 87 | 122 | 201 |
| MEK2 | MAP2K2 | 101 | 113 | 104 | 105 |
| MEK5 | MAP2K5 | 101 | 111 | 115 | 119 |
| MEKK2 | MAP3K2 | 110 | 97 | 100 | 110 |
| MEKK3 | MAP3K3 | 110 | 107 | 101 | 103 |
| MELK | MELK | 35 | 67 | 69 | 56 |
| MERTK | MERTK | 73 | 96 | 107 | 111 |

| Kinase Name | GeneID | kinase activity, % ctrl | | | |
|---|---|---|---|---|---|
| | | CS-KI-1 | CS-KI-1A | CS-KI-2 | CS-KI-2A |
| MET | MET | 80 | 84 | 88 | 92 |
| MINK1 | MINK1 | 84 | 81 | 78 | 104 |
| MKK3 | MAP2K3 | 146 | 134 | 134 | 135 |
| MKK4 | MAP2K4 | 119 | 128 | 125 | 118 |
| MKK6 SDTD | MAP2K6 | 164 | 139 | 134 | 134 |
| MKK7 | MAP2K7 | 120 | 109 | 134 | 117 |
| MKNK1 | MKNK1 | 99 | 188 | 98 | 119 |
| MKNK2 | MKNK2 | 104 | 93 | 86 | 101 |
| MLK4 | MAP3K21 | 102 | 87 | 85 | 85 |
| MST1 | STK4 | 102 | 88 | 93 | 104 |
| MST2 | STK3 | 119 | 99 | 92 | 108 |
| MST3 | STK24 | 94 | 92 | 88 | 90 |
| MST4 | STK26 | 115 | 108 | 115 | 86 |
| MTOR | MTOR | -1 | -3 | 148 | 94 |
| MUSK | MUSK | 71 | 96 | 95 | 98 |
| MYLK | MYLK | 126 | 92 | 84 | 98 |
| MYLK2 | MYLK2 | 122 | 108 | 106 | 105 |
| MYLK3 | MYLK3 | 116 | 89 | 104 | 100 |
| NDR1 | STK38 | 102 | 91 | 86 | 100 |
| NDR2 | STK38L | 113 | 102 | 101 | 144 |
| NEK1 | NEK1 | 101 | 102 | 96 | 102 |
| NEK11 | NEK11 | 112 | 109 | 102 | 93 |
| NEK2 | NEK2 | 96 | 90 | 130 | 84 |
| NEK3 | NEK3 | 71 | 91 | 110 | 100 |
| NEK4 | NEK4 | 93 | 97 | 85 | 94 |
| NEK6 | NEK6 | 104 | 87 | 90 | 115 |
| NEK7 | NEK7 | 107 | 95 | 98 | 107 |
| NEK9 | NEK9 | 125 | 121 | 114 | 121 |
| NIK | MAP3K14 | 126 | 126 | 181 | 82 |
| NLK | NLK | 108 | 119 | 111 | 105 |
| p38alpha | MAPK14 | 105 | 113 | 96 | 113 |
| p38beta | MAPK11 | 118 | 104 | 4 | 142 |
| p38delta | MAPK13 | 115 | 111 | 104 | 97 |
| p38gamma | MAPK12 | 110 | 111 | 108 | 107 |
| PAK1 | PAK1 | 167 | 106 | 91 | 118 |
| PAK2 | PAK2 | 99 | 95 | 92 | 88 |
| PAK3 | PAK3 | 119 | 100 | 101 | 105 |
| PAK4 | PAK4 | 107 | 98 | 85 | 91 |
| PAK6 | PAK6 | 73 | 71 | 84 | 89 |
| PAK7 | PAK7 | 98 | 96 | 101 | 92 |
| PASK | PASK | 106 | 119 | 113 | 102 |
| PBK | PBK | 102 | 75 | 83 | 138 |
| PDGFRalpha | PDGFRA | 127 | 104 | 6 | 10 |
| PDGFRbeta | PDGFRB | 105 | 86 | 41 | 7 |
| PDK1 | PDPK1 | 112 | 97 | 104 | 103 |
| PHKG1 | PHKG1 | 117 | 127 | 119 | 117 |
| PHKG2 | PHKG2 | 102 | 110 | 103 | 110 |
| PIM1 | PIM1 | 71 | 95 | 98 | 91 |
| PIM2 | PIM2 | 105 | 96 | 86 | 101 |
| PIM3 | PIM3 | 115 | 119 | 100 | 99 |
| PKA | PRKACA | 109 | 103 | 107 | 120 |

| Kinase Name | GeneID | kinase activity, % ctrl | | | |
|---|---|---|---|---|---|
| | | CS-KI-1 | CS-KI-1A | CS-KI-2 | CS-KI-2A |
| PKCalpha | PRKCA | 85 | 75 | 101 | 84 |
| PKCbeta1 | PRKCB | 80 | 75 | 87 | 90 |
| PKCbeta2 | PRKCB | 96 | 91 | 86 | 95 |
| PKCdelta | PRKCD | 113 | 96 | 97 | 94 |
| PKCepsilon | PRKCE | 117 | 116 | 149 | 122 |
| PKCeta | PRKCH | 96 | 88 | 95 | 95 |
| PKCgamma | PRKCG | 99 | 97 | 87 | 99 |
| PKCiota | PRKCI | 151 | 136 | 145 | 142 |
| PKCmu | PRKD1 | 98 | 113 | 93 | 99 |
| PKCnu | PRKCQ | 102 | 92 | 95 | 97 |
| PKCtheta | PRKCQ | 96 | 96 | 103 | 102 |
| PKCzeta | PRKCZ | 91 | 86 | 107 | 85 |
| PKMzeta | PRKCL1 | 74 | 84 | 84 | 88 |
| PKN3 | PKN3 | 88 | 109 | 85 | 83 |
| PLK1 | PLK1 | 106 | 102 | 109 | 93 |
| PLK3 | PLK3 | 104 | 96 | 91 | 94 |
| PRK1 | PKN1 | 113 | 118 | 101 | 101 |
| PRK2 | PKN2 | 136 | 111 | 120 | 118 |
| PRKD2 | PRKD2 | 123 | 117 | 117 | 114 |
| PRKG1 | PRKG1 | 95 | 99 | 103 | 106 |
| PRKG2 | PRKG2 | 102 | 94 | 86 | 119 |
| PRKX | PRKX | 101 | 96 | 92 | 105 |
| PYK2 | PTK2B | 118 | 116 | 119 | 106 |
| RAF1 YDYD | RAF1 | 109 | 101 | 5 | 14 |
| RET | RET | 99 | 106 | 109 | 103 |
| RIPK2 | RIPK2 | 71 | 54 | 71 | 81 |
| RIPK4 | RIPK4 | 104 | 89 | 100 | 107 |
| RIPK5 | RIPK5 | 103 | 102 | 113 | 111 |
| ROCK1 | ROCK1 | 111 | 111 | 97 | 103 |
| ROCK2 | ROCK2 | 89 | 81 | 87 | 86 |
| RON | MST1R | 122 | 115 | 101 | 129 |
| ROS | ROS1 | 55 | 118 | 78 | 80 |
| RPS6KA1 | RPS6KA1 | 99 | 93 | 101 | 90 |
| RPS6KA2 | RPS6KA2 | 101 | 99 | 92 | 99 |
| RPS6KA3 | RPS6KA3 | 96 | 92 | 95 | 86 |
| RPS6KA4 | RPS6KA4 | 128 | 117 | 116 | 115 |
| RPS6KA5 | RPS6KA5 | 112 | 119 | 112 | 109 |
| RPS6KA6 | RPS6KA6 | 114 | 126 | 107 | 108 |
| S6K | RPS6KB1 | 114 | 111 | 108 | 98 |
| S6Kbeta | RPS6KB2 | 87 | 92 | 82 | 97 |
| SAK | PLK4 | 82 | 77 | 54 | 68 |
| SGK1 | SGK1 | 105 | 103 | 91 | 104 |
| SGK2 | SGK2 | 101 | 110 | 103 | 106 |
| SGK3 | SGK3 | 114 | 116 | 115 | 122 |
| SIK1 | SIK1 | 97 | 94 | 105 | 100 |
| SIK2 | SIK2 | 105 | 107 | 105 | 93 |
| SIK3 | SIK3 | 97 | 106 | 88 | 107 |
| SLK | STK10 | 93 | 114 | 104 | 106 |
| SNARK | NUAK1 | 90 | 96 | 83 | 100 |
| SNK | NUAK2 | 95 | 114 | 102 | 92 |
| SRC | SRC | 92 | 91 | 97 | 89 |

| Kinase Name | GeneID | kinase activity, % ctrl | | | |
|---|---|---|---|---|---|
| | | CS-KI-1 | CS-KI-1A | CS-KI-2 | CS-KI-2A |
| SRMS | SRMS | 99 | 92 | 99 | 102 |
| SRPK1 | SRPK1 | 104 | 97 | 105 | 103 |
| SRPK2 | SRPK2 | 103 | 112 | 101 | 99 |
| STK17A | STK17A | 106 | 105 | 91 | 74 |
| STK17B | STK17B | 136 | 117 | 104 | 104 |
| STK23 | MST1 | 109 | 93 | 98 | 113 |
| STK25 | STK25 | 91 | 100 | 98 | 92 |
| STK33 | STK33 | 246 | 107 | 172 | 106 |
| STK39 | STK39 | 124 | 127 | 99 | 82 |
| SYK | SYK | 95 | 104 | 88 | 92 |
| TAOK2 | TAOK2 | 105 | 110 | 96 | 50 |
| TAOK3 | TAOK3 | 104 | 96 | 91 | 91 |
| TBK1 | TBK1 | 124 | 107 | 87 | 98 |
| TEC | TEC | 96 | 96 | 163 | 94 |
| TGFBR1 | TGFBR1 | 114 | 101 | 100 | 121 |
| TGFBR2 | TGFBR2 | 91 | 72 | 65 | 85 |
| TIE2 | TEK | 97 | 99 | 98 | 91 |
| TLK1 | TLK1 | 96 | 89 | 80 | 83 |
| TLK2 | TLK2 | 123 | 97 | 96 | 99 |
| TNK1 | TNK1 | 85 | 74 | 43 | 52 |
| TRKA | NTRK1 | 116 | 120 | 104 | 113 |
| TRKB | NTRK2 | 84 | 91 | 96 | 116 |
| TRKC | NTRK3 | 85 | 107 | 96 | 97 |
| TSF1 | DYRK3 | 54 | 86 | 66 | 66 |
| TSK2 | DYRK4 | 114 | 101 | 110 | 109 |
| TSSK1 | TSSK1 | 110 | 101 | 92 | 109 |
| TTBK1 | TTBK1 | 97 | 84 | 78 | 100 |
| TTBK2 | TTBK2 | 114 | 101 | 99 | 104 |
| TTK | TTK | 91 | 98 | 100 | 103 |
| TXK | TXK | 115 | 110 | 98 | 91 |
| TYK2 | TYK2 | 113 | 113 | 113 | 93 |
| TYRO3 | TYRO3 | 93 | 106 | 101 | 96 |
| ULK1 | ULK1 | 113 | 104 | 96 | 91 |
| ULK2 | ULK2 | 104 | 105 | 90 | 89 |
| ULK3 | ULK3 | 106 | 102 | 92 | 95 |
| VEGFR1 | FLT1 | 94 | 94 | 89 | 93 |
| VEGFR2 | KDR | 102 | 95 | 105 | 82 |
| VEGFR3 | FLT4 | 82 | 83 | 108 | 94 |
| VRK1 | VRK1 | 158 | 104 | 104 | 118 |
| VRK2 | VRK2 | 97 | 90 | 111 | 85 |
| WEE1 | WEE1 | 94 | 94 | 95 | 111 |
| WNK1 | WNK1 | 111 | 102 | 126 | 113 |
| WNK2 | WNK2 | 108 | 118 | 78 | 90 |
| WNK3 | WNK3 | 116 | 117 | 110 | 101 |
| YES | YES1 | 86 | 100 | 88 | 77 |
| ZAK | MAP3K20 | 105 | 105 | 94 | 83 |
| ZAP70 | ZAP70 | 91 | 57 | 49 | 58 |

***Table S4:*** Kinase profiling results.

### Table S5

| Compound Name | Structure | Inchi-key | Mean cone survival (%) | | | | Significant at any concentration | |
|---|---|---|---|---|---|---|---|---|
| | | | Cones | | Rods | | | |
| | | | 10 µM | 1 µM | 10 µM | 1 µM | Cones | Rods |
| **CK-1-I 1** | 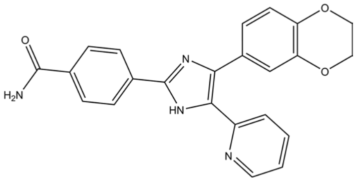 | DPDZHVCKYBCJHW-UHFFFAOYSA-N | 65 | 59 | 75 | 62 | * | * |
| **CK-1-I 2** | 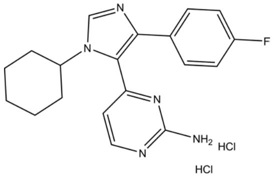 | PSNKGVAXBSAHCH-UHFFFAOYSA-N | 51 | 59 | 62 | 69 | * | * |
| **CK-1-I 3** | 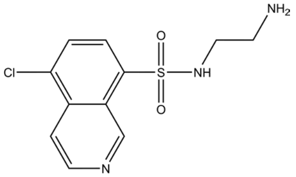 | OGKYMFFYOWUTKV-UHFFFAOYSA-N | 56 | 60 | 48 | 75 | * | * |
| **MAPK11-I 1** | 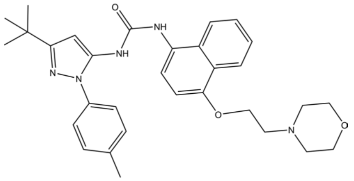 | MVCOAUNKQVWQHZ-UHFFFAOYSA-N | 48 | 51 | 48 | 56 | | |
| **MAPK11-I 2** | 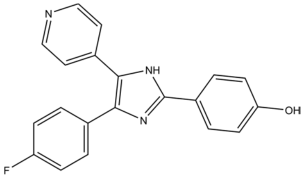 | QHKYPYXTTXKZST-UHFFFAOYSA-N | 59 | 54 | 57 | 47 | * | |
| **MAPK11-I 3** | 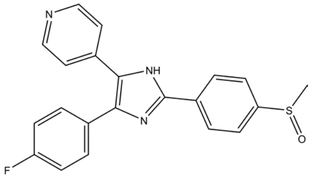 | CDMGBJANTYXAIV-UHFFFAOYSA-N | 57 | 61 | 46 | 68 | * | * |

***Table S5:*** Summary of CK-1 and MAPK11 inhibitors.

### Table S6

| Target | Species | Dilution | Source | Identifier |
|---|---|---|---|---|
| Bassoon | Mouse (monoclonal) | 1:800 | Enzo | SAP7F407 |
| ARR3 | Mouse (monoclonal) | 1:500 | Gift from the Laboratory of Wolfgang Baehr, University of Utah | - |
| NRL | Goat (polyclonal) | 1:500 | R&D systems | AF2945-SP |
| ONECUT2 | Sheep (polyclonal) | 1:100 | R&D systems | AF6294 |
| SOX9 | Rabbit (polyclonal) | 1:500 | Millipore | AB5535 |
| TRPM1 | Rabbit (polyclonal) | 1:200 | ATLAS | HPA014779 |
| CHAT | Goat (polyclonal) | 1:300 | Merck Millipore | AB144P |
| mCAR | Rabbit (polyclonal) | 1:200 | Merck Millipore | AB15282 |

***Table S6:*** Primary antibody list

### Table S7

| Kinase Name | GeneID | Kinase Concentration (nM) | ATP Concentration (µM) | Substrate Name | Substrate Concentration (µg/50µl) |
|---|---|---|---|---|---|
| ABL1 | ABL1 | 1.3 | 0.3 | Poly(Ala,Glu,Lys,Tyr)6:2:5:1 | 0.125 |
| ABL2 | ABL2 | 1.6 | 0.3 | Poly(Ala,Glu,Lys,Tyr)6:2:5:1 | 0.125 |
| ACK1 | TNK2 | 14.1 | 1 | Poly(Glu,Tyr)4:1 | 0.125 |
| ACVR1 | ACVR1 | 14.0 | 3 | GSK3(14-27) | 2.0 |
| ACVR1B | ACVR1B | 8.8 | 3 | RBER-CHKtide | 2.0 |
| ACVR2A | ACVR2A | 15.2 | 3 | Casein | 1.0 |
| ACVR2B | ACVR2B | 22.6 | 1 | RB-CTF | 1.0 |
| ACVRL1 | ACVRL1 | 5.8 | 1 | Casein | 1.0 |
| AKT1 | AKT1 | 11.9 | 1 | GSK3(14-27) | 2.0 |
| AKT2 | AKT2 | 59.3 | 10 | GSK3(14-27) | 4.0 |
| AKT3 | AKT3 | 11.9 | 3 | GSK3(14-27) | 4.0 |
| ALK | ALK | 9.9 | 0.3 | Poly(Glu,Tyr)4:1 | 0.125 |
| AMPKalpha1 | PRKAA1 | 43.4 | 3 | RBER-CHKtide | 2.0 |
| ARAF YDYD | ARAF | 61.0 | 1 | MEK1 K97M (kinase-dead) | 0.5 |
| ARK5 | NUAK1 | 38.4 | 3 | RBER-CHKtide | 1.0 |
| ASK1 | MAP3K5 | 1.6 | 3 | GSK3(14-27) | 2.0 |
| AuroraA | AURKA | 18.3 | 10 | tetra(LRRWSLG) | 0.5 |
| AuroraB | AURKB | 28.5 | 10 | tetra(LRRWSLG) | 0.25 |
| AuroraC | AURKC | 68.4 | 3 | tetra(LRRWSLG) | 0.25 |
| AXL | AXL | 5.2 | 0.3 | Poly(Glu,Tyr)4:1 | 0.25 |
| BLK | PTK6 | 4.6 | 0.3 | Poly(Glu,Tyr)4:1 | 0.125 |
| BMPR1A | BMPR1A | 57.9 | 3 | Casein | 1.0 |
| BMPR1B | BMPR1B | 14.6 | 3 | Casein | 1.0 |
| BMX | BMX | 2.4 | 1 | Poly(Glu,Tyr)4:1 | 0.25 |
| BRAF | BRAF | 14.5 | 1 | MEK1 K97M (kinase-dead) | 0.5 |
| BRK | BRK | 12.2 | 3 | Poly(Glu,Tyr)4:1 | 0.125 |
| BRSK1 | BRSK1 | 2.7 | 1 | RBER-CHKtide | 2.0 |
| BRSK2 | BRSK2 | 9.7 | 1 | RBER-CHKtide | 2.0 |
| BTK | BTK | 28.6 | 3 | Poly(Glu,Tyr)4:1 | 0.25 |
| BUB1B | BUB1B | 9.5 | 0.3 | Casein | 0.25 |
| CAMK1D | CAMK1D | 21.3 | 0.3 | RBER-CHKtide | 2.0 |
| CAMK2A | CAMK2A | 1.8 | 3 | RBER-CHKtide | 1.0 |
| CAMK2B | CAMK2B | 16.5 | 3 | RBER-CHKtide | 2.0 |
| CAMK2D | CAMK2D | 0.3 | 1 | RBER-CHKtide | 1.0 |
| CAMK2G | CAMK2G | 1.0 | 1 | S6-Peptide | 1.0 |
| CAMK4 | CAMK4 | 12.6 | 1 | JUN | 0.5 |
| CAMKK1 | CAMKK1 | 13.4 | 3 | RB-CTF | 4.0 |
| CAMKK2 | CAMKK2 | 3.4 | 3 | GSK3(14-27) | 2.0 |
| CDC42BPA | CDC42BPA | 1.7 | 0.1 | S6-Peptide | 1.0 |
| CDC42BPB | CDC42BPB | 1.2 | 0.1 | S6-Peptide | 1.0 |
| CDC7/DBF4 | CDC7 | 2.5 | 0.1 | Histone H1 | 0.25 |
| CDK1/CycA2 | CDK1 | 2.2 | 0.3 | RBER-CHKtide | 1.0 |
| CDK1/CycB1 | CDK1 | 5.6 | 1 | RBER-CHKtide | 2.0 |
| CDK1/CycE1 | CDK1 | 7.4 | 1 | RBER-CHKtide | 2.0 |
| CDK10/CycQ | CDK10 | 15.8 | 3 | RBER-IRStide | 2.0 |
| CDK11B/CycK | CDK11B | 35.4 | 0.1 | SUPT5 754-837 | 2.0 |
| CDK12/CycK | CDK12 | 29.4 | 0.3 | RBER-IRStide | 2.0 |
| CDK13/CycK | CDK13 | 14.6 | 0.1 | SUPT5 754-837 | 0.5 |
| CDK14/CycY | CDK14 | 9.4 | 0.1 | RBER-IRStide | 4.0 |

| Kinase Name | GeneID | Kinase Concentration (nM) | ATP Concentration (µM) | Substrate Name | Substrate Concentration (µg/50µl) |
|---|---|---|---|---|---|
| CDK15/CycA2 | CDK15 | 37.1 | 1 | RBER-NTRK3tide | 4.0 |
| CDK15/CycB1 | CDK15 | 43.1 | 0.3 | RBER-NTRK3tide | 4.0 |
| CDK16/CycY | CDK16 | 6.4 | 0.3 | GSK3(14-27) | 2.0 |
| CDK17/p35NCK | CDK17 | 19.8 | 3 | RBER-CDC25tide | 1.0 |
| CDK18/CycY | CDK18 | 3.3 | 1 | RBER-NTRK3tide | 2.0 |
| CDK19/CycC | CDK19 | 32.7 | 3 | RBER-IRStide | 2.0 |
| CDK2/CycA2 | CDK2 | 3.7 | 0.3 | RBER-CHKtide | 1.0 |
| CDK2/CycD1 | CDK2 | 49.7 | 3 | RBER-CHKtide | 2.0 |
| CDK2/CycE1 | CDK2 | 1.1 | 1 | RBER-CHKtide | 1.0 |
| CDK20/CycH | CDK20 | 36.6 | 0.1 | RBER-CHKtide | 2.0 |
| CDK20/CycT1 | CDK20 | 26.4 | 1 | RBER-CHKtide | 4.0 |
| CDK3/CycC | CDK3 | 37.7 | 3 | RBER-CHKtide | 2.0 |
| CDK3/CycE1 | CDK3 | 2.1 | 3 | RBER-CHKtide | 1.0 |
| CDK4/CycD1 | CDK4 | 8.4 | 3 | RBER-CHKtide | 2.0 |
| CDK4/CycD2 | CDK4 | 2.8 | 3 | RBER-CHKtide | 1.0 |
| CDK4/CycD3 | CDK4 | 10.3 | 10 | RBER-CHKtide | 2.0 |
| CDK5/p25NCK | CDK5 | 3.3 | 0.3 | RBER-CHKtide | 1.0 |
| CDK5/p35NCK | CDK5 | 0.8 | 0.1 | RBER-CHKtide | 1.0 |
| CDK6/CycD1 | CDK6 | 3.2 | 3 | RBER-CHKtide | 2.0 |
| CDK6/CycD2 | CDK6 | 5.4 | 10 | RBER-CHKtide | 1.0 |
| CDK6/CycD3 | CDK6 | 18.1 | 30 | RBER-CHKtide | 2.0 |
| CDK7/CycH/MAT1 | CDK7 | 6.6 | 3 | RBER-CHKtide | 2.0 |
| CDK8/CycC | CDK8 | 16.5 | 1 | RBER-IRStide | 1.0 |
| CDK9/CycK | CDK9 | 3.7 | 1 | RBER-CHKtide | 2.0 |
| CDK9/CycT1 | CDK9 | 6.3 | 1 | RBER-CHKtide | 1.0 |
| CHK1 | CHEK1 | 11.9 | 1 | RBER-CHKtide | 2.0 |
| CHK2 | CHEK2 | 3.0 | 1 | tetra(LRRWSLG) | 0.5 |
| CIT 1-450 | CIT | 2.5 | 0.1 | Histone H2B | 2.0 |
| CK1alpha1 | CSNK1A1 | 14.0 | 0.3 | Casein | 1.0 |
| CK1delta | CSNK1D | 0.3 | 0.3 | Casein | 0.5 |
| CK1epsilon | CSNK1E | 0.8 | 0.3 | Casein | 0.5 |
| CK1gamma1 | CSNK1G1 | 0.3 | 0.1 | Casein | 1.0 |
| CK1gamma2 | CSNK1G2 | 0.4 | 0.1 | Casein | 1.0 |
| CK1gamma3 | CSNK1G3 | 0.5 | 0.1 | Casein | 1.0 |
| CK2alpha1 | CSNK2A1 | 1.4 | 0.1 | Casein | 1.0 |
| CK2alpha2 | CSNK2A2 | 2.8 | 0.1 | Casein | 1.0 |
| CLK1 | CLK1 | 46.1 | 0.3 | H2O (Autophos.) | 0 |
| CLK2 | CLK2 | 0.3 | 0.3 | GSK3(14-27) | 1.0 |
| CLK3 | CLK3 | 2.7 | 0.3 | S6-Peptide | 1.0 |
| CLK4 | CLK4 | 4.7 | 0.1 | Myelin Basic Protein | 1.0 |
| COT | MAP3K8 | 83.8 | 3 | RBER-CHKtide | 3.0 |
| CSF1R | CSF1R | 5.1 | 0.3 | Poly(Glu,Tyr)4:1 | 0.125 |
| CSK | CSK | 2.5 | 1 | Poly(Glu,Tyr)4:1 | 0.125 |
| DAPK1 | DAPK1 | 4.2 | 0.1 | GSK3(14-27) | 2.0 |
| DAPK2 | DAPK2 | 2.1 | 0.1 | S6-Peptide | 2.0 |
| DAPK3 | DAPK3 | 0.8 | 0.1 | GSK3(14-27) | 2.0 |
| DCAMKL2 | DCLK2 | 9.7 | 1 | RBER-CHKtide | 1.0 |
| DDR2 | DDR2 | 12.9 | 0.1 | Poly(Ala,Glu,Lys,Tyr)6:2:5:1 | 0.125 |
| DMPK | DMPK | 3.1 | 0.1 | tetra(LRRWSLG) | 2.0 |
| DNAPK | PRKDC | 0.3 | 1 | S6-Peptide | 2.0 |
| DYRK1A | DYRK1A | 0.9 | 1 | RBER-CHKtide | 2.0 |
| DYRK1B | DYRK1B | 1.0 | 0.3 | RBER-CHKtide | 2.0 |

| Kinase Name | GeneID | Kinase Concentration (nM) | ATP Concentration (µM) | Substrate Name | Substrate Concentration (µg/50µl) |
|---|---|---|---|---|---|
| DYRK2 | DYRK2 | 0.7 | 1 | RBER-IRStide | 2.0 |
| DYRK3 | DYRK3 | 0.6 | 0.3 | RBER-IRStide | 1.0 |
| DYRK4 | DYRK4 | 2.3 | 0.1 | RBER-CHKtide | 1.0 |
| EEF2K | EEF2K | 0.4 | 0.3 | GSK3(14-27) | 2.0 |
| EGFR | EGFR | 2.2 | 0.3 | Poly(Glu,Tyr)4:1 | 0.125 |
| EIF2AK2 | EIF2AK2 | 2.3 | 0.3 | RB-CTF | 1.0 |
| EIF2AK3 | EIF2AK3 | 2.1 | 0.3 | Casein | 2.0 |
| EIF2AK4 | EIF2AK4 | 4.7 | 3 | S6-Peptide | 0.5 |
| EPHA1 | EPHA1 | 12.6 | 1 | Poly(Glu,Tyr)4:1 | 0.125 |
| EPHA2 | EPHA2 | 5.4 | 3 | Poly(Glu,Tyr)4:1 | 0.125 |
| EPHA3 | EPHA3 | 13.2 | 3 | Poly(Glu,Tyr)4:1 | 0.25 |
| EPHA4 | EPHA4 | 65.3 | 10 | Poly(Glu,Tyr)4:1 | 0.25 |
| EPHA5 | EPHA5 | 5.0 | 1 | Poly(Glu,Tyr)4:1 | 0.5 |
| EPHA6 | EPHA6 | 25.4 | 3 | Poly(Glu,Tyr)4:1 | 0.125 |
| EPHA7 | EPHA7 | 12.7 | 1 | Poly(Glu,Tyr)4:1 | 0.125 |
| EPHA8 | EPHA8 | 5.2 | 0.3 | Poly(Glu,Tyr)4:1 | 0.125 |
| EPHB1 | EPHB1 | 2.6 | 1 | Poly(Glu,Tyr)4:1 | 0.125 |
| EPHB2 | EPHB2 | 12.6 | 0.3 | Poly(Ala,Glu,Lys,Tyr)6:2:5:1 | 0.125 |
| EPHB3 | EPHB3 | 6.2 | 1 | Poly(Glu,Tyr)4:1 | 0.125 |
| EPHB4 | EPHB4 | 13.0 | 1 | Poly(Glu,Tyr)4:1 | 0.125 |
| ERBB2 | ERBB2 | 10.6 | 1 | Poly(Glu,Tyr)4:1 | 0.125 |
| ERBB4 | ERBB4 | 1.0 | 0.3 | Poly(Glu,Tyr)4:1 | 0.125 |
| ERK1 | MAPK3 | 7.2 | 1 | RBER-CHKtide | 2.0 |
| ERK2 | MAPK1 | 4.8 | 0.3 | RBER-CHKtide | 2.0 |
| ERK5 | MAPK7 | 79.3 | 1 | RBER-CHKtide | 2.0 |
| ERK7 | MAPK15 | 32.5 | 1 | RBER-CHKtide | 2.0 |
| FAK | PTK2 | 6.9 | 1 | Poly(Glu,Tyr)4:1 | 0.125 |
| FER | FER | 3.2 | 1 | Poly(Glu,Tyr)4:1 | 0.125 |
| FES | FES | 0.6 | 0.3 | Poly(Glu,Tyr)4:1 | 0.125 |
| FGFR1 | FGFR1 | 13.0 | 3 | Poly(Glu,Tyr)4:1 | 0.125 |
| FGFR2 | FGFR2 | 12.6 | 1 | Poly(Glu,Tyr)4:1 | 0.25 |
| FGFR3 | FGFR3 | 13.5 | 3 | Poly(Glu,Tyr)4:1 | 0.25 |
| FGFR4 | FGFR4 | 13.6 | 1 | Poly(Glu,Tyr)4:1 | 0.125 |
| FGR | FGR | 5.3 | 0.3 | Poly(Glu,Tyr)4:1 | 0.125 |
| FLT3 | FLT3 | 13.0 | 1 | Poly(Ala,Glu,Lys,Tyr)6:2:5:1 | 0.125 |
| FRK | FRK | 4.6 | 0.3 | Poly(Glu,Tyr)4:1 | 0.5 |
| FYN | FYN | 2.2 | 0.3 | Poly(Glu,Tyr)4:1 | 0.125 |
| GRK2 | ADRBK1 | 8.5 | 1 | Casein | 0.5 |
| GRK3 | ADRBK2 | 1.9 | 3 | Casein | 2.0 |
| GRK4 | GRK4 | 1.1 | 1 | Casein | 0.5 |
| GRK5 | GRK5 | 0.2 | 0.3 | Casein | 1.0 |
| GRK6 | GRK6 | 1.1 | 1 | Casein | 1.0 |
| GRK7 | GRK7 | 2.2 | 0.3 | Casein | 1.0 |
| GSG2 | HASPIN | 2.5 | 0.1 | Histone H3 (1-34) | 0.5 |
| GSK3alpha | GSK3A | 3.7 | 0.1 | SUPT5 754-837 | 1.0 |
| GSK3beta | GSK3B | 19.7 | 0.3 | RBER-CHKtide | 1.0 |
| HCK | HCK | 4.6 | 0.3 | Poly(Glu,Tyr)4:1 | 0.125 |
| HIPK1 | HIPK1 | 13.6 | 0.3 | RBER-CHKtide | 2.0 |
| HIPK2 | HIPK2 | 8.2 | 0.1 | RBER-CHKtide | 2.0 |
| HIPK3 | HIPK3 | 21.7 | 0.3 | RBER-CHKtide | 2.0 |
| HIPK4 | HIPK4 | 1.0 | 0.1 | RBER-IRStide | 1.0 |
| HRI | EIF2AK1 | 9.9 | 0.3 | Casein | 0.25 |

| Kinase Name | GeneID | Kinase Concentration (nM) | ATP Concentration (µM) | Substrate Name | Substrate Concentration (µg/50µl) |
|---|---|---|---|---|---|
| IGF1R | IGF1R | 7.8 | 1 | Poly(Glu,Tyr)4:1 | 0.125 |
| IKKalpha | CHUK | 7.9 | 0.3 | RBER-CHKtide | 2.0 |
| IKKbeta | IKBKB | 8.3 | 0.3 | RBER-CHKtide | 1.0 |
| IKKepsilon | IKBKE | 3.6 | 1 | GSK3(14-27) | 1.0 |
| INSR | INSR | 2.8 | 3 | Poly(Ala,Glu,Lys,Tyr)6:2:5:1 | 0.125 |
| INSRR | INSRR | 7.5 | 3 | Poly(Ala,Glu,Lys,Tyr)6:2:5:1 | 0.25 |
| IRAK1 | IRAK1 | 1.2 | 1 | RB-CTF | 2.0 |
| IRAK4 | IRAK4 | 7.5 | 3 | RB-CTF | 1.0 |
| ITK | ITK | 11.8 | 1 | Poly(Glu,Tyr)4:1 | 0.125 |
| JAK1 | JAK1 | 42.7 | 1 | RBER-IRStide | 2.0 |
| JAK2 | JAK2 | 6.4 | 0.3 | Poly(Ala,Glu,Lys,Tyr)6:2:5:1 | 0.125 |
| JAK3 | JAK3 | 2.9 | 0.1 | Poly(Ala,Glu,Lys,Tyr)6:2:5:1 | 0.125 |
| JNK1 | MAPK8 | 2.3 | 0.3 | ATF2 | 0.25 |
| JNK2 | MAPK9 | 2.0 | 1 | ATF2 | 1.0 |
| JNK3 | MAPK10 | 2.1 | 0.3 | ATF2 | 2.0 |
| KIT | KIT | 38.7 | 3 | Poly(Glu,Tyr)4:1 | 0.125 |
| LCK | LCK | 4.3 | 0.3 | Poly(Glu,Tyr)4:1 | 0.125 |
| LIMK1 | LIMK1 | 1.5 | 0.1 | Cofilin2 | 4.0 |
| LIMK2 | LIMK2 | 13.1 | 0.1 | Cofilin1 | 4.0 |
| LKB1/MO25a/STRADa | STK11 | 6.7 | 0.3 | RB-CTF | 2.0 |
| LRRK2 | LRRK2 | 2.9 | 0.3 | GSK3(14-27) | 2.0 |
| LTK | LTK | 5.4 | 0.3 | Poly(Glu,Tyr)4:1 | 0.125 |
| LYN | LYN | 1.1 | 0.3 | Poly(Glu,Tyr)4:1 | 0.125 |
| MAP3K1 | MAP3K1 | 2.4 | 0.1 | tetra(LRRWSLG) | 1.0 |
| MAP3K10 | MAP3K10 | 2.3 | 1 | Histone H2B | 0.5 |
| MAP3K11 | MAP3K11 | 13.3 | 3 | Myelin Basic Protein | 1.0 |
| MAP3K7/MAP3K7IP1 | MAP3K7 | 13.1 | 1 | S6-Peptide | 2.0 |
| MAP3K9 | MAP3K9 | 3.6 | 3 | GSK3(14-27) | 1.0 |
| MAP4K1 | MAP4K1 | 7.5 | 0.3 | Casein | 1.0 |
| MAP4K2 | MAP4K2 | 1.0 | 1 | PKC(19-31) | 0.5 |
| MAP4K4 | MAP4K4 | 16.8 | 1 | GSK3(14-27) | 2.0 |
| MAP4K5 | MAP4K5 | 0.5 | 0.3 | Casein | 1.0 |
| MAPKAPK2 | MAPKAPK2 | 3.8 | 0.1 | RBER-CHKtide | 1.0 |
| MAPKAPK3 | MAPKAPK3 | 4.6 | 0.1 | tetra(LRRWSLG) | 0.25 |
| MAPKAPK5 | MAPKAPK5 | 1.8 | 0.1 | RBER-CHKtide | 2.0 |
| MARK1 | MARK1 | 17.0 | 1 | RBER-CHKtide | 1.0 |
| MARK2 | MARK2 | 1.8 | 1 | RBER-CHKtide | 2.0 |
| MARK3 | MARK3 | 18.3 | 1 | RBER-CHKtide | 1.0 |
| MARK4 | MARK4 | 2.9 | 3 | RBER-CHKtide | 2.0 |
| MASTL | MASTL | 47.5 | 3 | RBER-NTRK3tide | 4.0 |
| MATK | MATK | 1.9 | 1 | Poly(Glu,Tyr)4:1 | 0.125 |
| MEK1 | MAP2K1 | 34.4 | 3 | ERK2 K54R (kinase-dead) | 2.0 |
| MEK2 | MAP2K2 | 20.2 | 1 | RBER-IRStide | 1.0 |
| MEK5 | MAP2K5 | 37.0 | 1 | RBER-CHKtide | 2.0 |
| MEKK2 | MAP3K2 | 19.9 | 0.3 | Casein | 0.5 |
| MEKK3 | MAP3K3 | 19.7 | 0.3 | Casein | 0.5 |
| MELK | MELK | 50.2 | 0.1 | RBER-CHKtide | 2.0 |
| MERTK | MERTK | 24.5 | 1 | Poly(Glu,Tyr)4:1 | 0.25 |
| MET | MET | 2.5 | 0.3 | Poly(Ala,Glu,Lys,Tyr)6:2:5:1 | 0.125 |
| MINK1 | MINK1 | 6.0 | 1 | RBER-CHKtide | 1.0 |
| MKK3 | MAP2K3 | 115.1 | 0.3 | p38-alpha K53A (kinase-dead) | 1.0 |
| MKK4 | MAP2K4 | 2.0 | 0.1 | JNK1 K55R K56R (kinase-dead) | 1.0 |

| Kinase Name | GeneID | Kinase Concentration (nM) | ATP Concentration (µM) | Substrate Name | Substrate Concentration (µg/50µl) |
|---|---|---|---|---|---|
| MKK6 SDTD | MAP2K6 | 5.9 | 1 | p38-alpha K53A (kinase-dead) | 1.0 |
| MKK7 | MAP2K7 | 56.7 | 3 | JNK1 K55R K56R (kinase-dead) | 2.0 |
| MKNK1 | MKNK1 | 15.0 | 1 | S6-Peptide | 2.0 |
| MKNK2 | MKNK2 | 4.3 | 0.1 | S6-Peptide | 1.0 |
| MLK4 | MAP3K21 | 58.7 | 3 | RB-CTF | 2.0 |
| MST1 | STK4 | 1.2 | 0.3 | RB-CTF | 2.0 |
| MST2 | STK3 | 5.3 | 1 | RB-CTF | 1.0 |
| MST3 | STK24 | 12.7 | 1 | Casein | 1.0 |
| MST4 | STK26 | 13.1 | 1 | RB-CTF | 1.0 |
| MTOR | MTOR | 2.4 | 1 | Casein | 0.5 |
| MUSK | MUSK | 43.9 | 1 | Poly(Ala,Glu,Lys,Tyr)6:2:5:1 | 0.125 |
| MYLK | MYLK | 10.4 | 0.1 | S6-Peptide | 1.0 |
| MYLK2 | MYLK2 | 3.2 | 1 | S6-Peptide | 1.0 |
| MYLK3 | MYLK3 | 5.3 | 0.1 | S6-Peptide | 2.0 |
| NDR1 | STK38 | 48.6 | 0.1 | GSK3(14-27) | 2.0 |
| NDR2 | STK38L | 48.6 | 0.1 | GSK3(14-27) | 4.0 |
| NEK1 | NEK1 | 7.0 | 3 | GSK3(14-27) | 2.0 |
| NEK11 | NEK11 | 5.9 | 1 | Myelin Basic Protein | 1.0 |
| NEK2 | NEK2 | 24.2 | 3 | RB-CTF | 1.0 |
| NEK3 | NEK3 | 46.5 | 3 | RBER-CHKtide | 4.0 |
| NEK4 | NEK4 | 0.5 | 0.3 | Myelin Basic Protein | 2.0 |
| NEK6 | NEK6 | 14.7 | 1 | GSK3(14-27) | 2.0 |
| NEK7 | NEK7 | 4.6 | 1 | Casein | 1.0 |
| NEK9 | NEK9 | 0.6 | 10 | GSK3(14-27) | 1.0 |
| NIK | MAP3K14 | 52.5 | 1 | RBER-CHKtide | 2.0 |
| NLK | NLK | 6.5 | 1 | GSK3(14-27) | 2.0 |
| p38alpha | MAPK14 | 7.2 | 1 | ATF2 | 1.0 |
| p38beta | MAPK11 | 1.4 | 1 | ATF2 | 1.0 |
| p38delta | MAPK13 | 0.9 | 1 | RBER-CHKtide | 2.0 |
| p38gamma | MAPK12 | 4.7 | 0.1 | RBER-IRStide | 2.0 |
| PAK1 | PAK1 | 2.2 | 10 | tetra(LRRWSLG) | 1.0 |
| PAK2 | PAK2 | 4.5 | 1 | tetra(LRRWSLG) | 0.25 |
| PAK3 | PAK3 | 3.3 | 3 | tetra(LRRWSLG) | 0.5 |
| PAK4 | PAK4 | 10.6 | 3 | tetra(LRRWSLG) | 0.5 |
| PAK6 | PAK6 | 2.9 | 0.3 | tetra(LRRWSLG) | 0.125 |
| PAK7 | PAK7 | 2.7 | 1 | tetra(LRRWSLG) | 0.25 |
| PASK | PASK | 6.5 | 0.1 | RBER-CHKtide | 2.0 |
| PBK | PBK | 59.2 | 3 | Histone H1 | 0.5 |
| PDGFRalpha | PDGFRA | 22.2 | 3 | Poly(Ala,Glu,Lys,Tyr)6:2:5:1 | 0.125 |
| PDGFRbeta | PDGFRB | 2.3 | 0.3 | Poly(Ala,Glu,Lys,Tyr)6:2:5:1 | 0.125 |
| PDK1 | PDPK1 | 1.8 | 0.3 | tetra(LRRWSLG) | 0.25 |
| PHKG1 | PHKG1 | 2.8 | 3 | PKC(19-31) | 0.5 |
| PHKG2 | PHKG2 | 2.7 | 0.1 | RBER-CHKtide | 2.0 |
| PIM1 | PIM1 | 0.6 | 0.1 | GSK3(14-27) | 1.0 |
| PIM2 | PIM2 | 15.6 | 0.3 | GSK3(14-27) | 1.0 |
| PIM3 | PIM3 | 7.8 | 1 | GSK3(14-27) | 2.0 |
| PKA | PRKACA | 4.6 | 1 | tetra(LRRWSLG) | 0.5 |
| PKCalpha | PRKCA | 0.9 | 10 | PKC(19-31) | 0.25 |
| PKCbeta1 | PRKCB | 1.0 | 10 | PKC(19-31) | 0.25 |
| PKCbeta2 | PRKCB | 3.9 | 10 | PKC(19-31) | 0.25 |
| PKCdelta | PRKCD | 4.5 | 10 | PKC(19-31) | 0.5 |
| PKCepsilon | PRKCE | 1.8 | 1 | PKC(19-31) | 0.125 |

| Kinase Name | GeneID | Kinase Concentration (nM) | ATP Concentration (µM) | Substrate Name | Substrate Concentration (µg/50µl) |
|---|---|---|---|---|---|
| PKCeta | PRKCH | 3.7 | 3 | Histone H2B | 0.25 |
| PKCgamma | PRKCG | 3.6 | 10 | PKC(19-31) | 0.25 |
| PKCiota | PRKCI | 10.2 | 10 | PKC(19-31) | 0.5 |
| PKCmu | PRKD1 | 3.6 | 0.3 | RBER-CHKtide | 1.0 |
| PKCnu | PRKCQ | 4.0 | 3 | tetra(LRRWSLG) | 0.25 |
| PKCtheta | PRKCQ | 1.8 | 3 | PKC(19-31) | 0.25 |
| PKCzeta | PRKCZ | 10.3 | 1 | PKC(19-31) | 0.125 |
| PKMzeta | PRKCL1 | 2.3 | 1 | PKC(19-31) | 0.25 |
| PKN3 | PKN3 | 7.9 | 1 | PKC(19-31) | 1.0 |
| PLK1 | PLK1 | 5.0 | 1 | RBER-CHKtide | 2.0 |
| PLK3 | PLK3 | 6.2 | 0.3 | Casein | 0.5 |
| PRK1 | PKN1 | 7.3 | 0.3 | RBER-CHKtide | 2.0 |
| PRK2 | PKN2 | 1.4 | 0.1 | RBER-CHKtide | 2.0 |
| PRKD2 | PRKD2 | 0.8 | 0.3 | RBER-CHKtide | 2.0 |
| PRKG1 | PRKG1 | 0.9 | 0.3 | PKC(19-31) | 1.0 |
| PRKG2 | PRKG2 | 0.3 | 1 | GSK3(14-27) | 2.0 |
| PRKX | PRKX | 5.8 | 0.3 | GSK3(14-27) | 1.0 |
| PYK2 | PTK2B | 10.3 | 1 | Poly(Glu,Tyr)4:1 | 0.125 |
| RAF1 YDYD | RAF1 | 2.6 | 0.3 | MEK1 K97M (kinase-dead) | 0.5 |
| RET | RET | 24.9 | 1 | Poly(Glu,Tyr)4:1 | 0.125 |
| RIPK2 | RIPK2 | 78.4 | 3 | RBER-CHKtide | 4.0 |
| RIPK4 | RIPK4 | 31.6 | 0.3 | Casein | 0.5 |
| RIPK5 | RIPK5 | 1.5 | 1 | RBER-CHKtide | 2.0 |
| ROCK1 | ROCK1 | 1.1 | 0.3 | PKC(19-31) | 1.0 |
| ROCK2 | ROCK2 | 1.1 | 0.3 | S6-Peptide | 1.0 |
| RON | MST1R | 15.4 | 1 | Poly(Glu,Tyr)4:1 | 0.125 |
| ROS | ROS1 | 0.5 | 0.3 | Poly(Ala,Glu,Lys,Tyr)6:2:5:1 | 0.125 |
| RPS6KA1 | RPS6KA1 | 6.8 | 1 | RBER-CHKtide | 2.0 |
| RPS6KA2 | RPS6KA2 | 1.1 | 1 | tetra(LRRWSLG) | 0.5 |
| RPS6KA3 | RPS6KA3 | 0.2 | 1 | tetra(LRRWSLG) | 0.25 |
| RPS6KA4 | RPS6KA4 | 8.8 | 0.3 | RBER-CHKtide | 2.0 |
| RPS6KA5 | RPS6KA5 | 12.6 | 1 | RBER-CHKtide | 2.0 |
| RPS6KA6 | RPS6KA6 | 0.9 | 1 | GSK3(14-27) | 2.0 |
| S6K | RPS6KB1 | 13.5 | 3 | GSK3(14-27) | 2.0 |
| S6Kbeta | RPS6KB2 | 24.3 | 1 | RBER-CHKtide | 4.0 |
| SAK | PLK4 | 28.9 | 1 | p38-alpha K53A (kinase-dead) | 2.0 |
| SGK1 | SGK1 | 53.4 | 1 | GSK3(14-27) | 1.0 |
| SGK2 | SGK2 | 5.8 | 1 | GSK3(14-27) | 1.0 |
| SGK3 | SGK3 | 12.0 | 1 | GSK3(14-27) | 2.0 |
| SIK1 | SIK1 | 29.2 | 3 | RBER-CHKtide | 2.0 |
| SIK2 | SIK2 | 3.3 | 1 | RBER-CHKtide | 2.0 |
| SIK3 | SIK3 | 15.9 | 1 | RBER-CHKtide | 2.0 |
| SLK | STK10 | 3.7 | 1 | S6-Peptide | 2.0 |
| SNARK | NUAK1 | 20.2 | 0.3 | Histone H2B | 0.5 |
| SNK | NUAK2 | 9.3 | 1 | GSK3(14-27) | 2.0 |
| SRC | SRC | 2.2 | 0.3 | Poly(Glu,Tyr)4:1 | 0.125 |
| SRMS | SRMS | 9.8 | 3 | Poly(Glu,Tyr)4:1 | 0.125 |
| SRPK1 | SRPK1 | 5.8 | 0.3 | Myelin Basic Protein | 1.0 |
| SRPK2 | SRPK2 | 3.7 | 0.3 | Myelin Basic Protein | 1.0 |
| STK17A | STK17A | 5.4 | 0.1 | RB-CTF | 2.0 |
| STK17B | STK17B | 14.2 | 0.1 | MRCL3 | 2.0 |
| STK23 | MST1 | 3.5 | 0.3 | RBER-CHKtide | 2.0 |

| Kinase Name | GeneID | Kinase Concentration (nM) | ATP Concentration (μM) | Substrate Name | Substrate Concentration (μg/50μl) |
|---|---|---|---|---|---|
| STK25 | STK25 | 2.6 | 0.3 | Casein | 1.0 |
| STK33 | STK33 | 16.2 | 1 | RBER-CHKtide | 2.0 |
| STK39 | STK39 | 13.2 | 1 | RBER-CDC25tide | 1.0 |
| SYK | SYK | 9.6 | 1 | Poly(Glu,Tyr)4:1 | 0.125 |
| TAOK2 | TAOK2 | 4.8 | 0.3 | Casein | 1.0 |
| TAOK3 | TAOK3 | 18.9 | 1 | PKC(19-31) | 0.5 |
| TBK1 | TBK1 | 1.2 | 1 | Casein | 1.0 |
| TEC | TEC | 39.3 | 10 | Poly(Glu,Tyr)4:1 | 0.125 |
| TGFBR1 | TGFBR1 | 6.2 | 1 | GSK3(14-27) | 1.0 |
| TGFBR2 | TGFBR2 | 2.9 | 0.1 | S6-Peptide | 2.0 |
| TIE2 | TEK | 14.5 | 1 | Poly(Glu,Tyr)4:1 | 0.125 |
| TLK1 | TLK1 | 6.3 | 0.3 | RB-CTF | 1.0 |
| TLK2 | TLK2 | 2.1 | 0.1 | S6-Peptide | 1.0 |
| TNK1 | TNK1 | 33.7 | 0.3 | Poly(Glu,Tyr)4:1 | 0.125 |
| TRKA | NTRK1 | 5.7 | 0.3 | Poly(Glu,Tyr)4:1 | 0.25 |
| TRKB | NTRK2 | 4.5 | 0.3 | Poly(Glu,Tyr)4:1 | 0.25 |
| TRKC | NTRK3 | 6.2 | 0.3 | Poly(Glu,Tyr)4:1 | 0.125 |
| TSF1 | DYRK3 | 7.8 | 0.1 | Casein | 1.0 |
| TSK2 | DYRK4 | 27.1 | 3 | RBER-CHKtide | 2.0 |
| TSSK1 | TSSK1 | 2.1 | 0.3 | RBER-CHKtide | 1.0 |
| TTBK1 | TTBK1 | 2.3 | 0.1 | RB-CTF | 1.0 |
| TTBK2 | TTBK2 | 2.4 | 0.1 | Casein | 0.5 |
| TTK | TTK | 15.8 | 0.3 | RBER-CHKtide | 1.0 |
| TXK | TXK | 1.7 | 1 | RBER-CHKtide | 2.0 |
| TYK2 | TYK2 | 2.9 | 0.1 | Poly(Ala,Glu,Lys,Tyr)6:2:5:1 | 0.125 |
| TYRO3 | TYRO3 | 5.1 | 1 | Poly(Glu,Tyr)4:1 | 0.5 |
| ULK1 | ULK1 | 8.4 | 0.3 | Casein | 1 |
| ULK2 | ULK2 | 3.2 | 0.3 | Casein | 0.5 |
| ULK3 | ULK3 | 4.9 | 0.3 | Casein | 1 |
| VEGFR1 | FLT1 | 11.2 | 1 | Poly(Glu,Tyr)4:1 | 0.125 |
| VEGFR2 | KDR | 11.5 | 1 | Poly(Glu,Tyr)4:1 | 0.125 |
| VEGFR3 | FLT4 | 11.7 | 3 | Poly(Glu,Tyr)4:1 | 0.125 |
| VRK1 | VRK1 | 53.2 | 3 | RBER-CHKtide | 2.0 |
| VRK2 | VRK2 | 48.1 | 1 | Casein | 0.5 |
| WEE1 | WEE1 | 53.4 | 1 | Poly(Ala,Glu,Lys,Tyr)6:2:5:1 | 0.125 |
| WNK1 | WNK1 | 127.6 | 30 | PKC(19-31) | 0.5 |
| WNK2 | WNK2 | 30.8 | 3 | RBER-CHKtide | 2.0 |
| WNK3 | WNK3 | 11.2 | 3 | S6-Peptide | 2.0 |
| YES | YES1 | 4.4 | 1 | Poly(Glu,Tyr)4:1 | 0.125 |
| ZAK | MAP3K20 | 1.2 | 1 | Myelin Basic Protein | 1.0 |
| ZAP70 | ZAP70 | 2.0 | 0.1 | Poly(Glu,Tyr)4:1 | 0.125 |

***Table S7:*** Kinase profiling assay conditions.
